# The role of population structure in computations through neural dynamics

**DOI:** 10.1101/2020.07.03.185942

**Authors:** Alexis Dubreuil, Adrian Valente, Manuel Beiran, Francesca Mastrogiuseppe, Srdjan Ostojic

**Affiliations:** Laboratoire de Neurosciences Cognitives et Computationnelles, INSERM U960, Ecole Normale Superieure - PSL Research University, 75005 Paris, France; Gatsby Computational Neuroscience Unit, UCL, London, Great Britain

## Abstract

Neural computations are currently investigated using two separate approaches: sorting neurons into functional populations, or examining the low-dimensional dynamics of collective activity. Whether and how these two aspects interact to shape computations is currently unclear. Using a novel approach to extract computational mechanisms from networks trained on neuroscience tasks, here we show that the dimensionality of the dynamics and cell-class structure play fundamentally complementary roles. While various tasks can be implemented by increasing the dimensionality in networks with fully random population structure, flexible input-output mappings instead required a non-random population structure that can be described in terms of multiple sub-populations. Our analyses revealed that such a population structure enabled flexible computations through a mechanism based on gain-controlled modulations that flexibly shape the dynamical landscape of collective dynamics. Our results lead to task-specific predictions for the structure of neural selectivity, inactivation experiments, and for the implication of different neurons in multi-tasking.

## 1 Introduction

The quest to understand the neural bases of cognition currently relies on two disjoint paradigms [Barack and Krakauer, 2021]. Classical works have sought to determine the computational role of individual cells by sorting them into functional populations based on their responses to sensory and behavioral variables [Hubel and Wiesel, 1959; Moser et al., 2017; Hardcastle et al., 2017]. Fast developing tools for dissecting neural circuits have opened the possibility of mapping such functional populations onto genetic and anatomic cell types, and given a new momentum to this cell-category approach [Adesnik et al., 2012; Ye et al., 2016; Kvitsiani et al., 2013; Hangya et al., 2014; Pinto and Dan, 2015; Hirokawa et al., 2019]. This viewpoint has however been challenged by observations that individual neurons often represent seemingly random mixtures of sensory and behavioral variables, especially in higher cortical areas [Churchland and Shenoy, 2007; Machens et al., 2010; Rigotti et al., 2013; Mante et al., 2013; Park et al., 2014], where sharply defined functional cell populations are often not directly apparent [Mante et al., 2013; Raposo et al., 2014; Hardcastle et al., 2017]. A newly emerging paradigm has therefore proposed that neural computations need instead to be interpreted in terms of collective dynamics in the state space of joint activity of all neurons [Buonomano and Maass, 2009; Rigotti et al., 2013; Mante et al., 2013; Gallego et al., 2017; Remington et al., 2018; Saxena and Cunningham, 2019]. This computation-through-dynamics framework [Vyas et al., 2020] hence posits that neural computations are revealed by studying the geometry of low-dimensional trajectories of activity in state space [Mante et al., 2013; Rajan et al., 2016; Chaisangmongkon et al., 2017; Remington et al., 2018; Wang et al., 2018; Sohn et al., 2019], while remaining agnostic to the role of any underlying population structure.

In view of the apparent antagonism between these two approaches, two works have sought to precisely assess the presence of functional cell populations in the posterior parietal cortex (PPC) [Raposo et al., 2014] and prefrontal cortex [Hirokawa et al., 2019]. Rather than define cell populations by classical methods such as thresholding the activity or selectivity of individual neurons, these studies developed new statistical techniques to determine whether the distribution of selectivity across neurons displayed non-random population structure [Hardcastle et al., 2017]. Using analogous analyses, but different behavioral tasks, the two studies reached opposite conclusions. Raposo et al found no evidence for non-random population structure in selectivity, and argued that PPC neurons fully multiplex information. Hirokawa et al also observed that individual neurons responded to mixtures of task features, but in contrast to Raposo et al, they detected important deviations from a fully random distribution of selectivity, a situation they termed non-random mixed selectivity. By clustering neurons according to their response properties, they defined separate, though mixed-selective populations that appeared to represent distinct task variables and to reflect underlying connectivity. To resolve the apparent discrepancy with Raposo et al, Hirokawa et al conjectured that revealing non-random population structure in higher cortical areas may require sufficiently complex behavioral tasks.

The conflicting findings of [Raposo et al., 2014; Hirokawa et al., 2019] therefore raise a fundamental theoretical question: do specific computational tasks require a non-random population structure, or alternatively can any task in principle be implemented with a fully random population structure as in Raposo et al. [2014]? To address this question, we trained recurrent neural networks on a range of systems neuroscience tasks [Sussillo, 2014; Barak, 2017; Yang et al., 2019] and examined the population structure that emerges in both selectivity and connectivity using identical methods as Raposo et al. [2014]; Hirokawa et al. [2019]. Starting from the premise that computations are necessarily determined by the underlying connectivity [Mastrogiuseppe and Ostojic, 2018], we then developed a new approach for assessing the computational role of population structure in connectivity for each task. Together, these analyses revealed that, while a fully random population structure was sufficient to implement a range of tasks, specific tasks appeared to require a non-random population structure in connectivity that could be described in terms of a small number of statistically-defined sub-populations. This was in particular the case when a flexible reconfiguration of input-output associations was needed, a common component of many cognitive tasks [Sakai, 2008] and more generally of multi-tasking [Yang et al., 2019; Duncker et al., 2020; Masse et al., 2018]. To extract the mechanistic role of this population structure for computations-through-dynamics, we focused on the class of low-rank models [Mastrogiuseppe and Ostojic, 2018; Schuessler et al., 2020a,b] that can be reduced to interpretable latent dynamics characterized by a minimal intrinsic dimension and number of sub-populations [Beiran et al., 2021]. We found that the subpopulation structure of the connectivity enables networks to implement flexible computations through a mechanism based on modulations of gain and effective interactions that flexibly modify the low-dimensional latent dynamics across epochs of the task. Specifically, at the level of the collective dynamics, the sub-population structure allows different inputs to act either as drivers or modulators [Sherman and Guillery, 1998; Salinas, 2004; Ferguson and Cardin, 2020]. Our results lead to task-specific predictions for the statistical structure of single-neuron selectivity, for inactivations of specific sub-populations, as well as for the implication of different neurons in multi-tasking.

## 2 Results

### 2.1 Identifying non-random population structure in trained recurrent networks

We trained recurrent neural networks (RNNs) on five systems neuroscience tasks spanning a range of cognitive components: perceptual decision-making (DM) [Gold and Shadlen, 2007], parametric working-memory (WM) [Romo et al., 1999], multi-sensory decision-making (MDM) [Raposo et al., 2014], contextual decision-making (CDM) [Mante et al., 2013] and delay-match-to-sample (DMS) [Miyashita, 1988]. Each network consisted of *N* units, and the activation *x*_*i*_ of unit *i* was given by

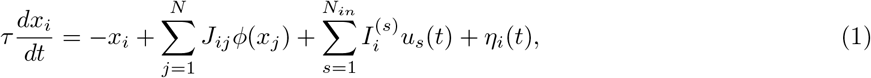

where *ϕ*(*x*) = tanh(*x*) is the single-unit non-linearity, *J*_*ij*_ is the recurrent connectivity matrix, *η*_*i*_(*t*) is a single-unit noise and the network receives *N*_*in*_ task-defined inputs 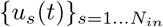 through a set of feed-forward weights 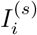 (see Methods 4.1). The output *z*(*t*) of the network was obtained by a linear readout of firing rates *ϕ*(*x*_*i*_) through a set of weights {*w*_*i*_}_*i*=1…*N*_ (Fig. 1a top). Each task was modeled as a mapping from a set of inputs representing stimuli and contextual cues to desired outputs (see Methods 4.3). For each task, we used gradient-descent to train 100 networks starting from different, random initial connectivities [Yang and Wang, 2020]. We then searched for evidence of non-random population structure by comparing the selectivity, connectivity and performance of the trained networks with randomized shuffles.

**Figure 1:**
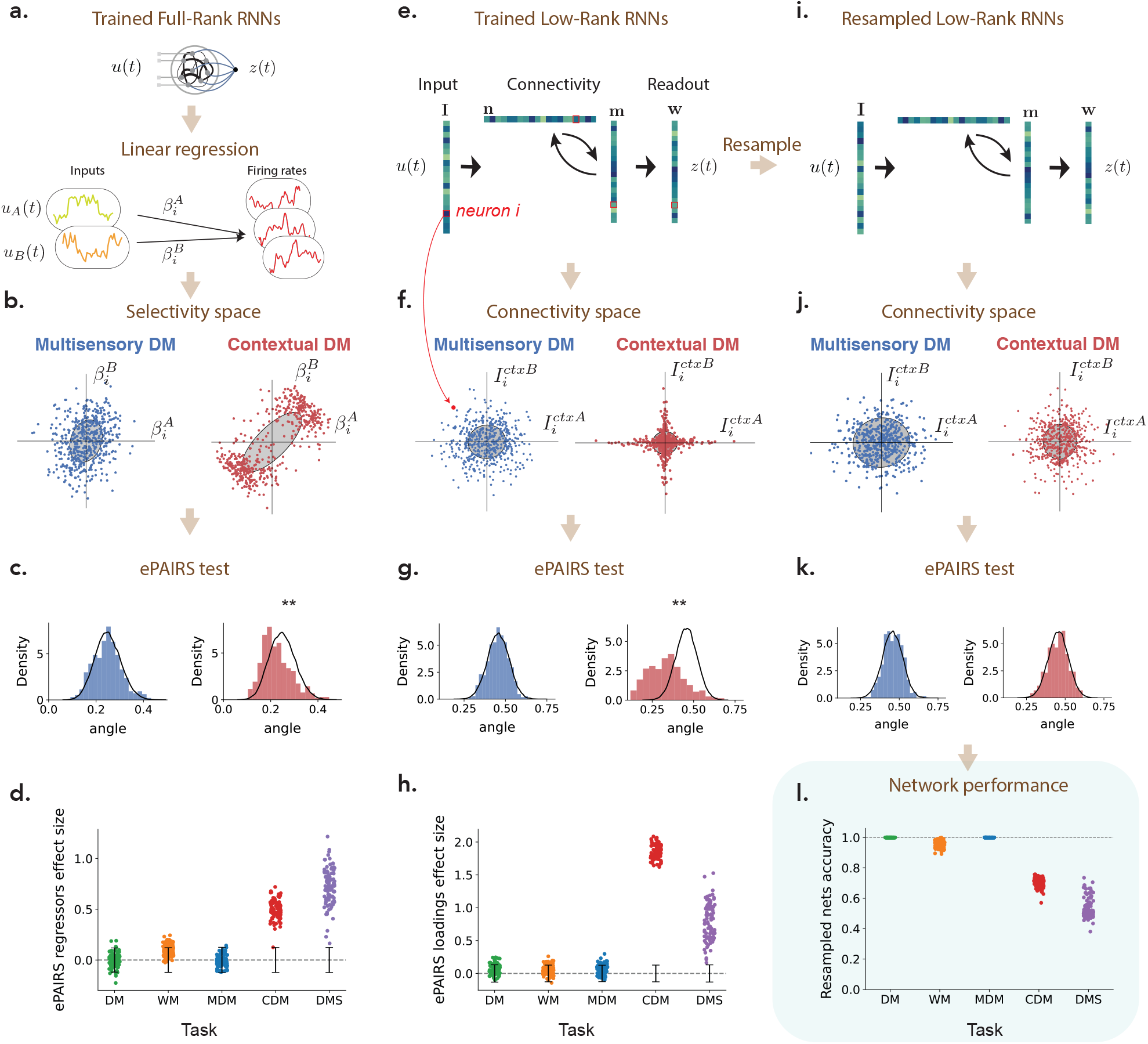
Identifying non-random population structure in selectivity, connectivity and computations. (a) Recurrent neural networks (RNNs) were trained separately on five tasks. For each task, and each trained RNN, selectivity was first quantified by computing linear regression coefficients 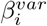 for each neuron *i* with respect to task-defined variables such as stimulus features or decision (see Methods 4.4). Each neuron was then represented as a point in a selectivity space where each axis corresponds to the regression coefficient with respect to one variable. For each network, we then compared the resulting distribution of points with a random shuffle corresponding to a multi-variate Gaussian with matching empirical covariance. (b) Illustration of the distribution of regression coefficients in selectivity space for two networks trained on respectively the multi-sensory (MDM) and context-dependent decision-making (CDM) tasks which received identical inputs (two stimuli *A* and *B* and two contextual cues) but required different outputs. The full selectivity space was four dimensional. The plots show two-dimensional projections of the selectivity distribution onto the plane defined by regression coefficients with respect to stimuli *A* and *B*. Gray ellipses correspond to the 1 s.d. ellipse of a Gaussian distribution with matching mean and covariance. (c) Distribution of angles between each point and its nearest neighbor in the selectivity space illustrated in panel b (colored histograms), compared with that of a matching multivariate Gaussian (black line). The mismatch between the two distributions was quantified using the ePAIRS test [Raposo et al., 2014; Hirokawa et al., 2019]. The mismatch was significant for the CDM task (*p* < 10^−7^), but not for the MDM task (*p* = 0.61). (d) Population structure in the selectivity space across networks and tasks: effect size of the ePAIRS test (see Methods 4.6) on the selectivity space for 100 networks trained on each of the five studied tasks (see Sup. Fig. S1 for p-values). Black bars represent 95% confidence intervals for null distributions. (e) To assess for population structure in connectivity, we focused on low-rank networks, where connectivity is fully specified by vectors over neurons [Mastrogiuseppe and Ostojic, 2018]. Each neuron is then characterized by one parameter on each vector (illustrated by colors, entries for a specific neuron are outlined in red), and can be represented as a point in connectivity space where each axis corresponds to the parameters on one vector. We assessed the presence of non-random population structure in that space using a procedure identical to the analysis of selectivity (c-d). (f) Illustration of the distribution of parameters in connectivity space for the two networks trained on respectively the MDM and CDM tasks. For these tasks, minimal trained networks were of rank *R* = 1 (Sup. Fig. S2), so that the connectivity space was of dimension 7 (four inputs, two recurrent vectors and one readout). The plots show two-dimensional projections of the full connectivity distribution onto the plane defined by parameters of contextual cues *A* and *B*. Gray ellipses correspond to the 1 s.d. ellipse of a Gaussian distribution with matching mean and covariance. (g) Comparison of distributions in connectivity space for trained networks and the randomized shuffles as in c. The difference is significant for the CDM task, but not for the MDM task. (h) Population structure in the connectivity space across networks and tasks: effect size of the ePAIRS test on the connectivity space for 100 networks trained on each of the five studied tasks (see Sup. Fig. S1 for p-values). (i) To identify the causal role of population structure on computations, we randomly generated new networks by resampling from the null distribution in connectivity space that preserved the mean and covariance structure but scrambled any non-random population structure. (j-k) In randomly resampled networks, the statistics of connectivity are by design identical to shuffles used for the ePAIRS test. (l) Performance of each randomly resampled network on its corresponding task as measured by accuracy.

#### Population structure in selectivity

We first asked if training on each task led to the emergence of non-random structure in selectivity, as previously assessed in the posterior parietal [Raposo et al., 2014] and prefrontal [Hirokawa et al., 2019] cortices. Following the approach developed in those studies, we represented each neuron as a point in a *selectivity space*, where each axis was given by the linear regression coefficient 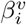 of neural firing rate with respect to a task variable *v* such as stimulus, decision or context (Fig. 1a). The dimension of the selectivity space ranged from 2 to 4 depending on the task (see Methods 4.4), and each trained network led to a distribution of points in that space (Fig. 1b). For each network, we compared the obtained distribution with a randomized shuffle corresponding to a multivariate Gaussian with matching empirical mean and covariance (Fig. 1b,c), and assessed the difference using the ePAIRS statistical test [Raposo et al., 2014; Hirokawa et al., 2019]. A non-significant outcome suggests an isotropic distribution of single-neuron selectivity, a situation that has been denoted as *fully-random population structure*, or non-categorical mixed selectivity [Raposo et al., 2014]. A statistically significant outcome instead indicates that neurons tend to be clustered along multiple axes of the selectivity space. Following Raposo et al. [2014]; Hirokawa et al. [2019], we refer to this situation as non-random mixed selectivity, or *non-random population structure*. The ePAIRS test on the selectivity distributions revealed the presence of non-random population structure for two out of the five tasks, the contextual decision-making and delay-match-to-sample tasks (proportion of statistically significant networks under the ePAIRS test, *p* < 0.05, Bonferroni corrected : DM task: 1/100, WM task: 6/100, MDM task: 10/100, CDM task: 87/100, DMS task: 100/100) (Fig. 1d). In particular, this analysis revealed a clear difference between the multi-sensory [Raposo et al., 2014] and context-dependent [Mante et al., 2013] decision making tasks, which had an identical input structure (two stimuli *A* and *B* and two contextual cues *A* and *B*, Fig. 3b) and therefore identical four-dimensional selectivity spaces, but required different mappings from inputs to outputs.

#### Population structure in connectivity

The selectivity in trained RNNs necessarily reflects the underlying connectivity [Mastrogiuseppe and Ostojic, 2018]. We therefore next sought to determine the presence of non-random population structure directly in the connectivity of networks trained on different tasks. Recent work has shown that training networks on simple tasks as considered here leads to a particular form of recurrent connectivity based on a *low-rank* structure [Schuessler et al., 2020b], meaning that the connectivity of each neuron is specified by a small number of parameters as detailed below. We leveraged this structure to represent trained networks in a low-dimensional *connectivity space*, and then assessed the presence of non-random population structure in that space using a procedure identical to the analysis of selectivity.

More specifically, we focused on RNNs constrained to have recurrent connectivity matrices *J*_*ij*_ of a fixed rank *R*, and for each task determined the minimal required *R* (Sup. Fig. S2). A matrix of rank *R* can in general be written as

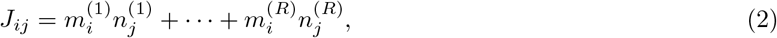

so that neuron *i* is characterized by 2*R* recurrent connectivity parameters 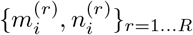. Each neuron moreover received *N*_*in*_ input weights, and sent out one readout weights (Fig. 1e), leading to a total of 2*R* + *N*_*in*_ + 1 parameters per neuron. We therefore represented the connectivity of each neuron as a point in a (2*R* + *N*_*in*_ + 1)-dimensional *connectivity space*, where each axis corresponds to entries along one connectivity vector. The connectivity of a full network can then be described as a distribution of points in that space (Fig. 1f). Similarly to the selectivity analysis, we assessed the presence of non-random population structure by comparing connectivity distributions of trained networks with randomized shuffles corresponding to multivariate Gaussians with matching empirical means and covariances, and quantified the deviations using the ePAIRS test. The results were consistent with the analysis of selectivity (Fig. 1g,h), and we again observed a clear gap between two groups of tasks (number of networks with statistically significant clustering for each task: DM: 3/100; WM: 5/100; MDM: 1/100; CDM: 100/100; DMS: 100/100; *p* < 0.05 with Bonferroni correction) (Fig. 1h). In particular and as was the case for selectivity, the MDM and CDM tasks led to opposite results although their connectivity spaces were identical (seven dimensional, with *N*_*in*_ = 4, *R* = 1, so that the total dimension was (2*R* + *N*_*in*_ + 1) = 7).

#### Computational role of population structure

The analyses of selectivity and connectivity provided a consistent picture on the absence or presence of non-random population structure across tasks. These analyses are however purely correlational, and do not allow us to infer a causal role of the observed structure. To determine when non-random population structure is computationally necessary, or conversely when random population structure is computationally sufficient, we therefore developed a new *resampling* analysis. For each task, we first generated new networks by sampling the connectivity parameters of each neuron from the randomized distribution used to assess structure in Fig. 1e-h, i.e. a multi-variate Gaussian distribution with mean and covariance matching the trained low-rank RNNs. This procedure preserved the rank of the connectivity (Fig. 1i), and the overall correlation structure of connectivity parameters, but scrambled any non-random population structure (Fig. 1j,k). We then quantified the performance of each randomly resampled network on the original task. This key analysis revealed that the randomly resampled networks led to a near perfect accuracy for the DM, WM and MDM tasks, but not for the CDM and DMS tasks (Fig. 1l). This demonstrates that, on one hand, random population structure is sufficient to implement the DM, WM and MDM tasks, while on the other hand non-random population structure is necessary for CDM and DMS tasks. These results held independently of the constraints on the rank of the connectivity, and in particular for unconstrained, full-rank networks in which only the learned part of the connectivity was resampled (Sup. Fig. S3).

It is important to stress that the performance of resampled networks is a much more direct assessment of the computational role of the non-random population structure than the analyses of selectivity and connectivity through the ePAIRS test. Indeed, the ePAIRS analyses can lead to false positives in which statistically significant non-random structure is found in both selectivity and connectivity although resampled networks with a single Gaussian still match the performance of the trained network (Sup. Fig. S4). As an illustration, networks trained on the DM task sometimes exhibited two diametrically opposed clusters in the connectivity space, suggesting two concurrent pools of self-excitatory populations, reminiscent of solutions previously found for this task [Wang, 2002; Williams et al., 2018; Schaeffer et al., 2020]. Generating resampled networks scrambled that structure, but still led to functioning networks, which showed that in the DM task the population structure does not bear an essential computational role, and might be an artifact of specific training parameters. Spurious structure can also appear in selectivity when the non-linearity is strongly engaged (Sup. Fig. S4).

In summary, our analyses of trained recurrent neural networks revealed that certain tasks can be implemented with a fully random population structure in both connectivity and selectivity, while others appeared to require additional organization in the connectivity that led to non-random structure in selectivity. We next sought to understand the mechanisms by which the population structure of connectivity determines the dynamics and the resulting computations. In a first step, we examined the situation in which the population structure is fully random. In a second step, in line with Hirokawa et al. [2019], we asked whether non-random population structure in the connectivity space could be represented in terms of separate clusters or sub-populations, and how this additional organization expands the computational capabilities of the network.

### 2.2 Interpreting computations in terms of latent dynamical systems

To unravel the mechanisms by which population structure impacts computations, we developed a method for interpreting the trained recurrent neural networks in terms of underlying low-dimensional dynamics [Vyas et al., 2020]. We specifically focused on networks with low-rank connectivity (Fig. 2a), which can be directly reduced to low-dimensional dynamical systems [Beiran et al., 2021] Here we first outline this model reduction approach, and next apply it on trained recurrent networks.

**Figure 2 :**
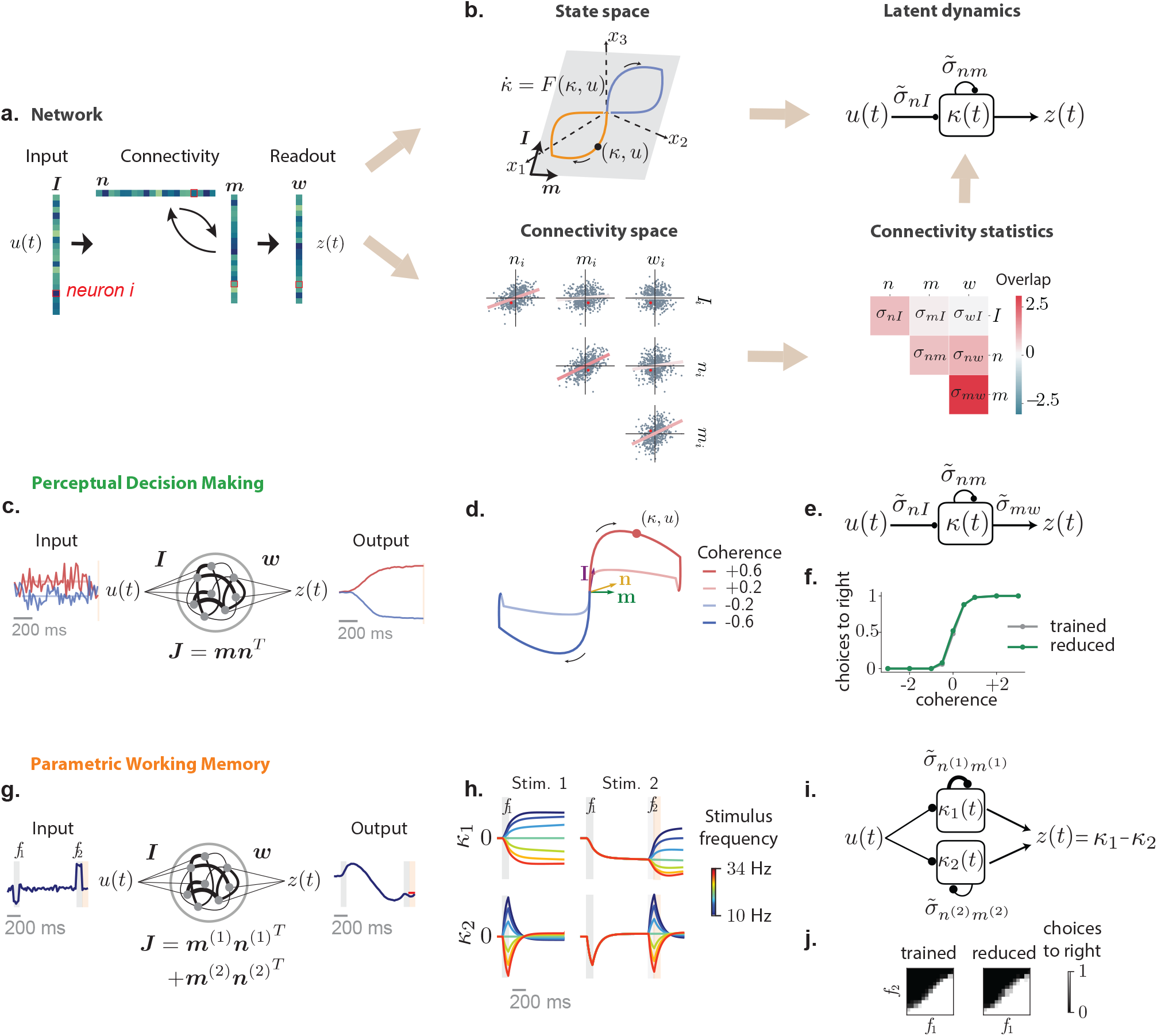
Low-dimensional latent dynamics explain computations in low-rank RNNs. (a-b) Reducing low-rank networks to low-dimensional latent dynamics. (a) The connectivity in a low-rank RNN is specified by a set of input, recurrent, and readout vectors over neurons. Here colors illustrate the entries of each neuron on these vectors, a specific neuron being outlined in red. (b) The connectivity vectors can be represented in two complementary manners that together determine low-dimensional dynamics. (top-left) In the *N*-dimensional state space, where each axis is the activity *x*_*i*_ of neuron *i*, connectivity vectors correspond to specific directions, illustrated as arrows. The connectivity constrains the trajectories of activity to lie in a low-dimensional subspace spanned by input vectors ***I***^(*s*)^ and recurrent vectors ***m***^(*r*)^. The activity trajectories (illustrated in color for two stimuli) are parametrized along those directions by inputs *u*_*s*_ and internal variables *κ*_*r*_, forming a latent dynamical system that fully determines the activity trajectory. (bottom left) The connectivity space provides a complementary representation, where each axis corresponds to a connectivity parameter along one vector. Any neuron (specific example in red) is represented as a point in this space, and the full network is described by the distribution of the cloud of points. Here we illustrate a four-dimensional distribution by its pairwise two-dimensional projections. (bottom right) A Gaussian distribution in connectivity space is specified by a covariance matrix that describes the shape of the point cloud, or equivalently the set of overlaps ***σ*** between all pairs of connectivity vectors. (top right) In that case, the latent dynamics can be reduced to an effective circuit (Eq. 5), in which each internal variable is represented as a unit that integrates external inputs, and interacts with itself (and other internal variables) through a set of effective couplings set by the connectivity overlaps. (c)-(e) Application to the perceptual decision making task. (c) A rank-one network was trained to output the sign of the mean of a noisy input signal. Example inputs and outputs are shown in red and blue for a positive and a negative input mean. (d) Low-dimensional trajectories in the two dimensional subspace spanned by vectors ***m*** and ***I***. (e) The latent dynamics are equivalent to an effective circuit governed by 2 effective couplings (Eq. 5), which are determined by the overlaps *σ*_*nI*_ and *σ*_*nm*_ of the vector ***n*** with ***I*** and *m* (see vectors in panel d). The readout from the network is set by the overlap *σ*_*mw*_ between the vectors ***m*** and ***w***. (f) Psychometric function showing the rate of positive outputs for the trained network, and a reduced network generated by controlling only three parameters corresponding to the effective couplings in e (see also Supplementary Fig. S5). (g)-(j) Application to the parametric working memory task. (g) A rank-two network was trained to compute the difference between two stimuli *f*_1_ and *f*_2_ separated by a variable delay. (h) The recurrent activity is described by two internal variables, *κ*_1_ and *κ*_2_ that correspond to activity along connectivity vectors ***m***^(1)^ and ***m***^(2)^. The variable *κ*_1_ acts as an integrator that encodes the stimuli persistently: *f*_1_ following the first stimulus, and *f*_1_ + *f*_2_ at the decision time following the second stimulus. The variable *κ*_2_ responds transiently to each stimulus, and therefore encodes *f*_2_ at the decision time. (i) The latent dynamics are described by an effective circuit where the two internal variables evolve independently, with different amounts of positive feedback (Eq. 47). (j) Psychometric response matrix for the trained network, and a reduced network generated by controlling only six parameters corresponding to the effective couplings in i. Each matrix displays the rate of positive responses for each combination of stimuli *f*_1_ and *f*_2_.

**Figure 3.**
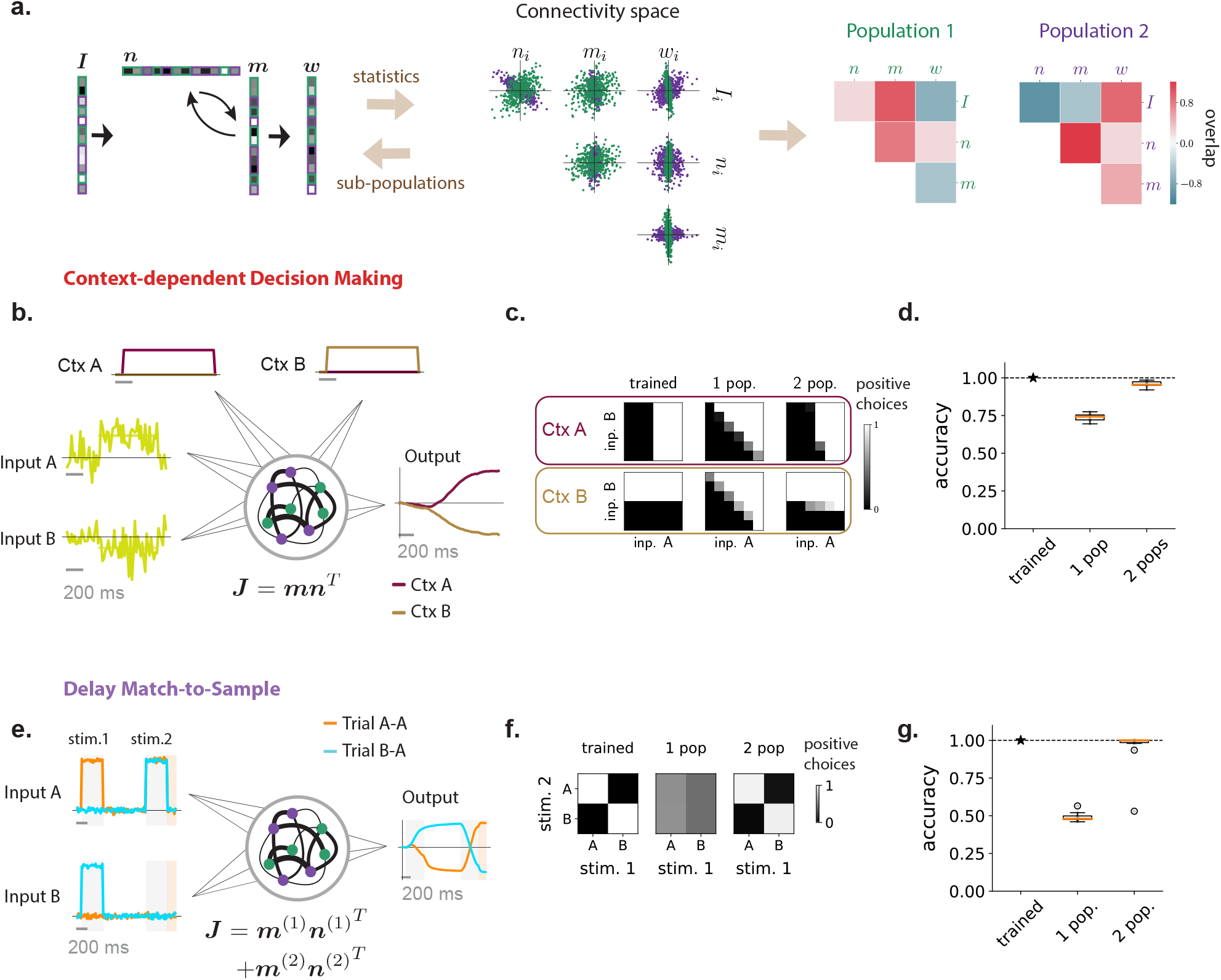
Multi-population connectivity structure captures the computational requirements for context-dependent tasks. (a) Illustration of the method for representing a low-rank connectivity structure in terms of multiple sub-populations. As in Fig. 1 the connectivity vectors (left panel) are first represented as a set of points in connectivity space, each point corresponding to connectivity parameters of one neuron. The center panel shows an illustration of different two-dimensional projections of the full distribution in connectivity space, which in this example is four dimensional. A mixture of Gaussians clustering algorithm then assigns every neuron to a sub-population based on the full distribution in connectivity space. The green and purple colors denote the two identified populations, which in this illustration have identical centers but different shapes. Each sub-population is therefore defined by a different set of covariances (right panel), that correspond to overlaps between input, recurrent and readout vectors shown in green and purple colors in the left panel. (b)-(d) Application to the context-dependent decision making task. (b) Networks received stimulus inputs consisting of a combination of two noisy features along two different input vectors, together with one of two contextual cues in each trial. Unit rank networks were trained to output the sign of the mean of the feature corresponding to the activated contextual cue. Here we illustrate two example trials sharing the same stimulus inputs and opposite contextual cues (context A activated in dark red, context B in pale brown), leading to opposite outputs. (c) Psychometric response matrices. Each matrix displays the rate of positive responses for each combination of means of stimulus features. Different rows show response matrices in different contexts. Different columns show response matrices for a trained network (left), and for networks generated by resampling connectivity from a single population (middle) or two populations (right). (d) Average accuracy of a trained network and of 10 draws of resampled single-population and two-population networks (boxplot, orange line: median, black box: first and third quartiles, outer lines: min and max in the limit of the median 1.5 interquartile intervals, standalone dots: outliers). (e)-(g) Application to the delayed match-to-sample task. (e) Networks received a sequence of two stimuli during two stimulation periods (in light gray) separated by a delay. Each stimulus belonged to one out two categories (A or B), each represented by a different input vector. Rank-two networks were trained to output during a response period (in light orange) a positive value if the two stimuli were identical, a negative value otherwise. Here we illustrate two trials with stimuli A-A and B-A respectively. (f) Psychometric response matrices. Rate of positive responses for each combination of first and second stimuli, for a trained network (left) and for networks generated by resampling connectivity from a single population (middle) or two populations (right). (g) Same as d for the DMS task.

In line with recent methods for analyzing large-scale neural activity [Buonomano and Maass, 2009; Cunningham and Byron, 2014; Gallego et al., 2017; Saxena and Cunningham, 2019], we start by representing the dynamics as trajectories *x*(*t*) = {*x*_*i*_(*t*)}_*i*=1…*N*_ in the high-dimensional state space, where the *i*-th dimension corresponds to the activation *x*_*i*_ of neuron *i* (Fig. 2b). For low-rank networks, the set of connectivity parameters can be interpreted as vectors over neurons that directly correspond to directions in the state-space (Fig. 2b). Indeed, each feed forward input corresponds to an input connectivity vector 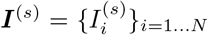, the low-rank parameters of the connectivity matrix (Eq. 2) can be represented as *R* pairs of recurrent connectivity vectors 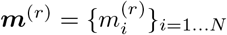 and 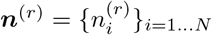 for *r* = … *R*, and the readout forms a vector ***w*** (Fig. 2a). Crucially, the low-rank connectivity structure directly restricts the dynamics to lie in a low-dimensional subspace spanned by the connectivity vectors ***I***^(*s*)^ and ***m***^(*r*)^ (Fig. 2b) [Mastrogiuseppe and Ostojic, 2018]. In line with dimensionality reduction approaches [Cunningham and Byron, 2014; Gallego et al., 2017], the collective activity in the network can therefore be fully described in terms of a small number of latent variables that quantify activity in this subspace [Mastrogiuseppe and Ostojic, 2018]. More specifically, ***x***(*t*) can be decomposed into a set of internal variables *κ*_*r*_ and inputs *u*_*s*_ which quantify respectively activity along ***m***^(*r*)^ and ***I***^(*s*)^ (Fig. 2b, Methods section 4.8.1), and correspond to recurrent and input-driven directions in state-space [Wang et al., 2018] :

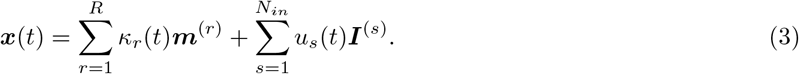

Altogether, the activity ***x***(*t*) is therefore embedded in a linear subspace of dimension *R*+*N*_*in*_ where *R* is the rank of the connectivity, and *N*_*in*_ is the dimensionality of feed-forward inputs. The dynamics are then fully specified by the evolution of the internal variables *κ* = {*κ*_*r*_}_*r*=1…*R*_ driven by inputs 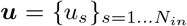. A mathematical analysis shows that the internal variables form a dynamical system [Remington et al., 2018; Vyas et al., 2020] with a temporal evolution of the form

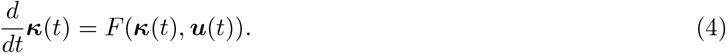

Here *F* is a non-linear function that determines the amount of change of ***κ*** at every time step. In the limit of large networks, the precise shape of *F* is set by the statistics of the connectivity across neurons (Methods section 4.8.4), i.e. precisely the distribution of points in the connectivity space that we previously examined in Fig. 1f,j. The connectivity in the network can therefore be represented in two complementary ways, either in terms of directions in the activity state-space (Fig. 2b top left) or in terms of distributions in the connectivity space (Fig. 2b bottom left), and these two representations together determine the low-dimensional latent dynamics.

In summary, in line with the computation-through-dynamics framework [Vyas et al., 2020], low-rank networks can be exactly reduced to low-dimensional, non-linear latent dynamical systems which determine the performed computations. We next examined how the population structure in trained recurrent networks impacts the resulting latent dynamical system. To facilitate the interpretation of computational mechanism, we focused on networks of minimal rank, which lead to latent dynamics of minimal dimensionality for each task (Methods 4.2). We later verify that the main conclusions carry over in absence of this constraint.

### 2.3 Latent dynamics and computations for fully random population structure

Our resampling analyses of trained RNNs revealed that a range of tasks could be performed by networks in which the population structure was fully random in connectivity space (Fig. 1l). We therefore first examine the latent dynamics underlying computations in that situation. Crucially, a fully random population structure limits the available parameter space, and strongly constrains the set of achievable latent dynamics independently of their dimensionality [Beiran et al., 2021]. We start be specifying these constraints on the dynamics, and show they nevertheless allow networks with random population structure to implement a range of tasks of increasing complexity by increasing the rank of the connectivity and therefore the dimensionality of the dynamics.

Networks with fully random population structure were defined in Fig. 1i-l as having distributions of connectivity parameters computationally equivalent to a Gaussian distribution. In such networks, the statistics of connectivity are therefore fully characterized by a set of covariances between connectivity parameters, each of which can be directly interpreted as the alignment, or overlap between two connectivity vectors (Fig. 2b bottom left, see Eq. 12). For this type of connectivity, a mean-field analysis shows that the latent low-dimensional dynamics can be directly reduced to an effective circuit, where internal variables *κ*_*r*_ integrate external inputs *u*_*s*_, and interact with each other through *effective couplings* set by the overlaps between connectivity vectors multiplied by a common, activity-dependent gain factor [Beiran et al., 2021]. In such reduced models, the role of individual parameters can then be analyzed in detail (Sup. Info).

As a concrete example, a unit-rank network (*R* = 1) with connectivity vectors ***m*** and ***n*** and a single feed-forward input vector ***I*** (*N*_*in*_ = 1) leads to two-dimensional activity (Eq. 3), fully described by a single internal variable *κ*(*t*) and a single external variable *u*(*t*) (Fig. 2b). The latent dynamics of *κ*(*t*) are given by

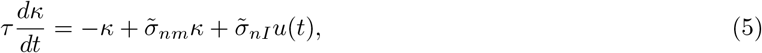

where 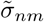 and 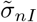 are effective couplings, that depend both on overlaps between connectivity vectors, and implicitly on *κ* and *u* through a gain factor, so that the full dynamics in Eq. 5 are non-linear despite their immediate appearance. More specifically, the effective couplings are defined as 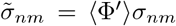 and 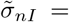 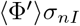, where *σ*_*nm*_ (resp. *σ*_*nI*_) is the fixed overlap between the vector ***n*** and the vector ***m*** (resp. ***I***). The connectivity vector ***n*** therefore selects inputs to the latent dynamics [Mastrogiuseppe and Ostojic, 2018]: the overlap between ***n*** and ***I*** controls how strongly the latent dynamics integrate feed-forward inputs, while the overlap between ***n*** and *m* controls the strength of positive feedback in the latent dynamics. Crucially, all the effective couplings are scaled by the same factor ⟨Φ′⟩ that represents the average gain of all neurons in the network. This gain depends on the activity in the network (Methods section 4.8.4), which makes the dynamics non-linear. The fact that all the effective couplings are scaled by the same factor however implies that, in networks with a fully random population structure, the overall form of the effective circuit is determined by the connectivity overlaps, which strongly limits the range of possible dynamics for the internal variables [Beiran et al., 2021]. Tasks for which a fully random population structure is sufficient are therefore those that can be implemented by a fixed effective circuit at the level of latent dynamics.

We first applied this model reduction framework to the perceptual decision making task, where a network received a noisy scalar stimulus *u*(*t*) along a random input vector, and was trained to report the sign of its temporal average along a random readout vector (Fig. 2c). Minimizing the rank of the trained recurrent connectivity matrix, we found that a unit-rank network was sufficient to solve the task (Sup. Fig. S2). The network connectivity was fully characterized by four connectivity vectors: the input vector ***I***, recurrent connectivity vectors ***n*** and ***m***, and the readout vector ***w*** (Fig. 2c). As a result, the activity *x*(*t*) evolved in a two-dimensional plane spanned by ***I*** and ***m***, and was fully described by two corresponding collective variables *u*(*t*) and *κ*(*t*) (Fig. 2d). The resampling analysis in Fig. 1l showed that trained networks were fully specified by the overlaps, or covariances between connectivity vectors, as generating new networks by sampling connectivity from a Gaussian distribution with identical covariances led to identical performance. The latent dynamics of *κ*(*t*) could then be reduced to a simple effective circuit (Fig. 2e, Eq. 5). Inspecting the values of covariances in the trained networks (Sup. Fig. S10) and analyzing the effective circuit (Sup. Fig. S5) revealed that the latent dynamics relied on a strong overlap *σ*_*nI*_ to integrate inputs, and an overlap *σ*_*nm*_ ≈ 1 to generate a long integration timescale via positive feedback. The internal variable *κ*(*t*) therefore represented integrated evidence along a direction in state space determined by the connectivity vector *m* (Fig. 2e,f). The readout vector *w* was aligned with *m*, so that the output directly corresponded to integrated evidence *κ*(*t*). Controlling only three parameters in the latent dynamics was sufficient to reproduce the psychometric input-output curve of the full trained network (Fig. 2f). Note that this network implementation is very similar to the implementation that has been proposed in previous work without making use of a learning algorithm [Mastrogiuseppe and Ostojic, 2018]. The findings from the perceptual decision task directly extended to the multi-sensory decision-making task [Raposo et al., 2014], in which the latent dynamics were identical, but integrated two inputs corresponding to two different stimulus features.

We next turned to the parametric working memory task [Romo et al., 1999], where two scalar stimuli *f*_1_ and *f*_2_ were successively presented along an identical input vector ***I***, and the network was trained to report the difference *f*_1_ − *f*_2_ between the values of the two stimuli (Fig. 2g). We found that this task required rank *R* = 2 recurrent connectivity (Sup. Fig. S2), so that the activity was constrained to the three-dimensional space spanned by ***I*** and the connectivity vectors ***m***^(1)^ and ***m***^(2)^. The low-dimensional dynamics could therefore be described by two internal variables *κ*_1_(*t*) and *κ*_2_(*t*) that represented activity along *m*^(1)^ and *m*^(2)^, and formed a two-dimensional dynamical system that integrated the input *u*(*t*) received along ***I***. The resampling analysis indicated that in this case also the trained connectivity was fully specified by covariances between connectivity vectors (Fig. 1i-l). Inspecting the connectivity distribution (Sup. Fig. S10) revealed that the two internal variables *κ*_1_ and *κ*_2_ did not directly interact, but instead independently integrated stimuli through dynamics given by Eq. 5 (Fig. 2i). For *κ*_1_, a strong overlap 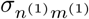 led to strong positive feedback that generated a persistent representation of the intensity *f*_1_ of the first stimulus along the direction of state space set by the connectivity vector ***m***^(1)^ (Fig. 2h top). For *κ*_2_, the overlap 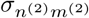, and therefore the positive feedback, was weaker, leading to a transient response that encoded the most recent stimulus along the direction *m*^(2)^ in the state space (Fig. 2h bottom). The readout vector *w* was aligned with both *m*^(1)^ and *m*^(2)^, but with overlaps of opposite signs, so that the output of the network in the decision period corresponded to the difference between *κ*_1_ and *κ*_2_, and therefore effectively *f*_2_ − *f*_1_ (Fig. 2i). Controlling only five parameters in the latent dynamics (Sup. Fig. S6) was therefore sufficient to reproduce the psychometric matrix describing the input-output mapping of the full trained network (Fig. 2j).

In summary, networks with random population structure can perform tasks of increasing complexity by relying on the dimensionality of recurrent dynamics to represent an increasing number of task-relevant latent variables. The random population structure however limits ways in which these latent variables can be combined by fixing the shape of the equivalent circuit. As a consequence, for more complex tasks a fully random population structure was not sufficient. We next sought to further elucidate this aspect.

### 2.4 Representing non-random connectivity structure with multiple populations

The resampling analysis in Fig. 1l indicated that tasks such as context-dependent decision-making and delayed-match-to-sample relied on a population structure in connectivity that was not fully random. To better understand the underlying structure and its computational role, we further examined RNNs trained on these two tasks, and asked whether their connectivity could be represented in terms of multiple populations. We first examined whether a multi-population connectivity structure is sufficient to implement the two tasks, and in a second step examined how such a structure modifies latent dynamics and expands their computational capacity.

To identify computationally-relevant populations, we took inspiration from Hirokawa et al. [2019], and first performed clustering analyses in the connectivity space where non-random population structure was found (Fig. 3a, Methods section 4.7). Each axis in that space represents entries along one connectivity vector, and each neuron corresponds to one point. Applying a Gaussian mixture clustering algorithm on the cloud of points formed by each trained network, we partitioned the neurons into separate sub-populations. In the trained networks, all clusters were centered close to the origin, but each had a different shape and orientation that corresponded to multiple peaks in the distribution of nearest-neighbour angles detected by the ePAIRS analysis (Fig. 1f-g). Each population was therefore characterized by a different set of covariances, or overlaps, between input, recurrent, and output connectivity vectors. We then extended our resampling approach from Fig. 1i-l, and generated new networks by first randomly assigning each neuron to a population, and then sampling its connectivity parameters from a Gaussian distribution with the fitted covariance structure. Finally, we inspected the performance of these randomly generated networks, and compared them with fully trained ones. By progressively increasing the number of fitted clusters, we determined the minimal number of populations needed to implement the task (Methods 4.7). Within this approach, networks with a fully random population structure such as those described in Fig. 2 correspond to a single overall population in connectivity space.

We first considered context-dependent decision making, where stimuli consisted of a combination of two scalar features that fluctuated in time [Mante et al., 2013]. Depending on a contextual cue, only one of the two features needed to be integrated (Fig. 3b), so that the same stimulus could require opposite responses, a hallmark of flexible input-output transformations [Fusi et al., 2016]. We implemented each stimulus feature and contextual cue as an independent input vector over the population, so that the dimension of feed-forward inputs was *N*_*in*_ = 4. We found that unit-rank connectivity was sufficient (Fig. S2), and focused on such networks. The analysis in Fig. 1l showed that generating networks by resampling connectivity from a single, fully-random population led to a strong degradation of the performance, although it remained above chance. A closer inspection of psychometric matrices representing input-output transforms (Fig. 3c) in different contexts revealed that single-population resampled networks in fact generated correct responses for stimuli requiring identical outputs in the two contexts, but failed for incongruent stimuli, for which responses needed to be flipped according to context (Fig. 3c right). This observation was not specific to unit rank networks, as randomizing population structure in higher rank (Sup. Fig. S11) and full rank networks (Sup. Fig. S3) led to a similar reduction in performance (Sup. Fig. S3). We therefore performed a clustering analysis in the connectivity space. The number of clusters varied across networks (Sup. Fig. S9), but the minimal required number was two. For such minimal networks, we found that randomly resampling from the corresponding mixture-of-Gaussian distribution led to an accuracy close to the original trained connectivity (Fig. 3d). In particular, the randomly generated networks correctly switched their response to incongruent stimuli across contexts (Fig. 3c), in contrast to networks with random population structure. This indicated that connectivity based on a structure in two populations was sufficient to implement the context-dependent decision-making task.

We next turned to the delayed-match-to-sample task [Miyashita, 1988; Engel and Wang, 2011; Chaisangmongkon et al., 2017], where two stimuli were interleaved by a variable delay period, and the network was trained to indicate in each trial whether the two stimuli were identical or different (Fig. 3e). This task involved flexible stimulus processing analogous to the context-dependent decision-making task because an identical stimulus presented in the second position required opposite responses depending on the stimulus presented in the first position (Fig. 3f). We found that this task required a rank two connectivity (Fig. S2), but, similarly to the context-dependent decision making task, a fully random population structure was not sufficient to perform the task, as networks generated by randomizing connectivity parameters reduced the output to chance level (Fig. 1l,3f,g). Fitting instead two clusters in the connectivity space showed that two sub-populations were sufficient, as networks generated by sampling connectivity based on a two-population structure led to a performance close to that of the fully trained network (Fig. 3g).

Altogether, our analyses based on clustering connectivity parameters, and randomly generating networks from the obtained multi-population distributions, indicated that connectivity distributions described by a small number of populations were sufficient to implement tasks requiring flexible input-output mappings. To identify the mechanistic role of this multi-population structure, we next examined how it impacted the latent dynamics implemented by trained networks.

### 2.5 Gain-based modulation of latent dynamics by multi-population connectivity

To unveil the mechanisms underlying flexible input-output mappings in networks with connectivity based on multiple populations, we examined how such a structure impacts the latent dynamics of internal variables. We first show that in contrast to a single-population, a multi-population structure allows external inputs to flexibly modulate the overall form of the circuit describing latent dynamics. We then show how this general principle applies specifically to the two flexible tasks described in Fig. 3. We focus here on networks with minimal rank and minimal number of populations, and show in the next section that the inferred predictions hold more generally.

In Fig. 3 we defined sub-populations as subsets of neurons characterized by different overlaps between input, recurrent and output connectivity vectors in a network of fixed rank. In a network with a multi-population structure, the number of internal variables describing low-dimensional dynamics is determined by the rank of the recurrent connectivity, as in networks without population structure (Fig. 2a). Remarkably, a mean-field analysis (Methods 4.8.4, [Beiran et al., 2021]) shows that the latent low-dimensional dynamics can still be represented in terms of an effective circuit where internal variables *κ*_*r*_ integrate inputs and interact with each other through effective couplings (Fig. 4a). The key effect of the multi-population structure is however to modify the form of the effective couplings and endow them with much greater flexibility than in the case of a single, fully random population. Indeed, in a network with a single population, the effective couplings were given by connectivity overlaps multiplied by a single, global gain factor, and modulating the gain therefore scaled all effective couplings simultaneously. In contrast, in networks with multiple populations, each population is described by its own set of overlaps between connectivity sub-vectors (Fig. 3a-c), and, importantly, by its own gain, which corresponds to the average slope *ϕ′*(*x*_*i*_) on the input-output nonlinearity of different neurons in the population. The effective couplings between inputs and internal variables are then given by a sum over populations of connectivity overlaps each weighted by the gain of the corresponding population (Methods Eq. (40)). As an illustration, in the case 374 of wo populations, the effective coupling between the input and the internal variable becomes

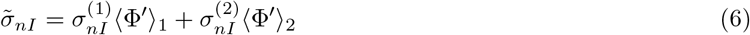

where 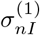 and 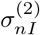 are the overlaps for each population between the input vector *I* and the input-selection vector *n*, while ⟨Φ′⟩_1_ and ⟨Φ′⟩_2_ are the gains of the two populations, which depend implicitly both on inputs and the values of internal variables.

**Figure 4.**
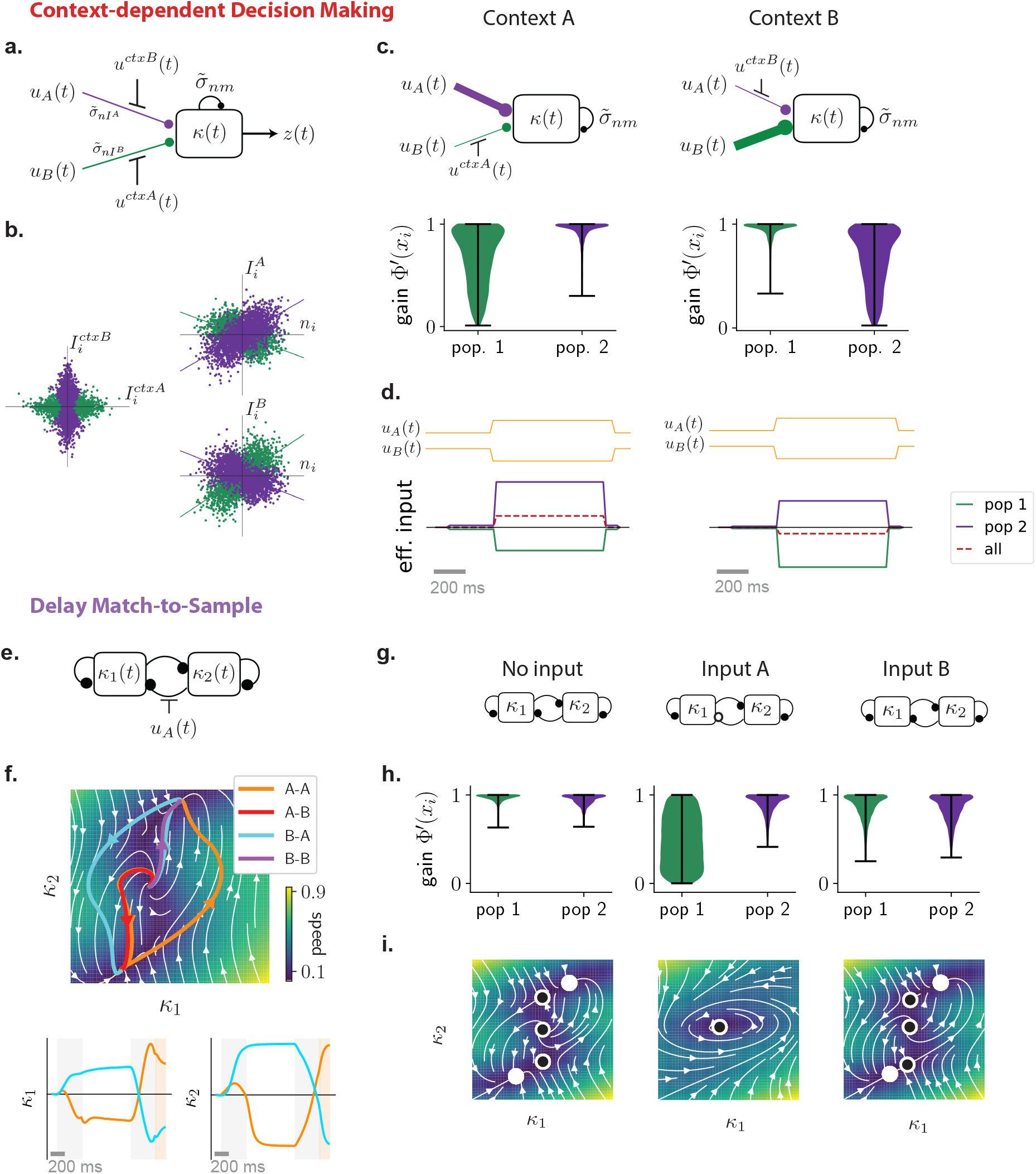
Mechanisms of computations based on a multi-population connectivity structure. (a) Circuit diagram representing latent dynamics in the reduced model of context-dependent decision-making task (Eq. 7). The internal variable *κ* is represented as a unit that integrates the two stimulus features *u*_*A*_ and *u*_*B*_ through effective couplings 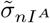 and 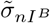. The coupling 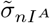 corresponds to the overlap between vectors ***n*** and ***I***^*A*^ for population 1, multiplied by the gain of that population, while 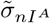 is the overlap between vectors ***n*** and ***I***^*B*^ for population 2, multiplied its gain. Contextual inputs 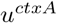 and 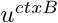 modulate the gains of the two populations and therefore the effective couplings that govern which stimulus feature is integrated. Lines with round ends represent effective couplings, lines with straight ends represent gain modulation. (b) Three two-dimensional projections of the six-dimensional connectivity space for a network trained on the task. Each point represents the parameters of one neuron, and the two colors indicate populations found by clustering neurons within the full six-dimensional space (Fig. 3a). Left: plane defined by components of the contextual-cue vectors ***I***^*ctxA*^ and ***I***^*ctxB*^ ; right: two planes defined by components on the input-selection vector ***n*** and the two stimulus feature vectors ***I***^*A*^ and ***I***^*B*^ (lines show linear regressions for each population). (c) Effective circuits in each context (top) and corresponding gains of neurons in each population (bottom). For each neuron *i*, the gain is defined as the slope of *ϕ*(*x*_*i*_) during stimulation period. Violin plots showing the distribution of gains for all neurons in each population in context A (left) and B (right). In context A, the average gain of neurons in population 1 (green) is lower than population 2 (purple), which decreases the effective connectivity between input feature B and the latent variable (top left circuit). The opposite happens in context B (top right circuit). (d) Effective inputs to the latent variable *κ* in the two context (bottom) in response to the same stimulus input (top). Solid lines show inputs mediated by each population (defined as 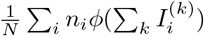, see Methods Eq. (38)), the dashed line shows the total input, which changes signs between the two contexts, leading to opposite responses. (e) Circuit diagram representing latent dynamics for a minimal network trained on the DMS task (Eq. 53). The network was of rank two, so that the latent dynamics were described by two internal variables *κ*_1_ and *κ*_2_. Input A acts as a modulator on the recurrent interactions between the two internal variables. (f) Top: Dynamical landscape for the autonomous latent dynamics in the *κ*_1_ *κ*_2_ plane. Colored lines depict trajectories corresponding to the 4 types of trials in the task. Background color and white lines encode the speed and direction of the dynamics in absence of inputs. Bottom: temporal evolution of *κ*_1_ and *κ*_2_ in two trials in which the second stimulus was identical, but the first one different. (g) Effective circuit diagrams in absence of inputs (left), and when input A (middle) or input B (right) are present. Filled circles denote positive coupling, open circles negative coupling. Input A in particular induces a negative feedback from *κ*_2_ to *κ*_1_. (h) Distribution of neural gains for each populations, in the three situations described above. The gain of population 1 (green) is specifically modulated by input A. (i) Dynamical landscapes in the 3 situations described above (see Methods). Filled and empty circles indicate respectively stable and unstable fixed points. The negative feedback induced by input A causes a limit cycle to appear in the latent dynamics.

Crucially, additional inputs restricted to a given population can modulate its gain independently of other populations by shifting the position of neurons on the non-linear input-output function. Additional inputs can thereby shape latent dynamics without directly driving them, but by modifying effective couplings (Methods 4.8.5). Indeed, as pointed out earlier, only inputs corresponding to input vectors aligned with the input-selection vectors ***n***^(*r*)^ directly drive internal variables through a non-zero effective coupling. In contrast, inputs corresponding to input vectors orthogonal to input-selection vectors do not directly drive the latent dynamics, but do modulate the values of the gain ⟨Φ′⟩_*p*_ of each population, and therefore the effective couplings. As a consequence, depending on the geometry between input vectors and input-selection vectors, different sets of inputs can play distinct roles of drivers and modulators [Sherman and Guillery, 1998] at the level of the effective circuit describing latent dynamics. Such a mechanism considerably extends the range of possible dynamics with respect to the case of a single overall population. In particular, modulating the gains of different populations allows the network to flexibly remodel the effective circuit formed by collective variables in different trials or epochs according to the demands of the task, in contrast to the single-population case, where the form of the effective circuit is fixed. We next describe how this general mechanism explains the computations in the two flexible tasks of Fig. 3.

For the context-dependent decision-making task, the minimal trained networks were of unit rank and consisted of two sub-populations (Fig. 3f). Analyzing the statistics of input and connectivity vectors for each population, we found that the input vectors ***I***^*A*^ and ***I***^*B*^ corresponding to the two stimulus features *u*_*A*_ and *u*_*B*_ had different overlaps with the input-selection vector ***n*** in the two populations (Fig. 4b right) so that the two stimulus features *u*_*A*_ and *u*_*B*_ acted as drivers of latent dynamics. The contextual input vectors ***I***^*ctxA*^ and ***I***^*ctxB*^ in contrast had weak overlaps with the input-selection vector *n* (Sup. Fig. S10), but strongly different amplitudes on the two populations (Fig. 4b left). They therefore modified the gains of the two populations in an opposite manner (Fig. 4c bottom), and played the role of modulators that modified the form of the effective circuit describing latent dynamics in each context (Fig. 4c top). More specifically, the latent dynamics of the internal variable *κ* could be approximated by (Methods 4.8.4 and Sup. Fig. S7):

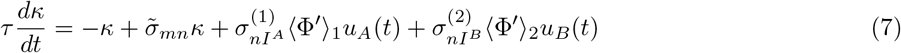

where ⟨Φ′⟩_1_ and ⟨Φ′⟩_2_ are the average gains of the two populations, 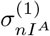 the overlap for the first population between the input vector for stimulus feature *A* and the input-selection vector ***n***, and 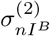 the overlap for the second population between ***n*** and the input vector for stimulus feature *B*. By modulating the gains of the two populations in a differential manner between the two contexts (Fig. 4c bottom), the contextual cues controlled the effective couplings between stimulus inputs and the internal variable *κ*, and determined which feature was integrated by the internal variable in each context (Fig. 4d). This mechanism implemented an effective input gating, but only at the level of the latent dynamics of the internal variable *κ* that integrated relevant evidence. Importantly, as observed in experimental data [Mante et al., 2013], on the level of the full network, the two stimulus features were instead equally represented in both contexts, but along directions in state space orthogonal to the direction *n* that encoded internal collective variable (Sup. Fig. S12) as observed in experimental data [Mante et al., 2013].

For the delayed-match-to-sample task, we found that the multi-population structure also led to a modulation of latent dynamics, but across task epochs rather than across trials. Fig. 4e-i describes an example minimal network implementing this task, where one of the stimuli played the role of a modulatory input, and transiently modified the latent dynamics when presented (Fig. 4e,g,i). More specifically, the network was of rank two, so that the latent dynamics were described by effective interactions between two internal variables *κ*_1_ and *κ*_2_ (Fig. 4e), and could be visualised in terms of a flow in a dynamical landscape in the *κ*_1_ *κ*_2_ plane (Fig. 4f). The minimal connectivity moreover consisted of two populations (Fig. 3i). Stimulus A modulated the gain of the first population (Fig. 4h), and therefore, when presented, modified the effective couplings in the latent dynamics and the dynamical landscape (Fig. 4i and Sup. Fig. S8)). The main effect of the inputs was therefore to shape the trajectories of internal variables by modulating the dynamical landscape at different trial epochs (Fig. 4i and Sup. Fig. S13). In particular, stimulus A strongly enhanced negative feedback (Fig. 4g), which led to a limit-cycle in the dynamics that opened a fast transient channel that could flip neural activity in the *κ*_1_ *κ*_2_ plane [Chaisangmongkon et al., 2017]. The four trials in the task therefore corresponded to different sequences of dynamical landscapes (Fig. 4i) leading to different neural trajectories and final states determining the correct behavioral outputs (Sup. Fig. S13).

In summary, we found that networks with multiple sub-populations implemented flexible computations by exploiting gain modulation to modify effective couplings between collective variables. The minimal solutions for the two tasks displayed in Fig. 3 and Fig. 4 illustrate two different variants of this general mechanism. In the context-dependent decision-making task, the sensory inputs acted as drivers of the internal dynamics, and contextual inputs as gain modulators that controlled effective couplings between the sensory inputs and the internal collective variable. In contrast, in the delayed-match-to-sample task, sensory inputs acted as modulators of recurrent interactions, and gain modulation controlled only the effective couplings between the two internal variables. More generally, modulations of inputs and modulations of recurrent interactions could be combined to implement more complex tasks.

### 2.6 Predictions for neural selectivity and inactivations

Analyzing networks of minimal rank and minimal number of population allowed us to identify the mechanisms underlying computations based on a multi-population structure in connectivity. We next sought to generate predictions of the identified mechanisms that are experimentally testable without access to details of the connectivity. We then tested these predictions on networks with a higher number of populations or higher rank, obtained by varying the constraints used during training. We focus here specifically on the context-dependent decision-making (CDM) task, and contrast it with the multi-sensory decision-making (MDM) task, for which networks received an identical input structure, but were required to produce an output independent of context.

For the CDM task, reducing the trained networks to effective circuits revealed that the key computations relied on a differential gain-modulation of separate populations by contextual inputs. For each neuron, contextual cues set the functioning point of the neuron on its non-linearity, and the gain of its response to incoming stimuli. A direct implication is that neurons more strongly modulated by context cues change more strongly their gain across contexts, and thereby the amplitude of their responses to stimulus features (Fig. 5a). An ensuing prediction at the level of selectivity of individual neurons is therefore that the pre-stimulus selectivity to context should be correlated with the change across contexts of regression coefficients to stimulus features (Fig. 5b). Our analyses therefore predict a specific form of multiplicative interactions, or non-linear mixed selectivity to stimulus features and context cues [Rigotti et al., 2013], but also imply that the two populations can be identified based on their selectivity to context (Fig. 5b).

**Figure 5.**
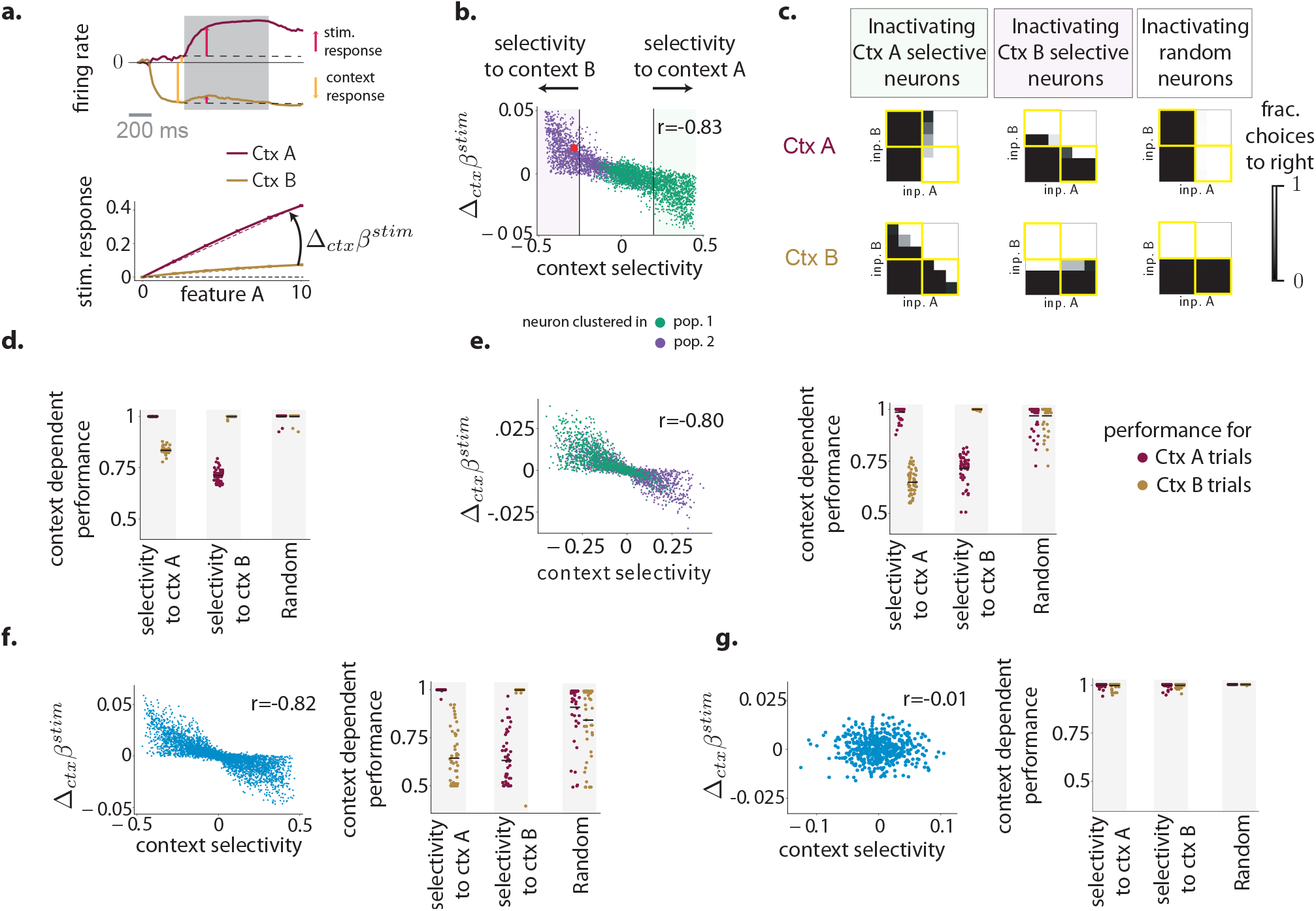
Predictions for neural selectivity and inactivations. (a-d) Predictions for the context-dependent decision-making task based on the minimal unit-rank, two-population network (Fig. 4a). (a) Context-dependent stimulus response for an example neuron that is strongly modulated by one of the contextual cues before stimulus onset. Top: response to an identical stimulus in two contexts, aligned to the time of presentation of the contextual cue. The grey box indicates the stimulus-presentation period. The context response was defined as the change of pre-stimulus baseline induced by the contextual cue (orange arrow). The stimulus response was defined in each context as the deviation from the pre-stimulus baseline (red arrows). Bottom: summary of context-dependent responses of the same neuron to stimuli with increasing strength of feature *A*. In each context, we computed the regression coefficient with respect to feature strength (dashed lines), and computed the change in stimulus selectivity Δ_*ctx*_*β*^*stim*^ as the difference between regression coefficients (see Methods 4.4, Eq. (21)). (b) Interaction between pre-stimulus context selectivity and the change in stimulus selectivity at the population level. For each neuron, a point shows the change in stimulus selectivity across contexts (as defined in (a)) versus its selectivity to context during pre-stimulus baseline (see Methods 4.4, Eq. (20)). Dots are colored according to the population to which neurons were assigned by the clustering procedure (Fig. 4). The red dot corresponds to the example neuron shown in (a). (c) Inactivations based on context selectivity lead to specific performance deficits. Psychometric response matrices (as defined in Fig. 3e) when inactivating the 256 out of 1024 neurons with highest positive context selectivity (left), highest negative context selectivity (middle) or randomly chosen across the whole network (right). (d) Summary of the effect of inactivation on performance. Each dot displays the context-dependent performance defined as the performance on non-congruent stimuli (yellow squares in the psychometric matrices in c), for one random subset of 256 out of 1024 inactivated neurons. (e-g) Tests of the predictions for selectivity (left panels) and inactivations (right panels) on: (e) a unit-rank network consisting of three populations (e, see Sup. Fig. S9); (f) a network trained without a rank constraint; (g) a network trained on the multi-sensory decision-making (MDM) task.

The multiplicative interaction between context and stimulus selectivity is a necessary, but not a sufficient condition for implementing context-dependent responding. A second, necessary component of the computational mechanism is that each population integrates dominantly one of the two features into the latent dynamics, as seen previously from the overlaps between the input vectors and the input-selection vectors (Fig. 4b right). This leads to a specific prediction for inactivation experiments: inactivating separately populations defined by their selectivity to context disrupts performance in one context, while leaving the other intact (Fig. 5c-d). In contrast, inactivating a random subset of neurons leads only to a slight overall decrease in performance independently of the context (Fig. 5c-d).

We first tested the two predictions on networks constrained to be of minimal, unit rank, but in which clustering analyses in connectivity space revealed more than two populations (Sup. Fig. S9), as in [Yang et al., 2019]. The two predictions for selectivity and inactivations were therefore directly borne out for such networks (Fig. 5e). We next turned to networks trained without rank constraint, and tested the two predictions without analyzing connectivity, as would be the case in experimental studies. The two predictions were again borne out (Fig. 5f), confirming that key aspects of the computational mechanisms extend to networks in which the connectivity structure was of higher rank, and the dynamics higher dimensional.

Finally we examined unit-rank networks trained on the MDM task. Such networks received an input structure identical to the CDM task, consisting of two stimulus features and two context cues. In contrast to the CDM task, the network was required to average the two stimulus features, and contextual cues were irrelevant, so that a fully random population structure was sufficient to perform the task (Fig. 1l). We therefore expected that the two predictions made for the CDM task do not necessarily hold in this case. We indeed found that training networks on the MDM task led to weaker selectivity to context, and weaker correlation between context selectivity and the change in stimulus selectivity (Fig. 5f). Specific neurons still exhibited selectivity to contextual cues, but inactivating them led to changes in performance similar to inactivating a random subset of neurons (Fig. 5f). Importantly, we controlled for the effect of context selectivity strength by manually increasing the amplitude of contextual inputs until the context selectivity matched that of networks trained on the CDM task. This increased the correlation between context selectivity and the change in stimulus selectivity but did not increase the impact of inactivating context-selective neurons (Sup. Fig. S15).

Altogether, our analyses therefore show that inactivating specific selectivity-defined populations leads to specific effects on performance in networks predicted to rely on non-random population structure, but not in networks for which population structure is expected to be computationally irrelevant.

### 2.7 Implications for multi-tasking

A recent study has reported that multiple populations emerge in networks trained simultaneously on multiple tasks, and can be repurposed across tasks [Yang et al., 2019]. Our results more specifically suggest that a multi-population structure in connectivity is needed only when an identical stimulus requires different outputs depending on the context set by the performed task. While this is the case in many multi-tasking situations, concurrent tasks are alternatively often based on different sets of stimuli [Cromer et al., 2010; Fritz et al., 2010; Elgueda et al., 2019]. Here we show that the reduced models developed by analyzing networks trained on individual tasks can be used to build networks that perform multiple tasks in parallel (Fig. 6). More specifically, multiple tasks on an identical set of stimuli can be performed by combining and repurposing multiple populations, while in contrast multiple tasks on separate sets of stimuli can be performed with a single population by relying on dynamics in orthogonal subspaces [Duncker et al., 2020; Zenke et al., 2017]. As a result, when identical stimuli are processed, some individual neurons exhibit task-specialisation, while for separate sets of stimuli all neurons are multi-taskers, and contribute to multiple tasks in parallel. These findings are in direct agreement with the activity of neurons in the prefrontal cortex during flexible categorisation, which show specialisation when identical stimuli are processed [Roy et al., 2010], and multi-tasking when separate stimuli sets are used [Cromer et al., 2010].

**Figure 6.**
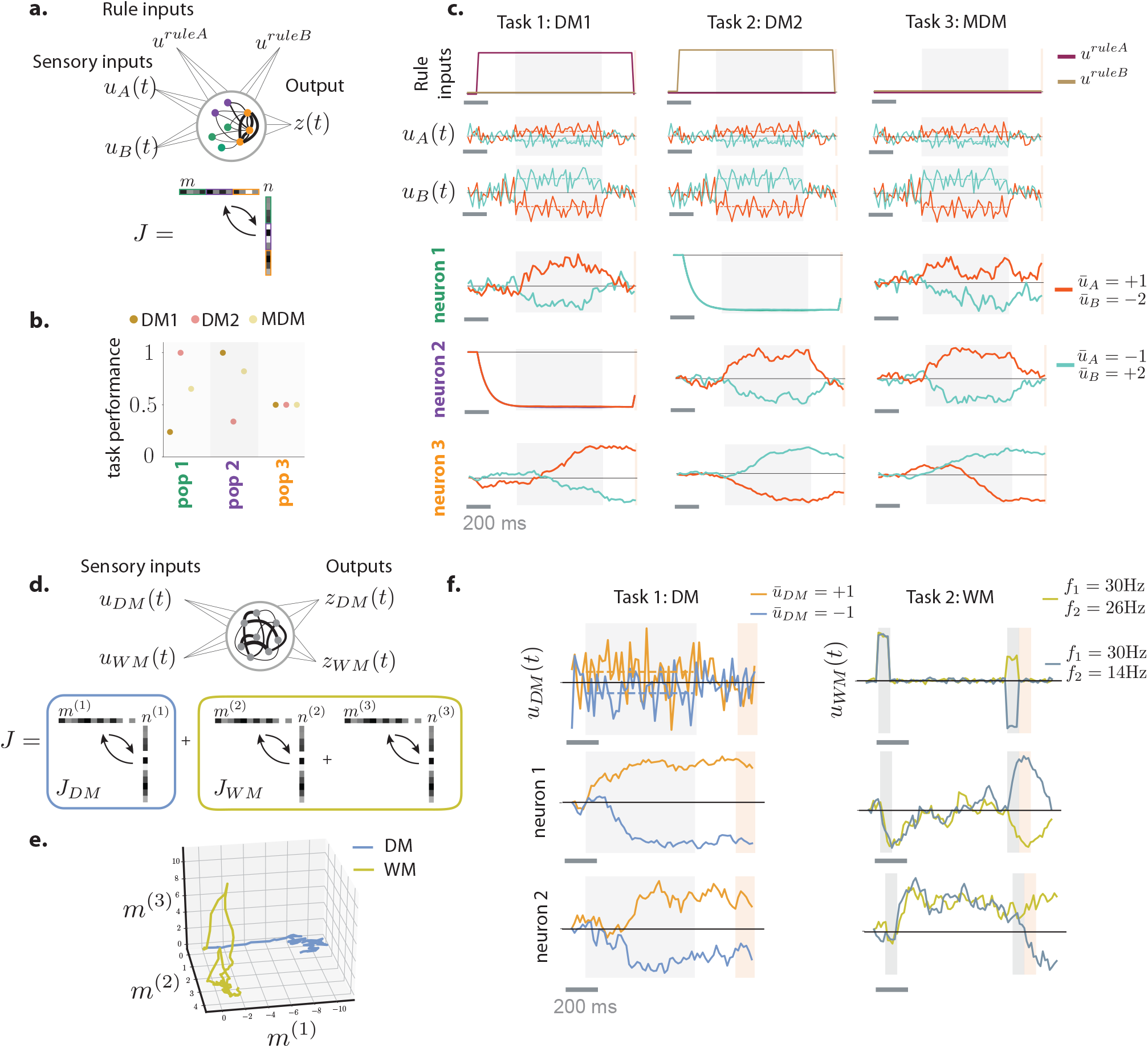
Implications of multi-population structure for multi-tasking. (a) A network performing three different tasks on the same set of stimuli consisting of two features *u*_*A*_ and *u*_*B*_: decision-making based on *u*_*A*_ (DM1), decision-making based on *u*_*B*_ (DM2), decision-making based on integrating *u*_*A*_ and *u*_*B*_ (MDM). The model is obtained from the unit-rank network performing the CDM task based on three populations indicated in color. (b) Effects on the performance of individual tasks when specific populations are inactivated. In each case one third of the neurons in the network is inactivated, corresponding to one of the three populations. (c) Illustration of task specialization of different populations. The orange population plays the role of an integrator, and participates to all tasks. Green and purple populations respectively relay *u*_*A*_ and *u*_*B*_. Different columns correspond to different tasks. Top three rows display stimulus and rule inputs. Bottom three rows display single unit activities of three selected neurons (one in each population) in two trials of each task. (d) A network performing two different tasks on distinct sets of stimuli, the decision-making (DM) task on *u*_*DM*_, and the working-memory task on *u*_*WM*_. This network is obtained by superposing the low-rank recurrent connectivity matrices corresponding to the two tasks (illustrated at the bottom). (e) The two tasks rely on neural activity in orthogonal subspaces of the state space. Each subspace is determined by the input connectivity vectors of the corresponding task. (f) Illustration of multi-tasking of two example neurons.

To illustrate task-specialization, we first consider a network that receives stimuli composed of two sensory features, and depending on a rule cue performs one out of three different tasks on them : perceptual decision-making on the first stimulus feature, perceptual decision-making on the second stimulus feature, or integration of the two features as in the multi-sensory decision making task (Fig. 6a). This multi-tasking setup is in fact a direct extension of context-dependent decision-making, and we implemented it using a simplified network based on the CDM task, consisting of unit-rank connectivity with three separate populations (Sup. Fig. S9). In that network, each population has a well defined computational role. One of the populations plays the role of an integrator, and endows the latent dynamics with a long time-scale through strong positive feedback, allowing the network to integrate evidence. That population is repurposed across all tasks (Fig. 6c brown traces), and inactivating it leads to performance degradation on all three tasks (Fig. 6b). The other two populations relay separately the two sensory features into the latent dynamics, as in the CDM task (Fig. 4b,c,d). Each of them participates in only two of the three tasks, as corroborated by changes in task performance after selective inactivations (Fig. 6b). Neurons belonging to these two populations are therefore specialised for specific tasks, as seen in their task-specific responses to stimuli (Fig. 6c green and purple neurons).

We next illustrate multi-tasking in a network that performs two tasks on distinct sets of stimuli, the perceptual decision-making (DM) and the parametric working-memory (WM) tasks (Fig. 6d). Such a network can be obtained by directly superposing the connectivity matrices ***J***_*DM*_ and ***J***_*WM*_ of two minimal networks of rank one and two that perform the individual tasks with random population structure (Fig. 2). The resulting connectivity ***J*** = ***J***_*DM*_ + ***J***_*W M*_ is of rank three, and has a random population structure. The corresponding latent dynamics are based on a recurrent sub-space of dimension three, and the two tasks rely on two orthogonal subspaces with one dimension implementing the DM task, and the other two implementing the WM task (Fig. 6e). Because of the random population structure, each neuron is a random combination of collective variables corresponding to different tasks, so that all neurons display multi-tasking activity (Fig. 6e).

## 3 Discussion

The goal of this study was to determine whether and when a non-random population structure is necessary for networks to perform a specific computation based on recurrent dynamics. To address this question, we first trained recurrent neural networks on a range of standard systems neuroscience tasks, and examined the emerging population structure in the selectivity and connectivity, and its relationship with the computations. We then identified underlying mechanisms by extracting the latent low-dimensional dynamics. Although a number of tasks could be implemented with random population structure in connectivity, we found that tasks based on flexible input-output mappings instead appeared to require an additional structure that could be accurately approximated in terms of a small number of sub-populations which played functionally distinct roles.

The starting motivation of this work was the apparent discrepancy between the experimental results of Raposo et al. [2014] and Hirokawa et al. [2019]. Analyzing neural activity in the rat posterior parietal cortex during a multi-sensory decision-making task, Raposo et al. [2014] found no evidence for non-random population structure in selectivity. Applying identical analyses to the prefrontal cortex, Hirokawa et al. [2019] instead identified population structure in activity during a more complex task that combined perceptual and value-guided decisions. Our results suggest that the difference between tasks provides a possible explanation for these diverging conclusions. Examining networks trained on an abstracted version of the multi-sensory integration task of Raposo et al. [2014], we found that a non-random population structure was not needed. Implementing a full version of the task used in Hirokawa et al. [2019] would have required reinforcement learning that falls beyond the scope of the supervised methods for training networks used here. The core component of that task was however a flexible weighing of two sensory features depending on the context set by reward history. That requirement of context-dependent weighing of input streams is in fact identical to the context-dependent decision-making task, in which all-or-none weights were assigned to the two stimulus features depending on the contextual cues. The gain-modulation mechanism underlying networks that performed the CDM task can more generally assign graded weights to each feature as required for the task of Hirokawa et al. [2019]. This mechanism requires multiple populations, so that our analyses predict that a non-random population structure is needed for the task used in Hirokawa et al. [2019].

Fundamental theoretical results guarantee that unconstrained recurrent neural networks are able to approximate any input-output function if the number of neurons is large enough [Doya, 1993]. Here we have instead sought to determine how far this property extends to networks with a random population structure in connectivity, as defined based on the null hypothesis of the analyses of Raposo et al. [2014] and Hirokawa et al. [2019]. A key point is that such a random population structure in fact sets constraints on the parameter space that the connectivity can explore, by precluding distributions of connectivity parameters more complex than a single Gaussian population. In a previous study, we demonstrated that low-rank recurrent networks with such a random population structure can generate only a limited range of autonomous dynamics independently of their rank, while having multiple sub-populations instead allows networks to approximate any low-dimensional dynamical system [Beiran et al., 2021]. Here we showed that these theoretical findings directly allow us to interpret the computational role of population structure in networks trained on neuroscience tasks.

We found that in trained networks relying on a non-random population structure, connectivity could be accurately described by a small number of sub-populations. Mechanistically, the role of such a sub-population structure can be understood from two perspectives. From the neural state-space perspective, the collective dynamics explore a low-dimensional recurrent subspace, and the sub-population structure shapes the non-linear dynamical landscape of the activity in this subspace [Sussillo and Barak, 2013]. Specifically, different inputs differentially activate different populations, and shift the recurrent sub-space into different regions of the state-space with different non-linear dynamical landscapes. A complementary picture emerges from the perspective of the effective circuits which describe the low-dimensional latent dynamics in terms of interactions between collective variables through effective couplings (Fig. 4c,g). In that picture, the sub-population structure allows inputs to control the effective couplings by modulating the average gain of different sub-populations. The computations then rely on two functionally distinct types of inputs: drivers that directly entrain the collective variables, and modulators that shape the gains of the different sub-populations, and thereby the interactions between collective variables. Interestingly, gain modulation has long been posited as a mechanism underlying selective attention [Rabinowitz et al., 2015], a type of processing closely related to flexible input-output tasks considered here. While patterns of gain modulation [Salinas and Thier, 2000; Ferguson and Cardin, 2020], and the distinction between drivers and modulators [Sherman and Guillery, 1998] are fundamentally physiological concepts, here we found that an analogous mechanism emerges in abstract trained networks at the collective level of latent dynamics. Note that in our framework, drivers and modulators are indistinguishable at the single cell level, where they both correspond to additive inputs (in contrast to eg neuro-modulation that may multiplicatively control the gain of individual neurons, see [Stroud et al., 2018]). The functional distinction between drivers and modulators instead stems from the relation between the collective pattern of inputs, and the recurrent connectivity in the network. Our analyses therefore establish a bridge between two levels of description, in terms of circuits, and in terms of collective dynamics [Barack and Krakauer, 2021].

To focus on the functional role of population structure, before training we initialized our networks with fully unstructured connectivity, and in particular did not include any explicit anatomical constraints such as Dale’s law. Our analyses nevertheless show that the non-random population structure that emerges through training can be accurately described in terms of abstract sub-populations, defined as clusters in the connectivity space. What could be the physiological counter-parts of the different functional sub-populations that we identified? There are at least two distinct possibilities. In the network trained on the context-dependent decision-making task, we found that the two sub-populations differed only in the relationship of their connectivity with respect to feed-forward and contextual inputs. Such sub-populations therefore bear an analogy with input- and output-defined cortical populations such as for instance defined by inputs from the thalamus [Harris and Mrsic-Flogel, 2013; Schmitt et al., 2017] or outputs to the striatum [Znamenskiy and Zador, 2013]. In the network trained on the delayed-match-to sample task, the two sub-populations instead differed at the level of recurrent connectivity: one population implemented positive, and the other negative feedback, the two being in general balanced (Sup. Fig. S10). This situation is reminiscent of excitatory and inhibitory sub-populations, which effectively implement positive and negative feedback in biological networks. Note that we do not mean to suggest that such population structure emerges biologically over the course of learning a task. Here we used artificial network training protocols to identify computational constraints, and we did not assume that they correspond to biological task-learning mediated by synaptic plasticity. The population wiring structure that emerges through network training could for instance be interpreted as the result of evolutionary selection leading to anatomic structure encoded at the genetic or developmental level [Zador, 2019].

Previous studies have reported that when training networks on a given task, some aspects of the solutions are invariant [Maheswaranathan et al., 2019] while others depend on the details of the implementation [Yang et al., 2019; Duncker et al., 2020; Flesch et al., 2021]. Our analyses confirmed these observations. Our main result for the computational requirement of non-random population structure in connectivity (Fig.1l) held independently of the details of the training, and in particular in absence of constraints on the rank of the network (Sup. Fig. S3). For tasks requiring a non-random population structure, the number of sub-populations needed to approximate connectivity however varied across networks (Sup. Fig. S9). For those tasks, our results show that a single global population is insufficient and that fundamental computational mechanisms are conserved across a range of different networks (Fig. 5). Our analyses however do not predict the specific dimensionality or number of populations to be expected. More systematic model selection could for instance be performed by further constraining recurrent neural networks based on recorded neural activity [Rajan et al., 2016; Aoi et al., 2020].

The fact that neurons are selective to mixtures of task variables rather than individual features has emerged as one of the defining properties of higher order areas in the mammalian cortex [Fusi et al., 2016]. Following up on analyses of Hirokawa et al. [2019], our results further clarify that mixed selectivity however does not necessarily preclude the role of any population structure, and demonstrate how such a structure influences collective dynamics that underlie computations.

## Acknowledgements

The project was supported by the ANR project MORSE (ANR-16-CE37-0016), the CRCNS project PIND, the program “Ecoles Universitaires de Recherche” launched by the French Government and implemented by the ANR, with the reference ANR-17-EURE-0017. There are no competing interests. SO thanks Joshua Johansen and Bijan Pesaran for fruitful discussions.

## Author contributions

A.D., A.V. and S.O. designed the study. A.D., A.V., S.O. developed the training and analysis pipelines. A.D., A.V., M.B., F.M., S.O. performed research and contributed to writing the manuscript.

## Code availability

Code will be made available on Github upon publication.

## Data availability

Trained models will be made available on Github upon publication.

## 4 Methods

### 4.1 Recurrent Neural Networks

We considered networks of *N* rate units that evolve over time according to

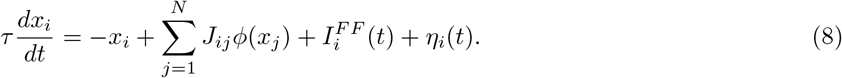

Here *x*_*i*_ represents the *activation* or total current received by the *i*-th unit, and *ϕ*(*x*_*i*_) = tanh(*x*_*i*_) is its firing rate. Moreover, each neuron received a feed-forward input 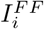 and an independent white-noise input *η*_*i*_(*t*) specified below.

The recurrent connectivity is set by the connectivity matrix ***J*** = {*J*_*ij*_}_*i,j*=1…*N*_. For full-rank networks, the coefficients *J*_*ij*_ were treated as independent parameters. For low-rank networks ***J*** was constrained to be of rank *R*, and parametrized as

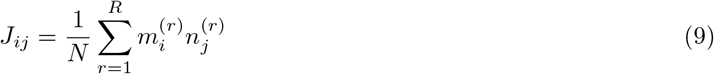

i.e. ***J*** was a sum of *R* outer-products of vectors 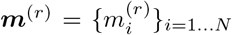 and 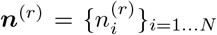. Throughout the text, we refer to the vectors ***m***^(*r*)^ and ***n***^(*r*)^ as the *connectivity vectors*, with ***m***^(*r*)^ the *r*-th output vector, and ***n***^(*r*)^ the *r*-th input-selection vector. Without loss of generality, we will assume that all the output vectors (and respectively all the input-selection vectors) are mutually orthogonal. Such a representation is uniquely defined by the singular-value decomposition of ***J*** by taking **m**^(*r*)^ to be the left singular vectors, and **n**^(*r*)^ the right singular vectors multiplied by the corresponding singular values.

The feedforward inputs 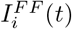 were generated by *N*_*in*_ temporally-varying scalar stimuli *u*_*s*_(*t*), each fed into the unit *i* through a set of weights 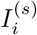:

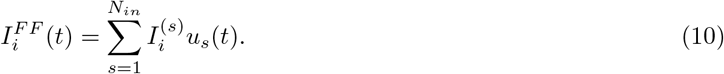

We refer to 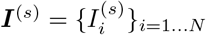 as the *s*-th *input vector*.

The output of the network was defined by a readout value

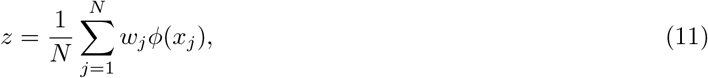

where ***w*** = {*w*_*i*_}_*i*=1…*N*_ is the *readout vector*.

The time constant of neurons was *τ* = 100ms. For simulation and training, equation (8) was discretized using Euler’s method with a time step Δ*t* = 20ms. The white noise *η*_*i*_ was simulated by drawing at each time step a random number from a centered Gaussian distribution of standard deviation 0.05.

For any pair of *N*-dimensional vectors ***a*** and ***b***, the *overlap σ*_*ab*_ was defined as the empirical covariance of their entries:

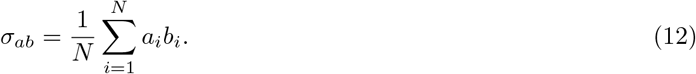

### 4.2 Network training procedure

We used backpropagation through time [Werbos, 1990] to train networks to minimize loss functions corresponding to specific tasks. For each task (see details below), we specified the temporal structure of trials and the desired mapping from combinations of stimulus inputs to target readouts 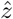, and then stochastically generated trials. We minimized the mean squared error loss function

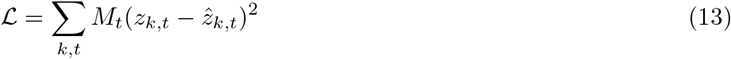

where *z*_*k,t*_ and 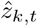 are respectively the actual, and the target readout values and the indices *k, t* respectively run over trials and time steps. The terms *M*_*t*_ are {0, 1} masks that were non-zero only during a decision period at the end of each trial, when the readouts were required to match their target values. For each task we also define a performance measure called accuracy, defined as the percentage of t t trials for which the network output has the same sign as the expected output (i.e. 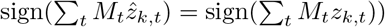)

For full-rank networks (Figs. 1,5) the gradients were computed with respect to individual entries *J*_*ij*_ of the connectivity matrix. For results on full-rank networks in Fig. 1 (left column) and Sup. Fig. S3, matrices ***J*** were initialized with random independent Gaussian weights of mean 0 and variance *ρ* = 1/*N*. For the supplementary results on Sup. Fig. S3, we also trained networks whose weights were initialized with a variance *ρ* = 0.1/*N*, since these tend to be approximated more easily by low-rank networks [Schuessler et al., 2020b].

For low-rank networks, we specifically looked for solutions in the sub-space of connectivity matrices with rank *R*. The loss functions were therefore minimized by computing gradients with respect to the elements of connectivity vectors {***m***^(*r*)^}_*r*=1…*R*_, {***n***^(*r*)^}_*r*=1…*R*_. Unless specified otherwise in the description of individual tasks, we did not train the entries of input vectors 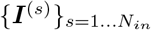 and the readout vectors {**w**} but only an overall amplitude factor for each and readout vector. All vectors were initialized with their entries drawn from Gaussian distributions with zero mean and unit standard deviation, except for the readout vector, for which the standard deviation was 4. The initial network state at the beginning of each trial was always set to **0**. We used the ADAM optimizer [Kingma and Ba, 2014] in pytorch [Paszke et al., 2017] with the decay rates of the First and second moments of 0.9 and 0.999, and learning rates between 10^−3^ and 10^−2^.

To identify networks of minimal rank that performed each task, the number of pairs of connectivity vectors *R* was treated as a hyper-parameter. We first trained full rank networks (*R* = *N*) and determined the accuracy with which they solved the task. We then started training rank *R* = 5 networks, and progressively decreased the rank until there was a sharp decrease in accuracy (Sup. Fig. S2). The minimal rank *R*\* was defined for each task such that the accuracy at *R*\* was at least of 95%.

To ease the clustering and resampling procedure, and approach mean-field solutions, we trained large networks (of sizes 512 neurons for the networks of figures 1 and 2, 4096 neurons for the context-dependent DM and DMS task networks of Figures 3 and 4, and 1024 neurons in figure 5).

### 4.3 Definition of individual tasks

#### 4.3.1 Perceptual decision making (DM) task

##### Trial structure

A fixation epoch of duration *T*_*fix*_ = 100ms was followed by a stimulation epoch of duration *T*_*stim*_ = 800ms, a delay epoch of duration *T*_*delay*_ = 100ms and a decision epoch of duration *T*_*decision*_ = 20ms.

##### Inputs and outputs

The feed-forward input to neuron *i* on trial *k* was

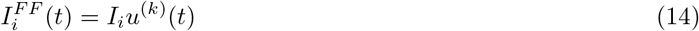

where, during the stimulation period, 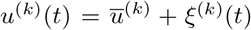, with *ξ*^(*k*)^(*t*) a zero-mean Gaussian white noise with standard deviation *σ*_*u*_ = 0.1. The mean stimulus 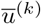 was drawn uniformly from ±0.1 × {1, 2, 4} on each trial. The elements *I*_*i*_ of the input vector were generated from a Gaussian distribution with zero mean and unit standard deviation, and fixed during training. During the decision epoch, the output *z* was evaluated through a readout vector ***w*** = {*w*_*i*_}_*i*_=1…*N*, the elements *w*_*i*_ of which were generated from a Gaussian distribution with zero mean and standard deviation of 4, and fixed during the training. On trial *k*, the target output value 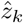 in the loss function (Eq. (13)) was defined as the sign of the mean input 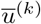.

#### 4.3.2 Parametric working memory (WM) task

##### Trial structure

A fixation epoch of duration *T*_*fix*_ = 100ms was followed by a first stimulation epoch of duration *T*_*stim*1_ = 100ms, a delay epoch of duration *T*_*delay*_ drawn from a uniform distribution between 500 and 2000ms, a second stimulation epoch of duration *T*_*stim*2_ = 100ms and a decision epoch of duration *T*_*decision*_ = 100ms.

##### Inputs and outputs

The feed-forward input to neuron *i* on trial *k* was

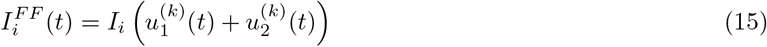

where 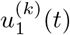 and 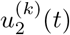 were non-zero during the first and second stimulation epochs respectively. On trial *k* and during the corresponding stimulation epoch, the values of these inputs were 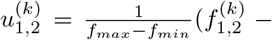 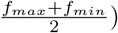, with 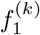 and 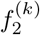 drawn uniformly from {10, 11, … , 34}, and *f*_*min*_ = 10 and *f*_*max*_ = 34. The elements *I*_*i*_ of the input vector were generated from a Gaussian distribution with zero mean and unit standard deviation, and fixed during the training.

During the decision epoch, the output *z* was evaluated through a readout vector ***w*** = {*w*_*i*_}_*i*=1…*N*_, the elements *w*_*i*_ of which were generated from a Gaussian distribution with zero mean and standard deviation of 4, and fixed during the training. On trial *k*, the target output value 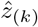 in the loss function (Eq. (13)) was defined as 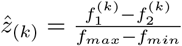.

#### 4.3.3 Context-dependent decision making (CDM) task

##### Trial structure

A fixation epoch of duration *T*_*fix*_ = 100ms was followed by a first context-only epoch of duration *T*_*ctxt*1_ = 0ms for figure 1 and 350*s* for figure 3 and 4 plots, followed by a stimulation epoch of duration *T*_*stim*_ = 800ms, a second context-only epoch of *T*_*ctxt*2_ = 500ms, and a decision epoch of *T*_*decision*_ = 20ms.

##### Stimuli and outputs

The feed-forward input to neuron *i* on trial *k* was

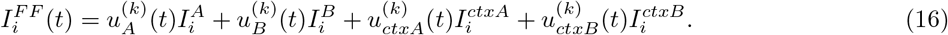

Here 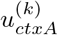 and 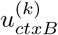 correspond to contextual cues. On each trial, during the context-only and the stimulation epochs, one of the two cues took a value +0.1 (or +0.5 for figures 3 and 4), while the other was 0. The inputs 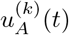 and 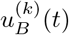 represent two sensory features of the stimulus. They were non-zero only during the stimulation epoch, and took the same form as in the perceptual decision-making task, with means 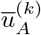 and 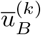, and fluctuatingparts 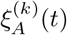 and 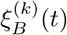 drawn independently for each feature, on each trial. The elements of the input vectors were generated from a Gaussian distribution with zero mean and unit standard deviation on both populations. For the networks presented in the main text, input vectors were trained, while for the networks reported in supplementary section S2.3 all the input vectors were fixed throughout training. During the decision epoch, on trial *k* the target 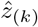 in the loss function (Eq. (13)) was defined as the sign of the mean 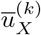 of feature *X* = *A* or *B* for which the contextual cue was activated, i. e. 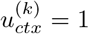. The readout vector was fixed throughout training.

#### 4.3.4 Multi-sensory decision making (MDM) task

##### Trial structure

A fixation epoch of duration *T*_*fix*_ = 100ms was followed by a context-only period of duration *T*_*ctx*_ = 350ms, a stimulation epoch of duration *T*_*stim*_ = 800ms, a delay epoch of duration *T*_*delay*_ = 300ms and a decision epoch of duration *T*_*decision*_ = 20ms.

##### Inputs and outputs

The feed-forward input to neuron *i* on trial *k* had the same structure as for the context-dependent decision-making task, and was given by:

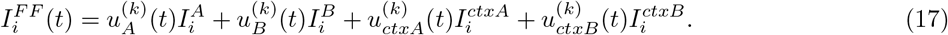

where the two stimulus inputs 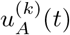 and 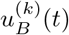 represent two sensory modalities, and 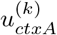 and 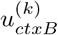 are contextual inputs. In this task, the contextual inputs were irrelevant for the output, and we included them as a control. The inputs 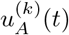 and 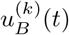 were generated as for the CDM task, with the difference that on each trial the two inputs provided congruent evidence for the output, i.e. their means were of the same sign.

Specifically in each trial a sign *s*_*k*_ ∈ {−1, 1} is generated randomly, as well as a modality that can be A, B, or AB. Then if the modality is A or AB, a mean 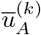 is chosen from 0.1 × *s*_k_ × {1, 2, 4} and the signal 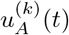 during the stimulation period is set to that mean plus a gaussian white noise as in the perceptual decision making task. A contextual input signal 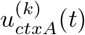 is set to 0.1 from the beginning of the contextual period to the end of the trial. If the modality is B, then the signal 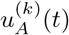 is only equal to the zero-centered gaussian white noise. The signals 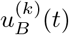 and 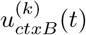 are set in a similar manner. During the decision epoch, the target 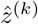 is the underlying common sign *s*_*k*_.

The networks received input signals through input vectors ***I***^*A*^, ***I***^*B*^, ***I***^*ctxA*^ and ***I***^*ctxB*^ which were trained, and output was read through a readout vector ***w*** which was fixed throughout training.

#### 4.3.5 Delayed-match-to-sample task

##### Trial structure

A fixation epoch of duration *T*_*fix*_ = 100ms was followed by a first stimulus epoch of duration *T*_*stim*1_ = 500ms, a delay epoch of a duration drawn uniformly between 500ms and 3000ms, a second stimulus epoch of duration *T*_*stim*2_ = 500ms, and a decision epoch of duration *T*_*decision*_ = 1000ms.

##### Stimuli and outputs

During each stimulus epoch, the network received one of two stimuli *A* or *B*, which were randomly and independently chosen on each trial and stimulus epoch. These two stimuli were represented by two input vectors ***I***^*A*^ and ***I***^*B*^, so that the feed-forward input to neuron *i* on trial *k* was:

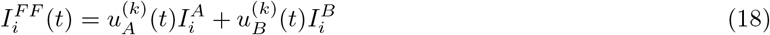

where the inputs 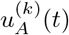 and 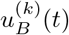 were non-zero only when stimuli *A* or *B* are respectively received, in which case they were equal to one.

During the decision epoch, the target output value 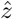 in the loss function (Eq. (13)) was equal to +1 if the same stimulus was received in both stimulation epochs and −1 otherwise.

### 4.4 Regression analyses and selectivity space

We used multivariate linear regression to predict time-averaged neural firing rates *r*_*i*_ = *ϕ*(*x*_*i*_) from task variables, using a linear model :

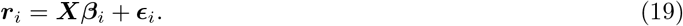

Here ***r***_*i*_ = {*r*_*i,*1_, … , *r*_*i,K*_} is a vector containing the time-averaged firing rates of neuron *i* in trials 1 to *K*, ***X*** is the design matrix where rows correspond to different trials and columns correspond to *D* task variables such as stimulus, context and decision in each condition (defined below for each task), ***β***_*i*_ is a *D*-by-1 vector of regression coefficients, and ***ϵ***_*i*_ is a *K*-by-1 vector of residuals.

The regression coefficients defined the *selectivity space* (Fig. 1a-d) of dimension *D* where each axis corresponded to the regression coefficient with respect to one task variable, and each neuron was represented as point ***β***_*i*_. The choice of task variables and window of time-averaging of firing rates depended on the task:

- For the DM task, two regressions were performed on different time windows, leading to *D* = 2 two coefficients per neuron: a regression of average firing rate during the first 100ms of stimulation period against mean stimulus which defined the coefficient 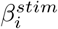 and a regression of average firing rate during the decision period against network choice which defined the coefficient 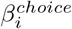. This was done to avoid ill-conditioning due to correlations between choice and stimulus.
- For the WM task, the mean firing rate during the decision period was regressed against both *f*_1_ and *f*_2_, leading to *D* = 2 two coefficients per neuron.
- For the MDM task and the CDM task, the average firing rate during the stimulation period was regressed against both mean stimulus features 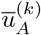 and 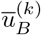 and both contextual input signals 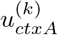 and 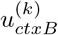, leading to *D* = 4 coefficients per neuron, 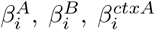 and 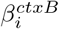. In Fig. 5, the selectivity to context was characterized by a single regression coefficient 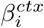 obtained by regressing the absolute value of the firing rate |*r*_*i*_|, averaged over the pre-stimulus period where only the contextual cues are non-zeros, against a regressor *X* that takes the value +1 in context A and 1 in context B. The context selectivity is extracted through the linear model for *K* trials

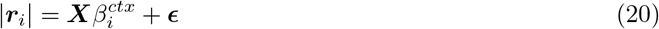 In order to characterize the changes in selectivity with context, we substracted the pre-stimulus firing rate to the firing rate averaged over the first 100ms of stimulus presentation, and regressed this quantity against 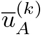 and 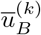 separately in each context to obtain the regression coefficients 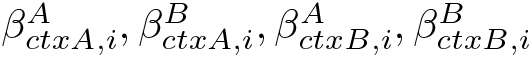. The change in selectivity is then given by

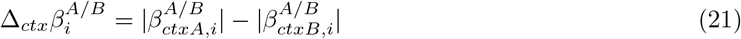 In Fig. 5 the analysis is presented for feature A, similar results are obtained for feature B (not shown).
- For the DMS task, the average firing rate during the decision period was regressed against both first and second stimulus identity (with *X*_*k,s*_ = 1 if stimulus *s* is A in trial *k*, 0 otherwise, *s* ∈ {0, 1}), leading to *D* = 2 regression coefficients per neuron.

### 4.5 Connectivity space

For a low-rank network, the connectivity is specified by 2*R* + *N*_*in*_ + 1 parameters for each neuron, corresponding to its entries 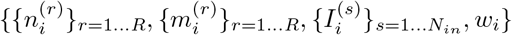 on the input, connectivity and output vectors. The connectivity of each neuron can therefore be represented as a point in a space of dimension 2*R* + *N*_*in*_ + 1 that we term *connectivity space*. For each network, the distribution of points in this space is analysed for randomness in Figure 1, and used in the resampling procedures of figures 1, 2 and 3. Our mean-field theory shows that in the limit of large networks, the distribution of points in this space determines the low-dimensional latent dynamics of the network (see Section 4.8.2).

### 4.6 ePairs analysis

To statistically assess the presence of non-random population structure in the selectivity and connectivity spaces of trained networks, we implemented a version of the ePAIRS statistical test [Hirokawa et al., 2019], which is itself derived from the PAIRS test developed in Raposo et al. [2014]. We consider a point cloud **X** = (*X*_*ij*_)_1≤*i*≤*N*,1≤*j*≤*d*_, where the rows **x**_*i*_ corresponds to different points (here neurons) and columns correspond to different axes of the considered space (regression coefficients to different variables in the selectivity space, entries of different input, connectivity nd readout vectors in the connectivity space), which is centered by removing the mean (so that for each*j*, Σ_*i*_*X*_*ij*_. The ePAIRS test examines the directional distribution of points, i.e. the empirical distribution of **x**_*i*_/∥**x**_**i**_∥, and determines whether it is statistically distinguishable from the null distribution generated by a multi-variate Gaussian with a covariance matrix identical to the covariance of **X**. A significant outcome indicates of the ePairs test that the empirical distribution presents multiple “preferred” directions incompatible with a Gaussian.

More specifically, the analysis proceeds as follows:

1. For each point **x**_*i*_, we determine its *l* nearest neighbors in terms of the cosine metric (*ie.* the *l* points for which 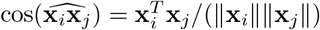 are the highest, *l* being a hyperparameter set to 3 in our case).
2. For each neuron, we compute the mean angle *α*_*i*_ with its *l* nearest neighbors, defining an empirical distribution 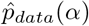.
3. To generate the corresponding null distribution, a multivariate Gaussian distribution 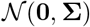 is fit to the cloud of points **X**, with **Σ** the empirical covariance of **X**, computed as 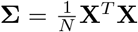. Then the steps 1-2 are applied on 500 samples of the multivariate Gaussian with the same number *N* of data points to compute a Monte-Carlo null distribution 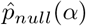.
4. Finally, the difference between the data and the null distributions is assessed using a Wilcoxon’s rank-sum test, giving a p-value, and the effect size *c* is computed as

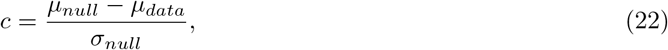

where *μ* and *σ* represent the means and standard deviations of 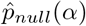 and 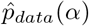. An effect size *c* > 0 indicates that angles between neighbors are smaller in the data than in the resampled point clouds, meaning that points are more highly clustered than expected. On the contrary, *c* < 0 indicates that points are more regularly dispersed than expected from random.

### 4.7 Resampling and clustering trained networks

For a given trained network, we first fitted a single multivariate Gaussian to its connectivity distribution by computing the empirical covariance matrix (matrix of size (*N*_*in*_ + 2*R* + 1)^2^). We then generated networks by resampling connectivity parameters from this distribution, and examined their performance (Fig. 1i and Sup. Fig. S3). In all trained networks we examined, the empirical means were close to zero, and we neglected them.

For the CDM and DMS tasks, we performed a clustering analysis in the connectivity space by fitting multivariate mixtures of Gaussians with an increasing number of clusters, and by resampling from the obtained distributions until we found networks that were able to optimally perform the task, as defined by an accuracy higher than 95% for at least 95% of the sampled networks. We used variational inference with a gaussian prior for the mean with a precision equal to 10^5^ to enforce a zero-mean constraint for all components of the mixtures, and a Dirichlet process prior for the weights with concentration 1 divided by number of components, using the model `BayesianGaussianMixture` of the package scikit-learn [Pedregosa et al., 2011].

Since the inference and resampling processes are susceptible to finite-size fluctuations, for the DMS task in Fig. 3 we complemented the clustering with some retraining of the covariance matrices found for each component. For this we developed a class of Gaussian mixture, low-rank RNNs, in which the covariance structure of each population is trainable. Directly training the covariance matrices is difficult given that they need to be symmetric definite positive; we therefore used a trick akin to the reparametrization trick used in variational auto-encoders [Kingma and Welling, 2013]: the set of input, connectivity and readout vectors were defined as a linear transformation of a basis of i.i.d. normal vectors, such that for any connectivity vector ***a***:

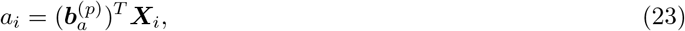

where *p* is the population index of neuron *i* (sampled from a categorical distribution with weights {*α*_*p*_}_*p*=1…*P*_ derived by the variational inference), 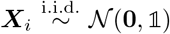 are random normal vectors of dimension *N*_*in*_ + 2*R* + 1, and the vectors 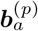 correspond to the rows of the Cholesky factorization of the covariance matrix (such that 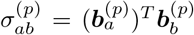 see SI section S1 for more details). We then trained the vectors 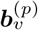, with the population indices being sampled only once, and the ***X***_*i*_ being resampled at each training epoch.

### 4.8 Analysis of latent dynamics in low-rank networks

Here we provide an overview of the reduction of low-rank networks to low-dimensional latent dynamics. A more complete derivation can be found in [Beiran et al., 2021]. For simplicity, we consider the noise free case (*η*_*i*_(*t*) = 0 in Eq. (8)), and we assume the initial condition *x*_*i*_ = 0 at *t* = 0 for all *i* = 1 … *N*.

#### 4.8.1 Low-dimensional dynamics

The dynamics defined by Eq. (8) can be represented as a trajectory in the *N*-dimensional state space in which each axis corresponds to the activation *x*_*i*_ of unit *i*. When the connectivity is constrained to be of low rank, the dynamics are restricted to a low-dimensional subspace of this state-space [Mastrogiuseppe and Ostojic, 2018]. Indeed, inserting Eqs. (9) and (10) into Eq. (8), leads to

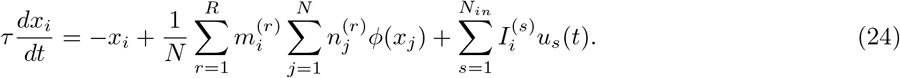

At any time *t*, the right-hand-side is confined to the linear subspace spanned by the vectors {***m***^(*r*)^}_*r*=1…*R*_ and 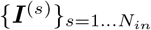. Since we assumed *x*_*i*_ = 0 at *t* = 0, the dynamics of ***x***(***t***) = {*x*_*i*_(*y*)}_*i*=1…*N*_ remain in that subspace for all *t*. The activation vector ***x*** can therefore be expressed in terms of *R* internal collective variables *κ*_*r*_, and *N*_*in*_ external collective variables *υ*_*s*_:

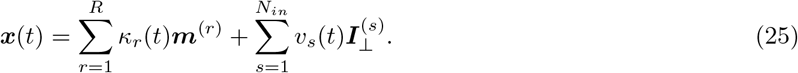

The first term on the right-hand side in Eq. (25) represents the component of the activity on the *recurrent space* [Wang et al., 2018; Remington et al., 2018] defined as the sub-space spanned by the output connectivity vectors {***m***^(*r*)^}_*r*=1…*R*_. The corresponding internal collective variables *κ*_*r*_ are defined as projections of the activation vector ***x*** on the ***m***^(*r*)^:

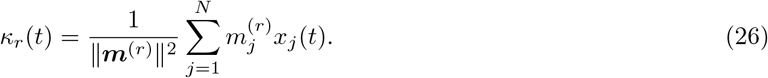

The second term on the right-hand side in Eq. (25) represents the component of the activity on the *input space* defined as the sub-space spanned by 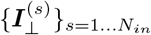, the set of input vectors orthogonalized with respect to the recurrent sub-space. The corresponding external collective variables *υ*_*s*_ are defined as projections of the activation vector ***x*** on the 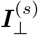:

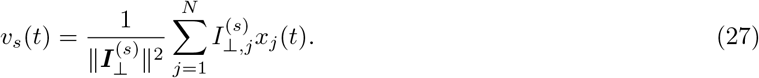

The dimensionality of the dynamics in state space is thus given by the sum of the dimension *R* of the recurrent sub-space, i.e. the rank of the connectivity, and the dimensionality *N*_*in*_ of the input space.

The dynamics of the internal variables *κ*_*r*_ are obtained by projecting Eq. (8) onto the output connectivity vectors ***m***^(*r*)^:

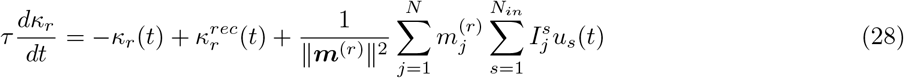

where 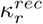 represents the recurrent input to the *r*-th collective variable, defined as the projection of the firing rate vector *ϕ*(***x***) onto the input-selection vector ***n***^(*r*)^:

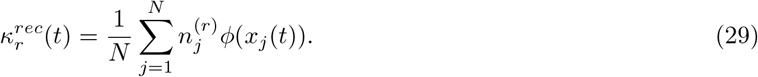

Inserting Eq. (25) into 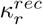 leads to a closed set of equations for the *κ*_*r*_:

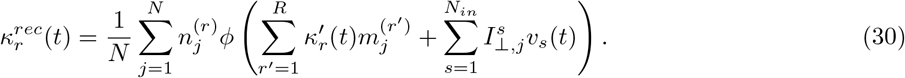

The dynamics of the external variables *υ*_*s*_ is obtained by projecting Eq. (8) onto the orthogonalized input vectors 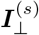. They are given by external inputs *u*_*s*_(*t*) filtered by the single neurons time constant *τ*

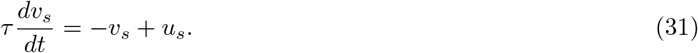

Throughout the main text, we assume for simplicity that the stimuli *u*_*s*_ vary on a timescale slower than *τ*, and replace *υ*_*s*_ with *u*_*s*_. We also assume throughout the main text that input vectors are orthogonal to the output connectivity vectors, ie. 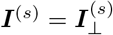 for all *s*. Hence the third term on the r.h.s. of equation (28) equals zero. Using Eq. (25), the readout value *z* can be expressed in terms of the collective variables as

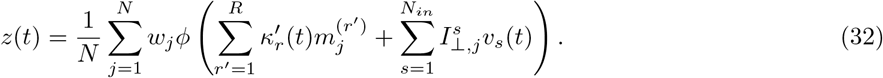

#### 4.8.2 Connectivity space and mean-field limit

The dynamics of the collective variables are fundamentally determined by the components of connectivity and input vectors through Eq. (30). Neuron *i* is therefore characterized by the 2*R* + *N*_*in*_ + 1 parameters

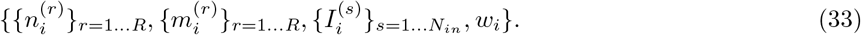

Each neuron can thus be represented as a point in the *connectivity space* of dimension 2*R* + *N*_*in*_ + 1, and the connectivity of the full network can therefore be described as a set of *N* points in this space. Note that the right-hand-side of Eq. (30) consists of a sum of *N* terms, where the term *j* contains only the connectivity parameters of neuron *j*. The connectivity parameters of different neurons therefore do not interact in 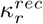, so that the r.h.s of Eq. (30) can be interpreted as an average over the set of points corresponding to all neurons in the connectivity space.

Our main assumption will be that in the limit of large networks (*N* → ∞), the set of points in the connectivity space is described by a probability distribution 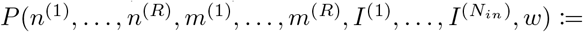 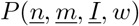. In this mean-field limit, the r.h.s. of Eq. (30) becomes:

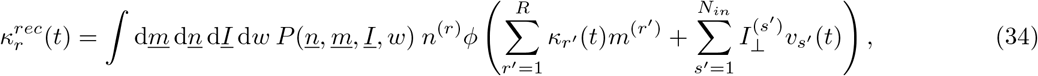

where we have used the shorthand 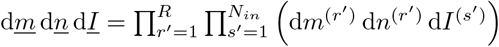. The collective dynamics are therefore fully specified by the single-neuron distribution of connectivity parameters. Once this distribution is specified, any network generated by sampling from it will have identical collective dynamics in the limit of a large number of neurons.

The joint distribution of connectivity parameters *P* (<u>*n*</u>, <u>*m*</u>, <u>*I*</u>, *w*) also determines the values of the readout:

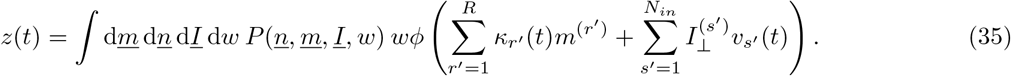

#### 4.8.3 Statistics of connectivity and sub-populations

To approximate any arbitrary joint distributions of connectivity parameters *P* (<u>*n*</u>, <u>*m*</u>, <u>*I*</u>, <u>*w*</u>), we used multivariate Gaussian mixture models (GMMs). This choice was based on the following considerations: (i) GMMs are able to approximate an arbitrary multi-variate distribution [Kostantinos, 2000]; (ii) model parameters can be easily inferred from data using GMM clustering; (iii) GMMs afford a natural interpretation in terms of sub-populations (iv) GMMs allow for a mathematically tractable and transparent analysis of the dynamics as shown below [Beiran et al., 2021].

In a multivariate Gaussian mixture model, every neuron belongs to one of *P* sub-populations. For a neuron in sub-population *p*, the set of parameters 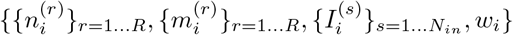 is generated from a multivariate Gaussian distribution with mean ***μ***_*p*_ and covariance **Σ**_*p*_, where ***μ***_*p*_ is a vector of size 2*R* + *N*_*in*_ + 1, and **Σ**_*p*_ is a covariance matrix of size (2*R* + *N*_*in*_ + 1)^2^. The full distribution of connectivity parameters is therefore given by

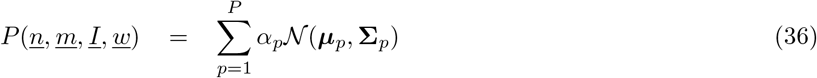

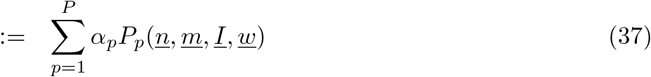

where the coefficients *α*_*p*_ define the fraction of neurons belonging to each sub-population.

Each sub-population directly corresponds to a Gaussian cluster of points in the connectivity space. The vector ***μ***_*p*_ determines the center of the *p*-th cluster, while the covariance matrix **Σ**_*p*_ determines its shape and orientation. For a neuron *i* belonging to population *p*, we will write as 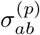 the covariance between two connectivity parameters *a* and *b*, with *a, b* ∈ {{*n*^(*r*)^}_*r*=1…*R*_, {*m*^(*r*)^}_*r*=1…*R*_, 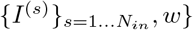. Note that because the output vectors ***m***^(*r*)^ (resp. input-selection vectors ***n***^(*r*)^) are mutually orthogonal, the covariances between the parameters 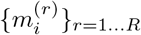 (respectively 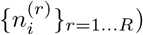 vanish.

Since every neuron belongs to a single population, the r.h.s of Eq. (30) can be split into P terms, each corresponding to an average over one population. As within each population the distribution is a joint Gaussian, Eq. (34) becomes a sum of *P* Gaussian integrals

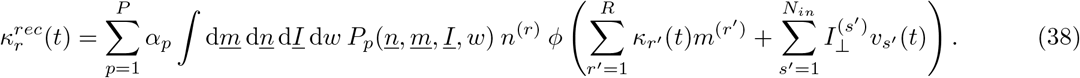

#### 4.8.4 Effective circuit description of latent dynamics

In the following, we focus on zero-mean multivariate Gaussian mixture distributions for the connectivity parameters, and input vectors orthogonal to {***m***^(*r*)^}_*r*=1…*R*_, as distributions with these assumptions were sufficient to describe trained networks. The more general case of Gaussian mixtures with non-zero means is treated in [Beiran et al., 2021]. Using Stein’s lemma for Gaussian distributions, the dynamics of the internal collective variables can be expressed as a dynamical system (see SI section S1)

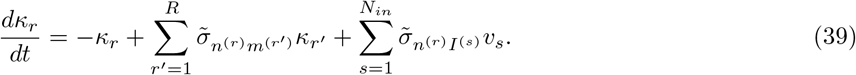

In the main text, *υ*_*s*_ were replaced by *u*_*s*_ which amounts to assume that inputs vary slowly with respect to the single neuron time constant *τ*.

In Eq. (39), 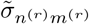 represents the effective self-feedback of the collective variable *κ*_*r*_, 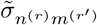 sets the interaction between the collective variables *κ*_*r*_ and *κ*_*r*′_, and 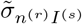 is the effective coupling between the input *u*_*s*_ and *κ*_*r*_. These effective interactions between the internal variables are given by weighted averages over populations

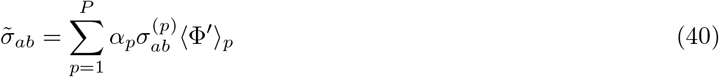

where 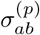 is the covariance between connectivity parameters *a* and *b* for population *p*, and ⟨Φ′⟩_*p*_ is the average gain of population *p*, defined as

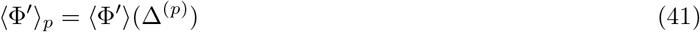

with

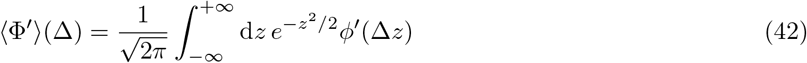

and

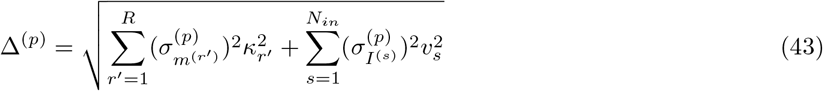

the standard deviation of activation variables in population *p*, where 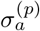 is the variance of a vector ***a*** on population *p*.

In Eq. (39), the covariances 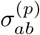 are set by the statistics of the connectivity and input vectors, but the gain factors ⟨Φ′⟩_*p*_ in general depend both on internal and external collective variables *κ*_*k*_ and *v*_*j*_. As a consequence, the dynamics in Eq. (39) is non-linear, and in fact it can be shown that given a sufficient number of sub-populations, the right-hand side in Eq. (39) can approximate any arbitrary dynamical system [Beiran et al., 2021].

In the special case of linear networks (i.e. Φ(*x*) = *x*), the gain is constant so that the effective couplings 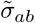 in Eq. 40 are equal to the overlaps *σ*_*ab*_ of vectors *a* and *b* over the full population, as defined in Eq. 12. The population structure therefore only plays a role for non-linear networks.

The value of the readout (Eq. (35)) can also be expressed in terms of effective interactions as

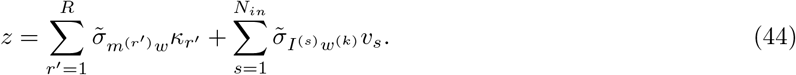

#### 4.8.5 Drivers and modulators of latent dynamics

Eq. (39) shows that feed-forward inputs to the network can have two distinct effects on the collective dynamics of internal variables *κ*_*r*_. If the input vector *I*^(*s*)^ overlaps with the *r*-th input-selection vector *n*^(*r*)^, i.e. the corresponding covariance 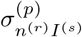 is non-zero for population *p*, the input directly drives the latent dynamics, in the sense that *v*_*s*_ acts as an effective external input to the dynamics of *κ*_*r*_ in Eq. (39).

In contrast, when all covariances between the input vector *s* and the input selection vectors are zero (i.e. 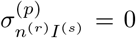 for all *r, p*), the corresponding input does not drive the latent dynamics, but can still modulate them by modifying the gain through Eq. (43) if the variance 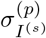 of the input on some population *p* is non-zero.

The inputs can therefore play roles of drivers and modulators of latent dynamics, depending on whether the corresponding input vectors overlap or not with the input selection vectors *n*^(*r*)^.

### 4.9 Reduced models of latent dynamics for individual tasks

#### 4.9.1 Perceptual decision making task

##### Latent dynamics and reduced model

We found that computations in the rank one, single population trained networks could be reproduced by a reduced model with two non-zero covariances *σ*_*nI*_ and *σ*_*nm*_ (Sup. Fig. S5a). For this reduced model, the dynamics of the internal collective variable is given by

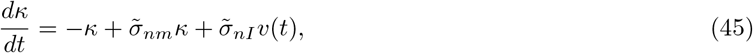

where 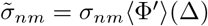 and 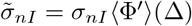 with ⟨Φ⟩(Δ) defined in Eq. (41), and the effective population variance Δ given by:

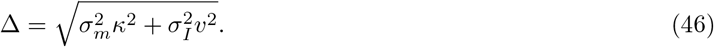

Here *υ*(*t*) corresponds to the integrated input *u*(*t*), see Eq. (31).

An analysis of nonlinear dynamics defined by Eq. (45) showed that adjusting these parameters was sufficient to implement the task, as additional parameters only modulate the overall gain (see SI section S2.1). In particular the value of *σ*_*mn*_, determines the qualitative shape of the dynamical landscape on which the internal variable evolves and sets the time scale on which it integrates inputs (see SI S2.1 for more details).

#### 4.9.2 Parametric working memory task

##### Latent dynamics and reduced model

We found that computations in the rank two, single population trained networks could be reproduced by a reduced model with four non-zero covariances 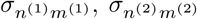 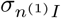 and 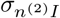 (Sup. Fig. S6a). In particular covariances 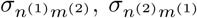 across the two vectors could be set to zero without performance impairment.

For this reduced model, the dynamics of the two internal collective variables is given by:

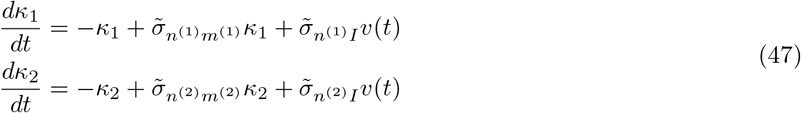

where 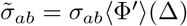, with ⟨Φ′⟩(Δ) defined in Eq. (41), and the effective noise Δ given by:

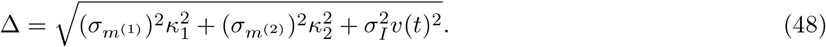

Here *υ*(*t*) corresponds to the integrated input *u*(*t*), see Eq. (31).

The two internal collective variables are therefore effectively uncoupled, and integrate the incoming feed-forward inputs at two different timescales due to different levels of positive feedback. For the first collective variable, a strong, fine-tuned positive feedback 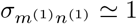 leads to an approximate line attractor along *κ*_1_ that persistently encodes the first stimulus throughout the delay and the sum of the two stimuli at the decision epoch. For the second internal variable, a weaker positive feedback 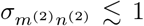 leads to a shorter timescale of a transient response to stimuli along *κ*_2_, such that the first stimulus is forgotten during the delay and that the second stimulus is represented during the decision epoch.

#### 4.9.3 Context-dependent decision making task

##### Latent dynamics and reduced model

We found that the computations in the unit rank, two populations network relied on the following conditions for the covariances in the two populations (Sup. Fig. S7a): (i) ***I***^*ctxA*^ and ***I***^*ctxB*^ were essentially orthogonal to the input-selection vector **n**, implying that 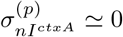 and 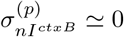 for both populations *p* = 1, 2; (ii) on each population, each of the two input-selection vectors was correlated with only one of the input-feature vectors, i.e. 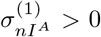 and 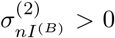, while 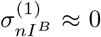 and 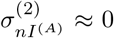; (iii) each context-cue vector had a strong variance on a different sub-population, i.e. for the first population ***I***^*ctxA*^ and ***I***^*ctxB*^ had respectively weak and strong variance (i.e. 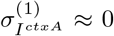 and 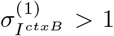), and conversely for the second population 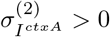 and 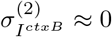.

The computation could therefore be described by a reduced model, in which the covariances 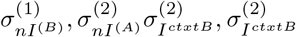 were set to zero. The dynamics of the internal variable was then given by

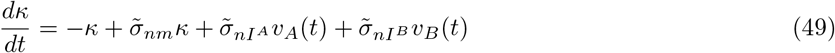

with effective couplings

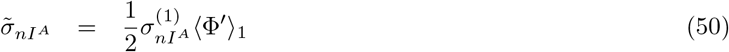

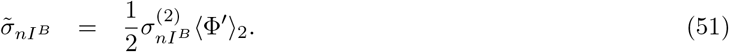

The averaged gains for each population were given by equations (42), with the standard deviations of currents onto each population

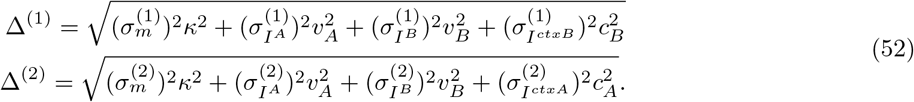

Here *υ*_*A*_(*t*) and *υ*_*B*_(*t*) correspond to the integrated inputs *u*_*A*_(*t*) and *u*_*B*_(*t*), see Eq. (31).

As for the perceptual decision making task, the value of *σ*_*mn*_, determines the qualitative shape of the dynamical landscape on which the internal variable evolves and sets the time scale on which it integrates inputs. Large values of the variances 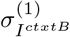 and 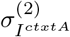 allow the contextual cues to differentially vary the gain of the two populations in the two contexts, leading to an effective gating of the inputs integrated by the internal collective variable (see SI section S2.3 for more details).

#### 4.9.4 Delayed-match-to-sample task

##### Latent dynamics and reduced model

We found that the computations in the rank two, two population network relied on the following conditions for the covariances in the two populations (Sup. Fig. S8a): (i) on one population, the two connectivity modes were coupled through 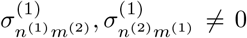, with a specific condition on their values to induce a limit cycle (that the difference 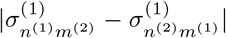 is large, see SI and [Mastrogiuseppe and Ostojic, 2018; Beiran et al., 2021]); (ii) on the other population, the covariances were in contrast set to counter-balance the first population, and cancel the rotational dynamics 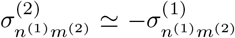 and 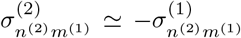; (iii) the input-selection and output vectors for the second connectivity mode on the second population had a strong overlap 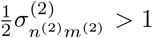 that led to strong positive feedback; (iv) the input vectors *I*^*A*^ had a strong variance on population 2, 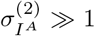 while other input sub-vectors had small variances 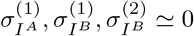.

For this reduced model, the dynamics of the two internal collective variables is given by:

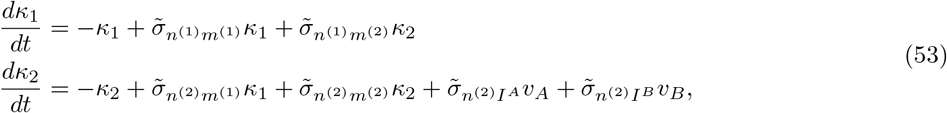

with the effective couplings mediating inputs

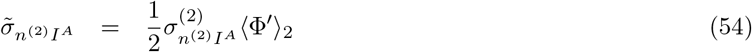

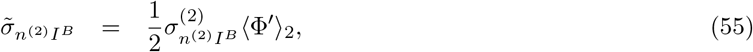

and effective couplings governing the autonomous dynamics:

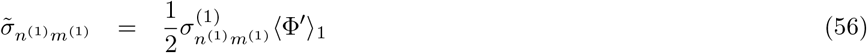

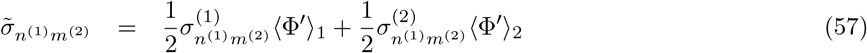

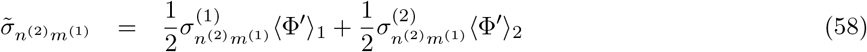

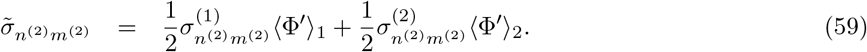

The average gains are given by equations (42), with standard deviations of currents onto each population

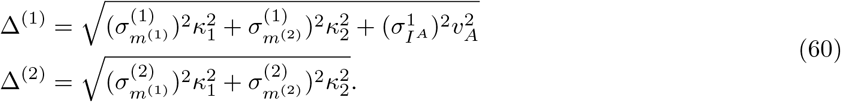

Here *υ*_*A*_(*t*) and *υ*_*B*_(*t*) correspond to the integrated inputs *u*_*A*_(*t*) and *u*_*B*_(*t*), see Eq. (31).

Conditions (i) to (iv) on the covariances allow to implement the dynamical landscape modulation of Fig. 4h (see Sup. Fig. S8d). When stimulus A is present (*u*_*A*_ = 1), the gain of population 2 is set to (Φ′)_2_ ≃ 0 because of 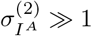 (see Eq. (60)), and the specific values of covariances for sub-vectors in population 1 induce a limit cycle (see SI section S2.4). In absence of inputs, or when input B was present, gains were approximately equal for the two populations (Sup. Fig. S8c), leading to a cancellation of the cross effective couplings 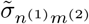 and 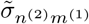, while positive feedback implemented through 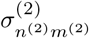 shaped a dynamical landscape with two fixed-points.

## Supplementary information

**Figure S1.**
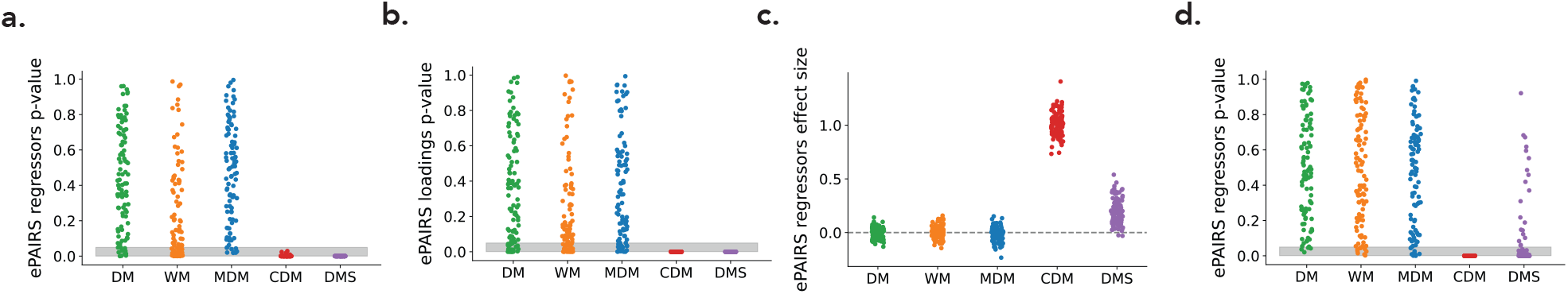
Additional ePAIRS results. (a) p-values given by the ePAIRS test on regression spaces for the full-rank networks displayed in Fig. 1d. (b) p-values given by the ePAIRS test on connectivity spaces for the low-rank networks displayed in Fig 1h. (c) ePAIRS effect size on the regression space for the same low-rank networks. (d) Associated ePAIRS p-values.

**Figure S2.**
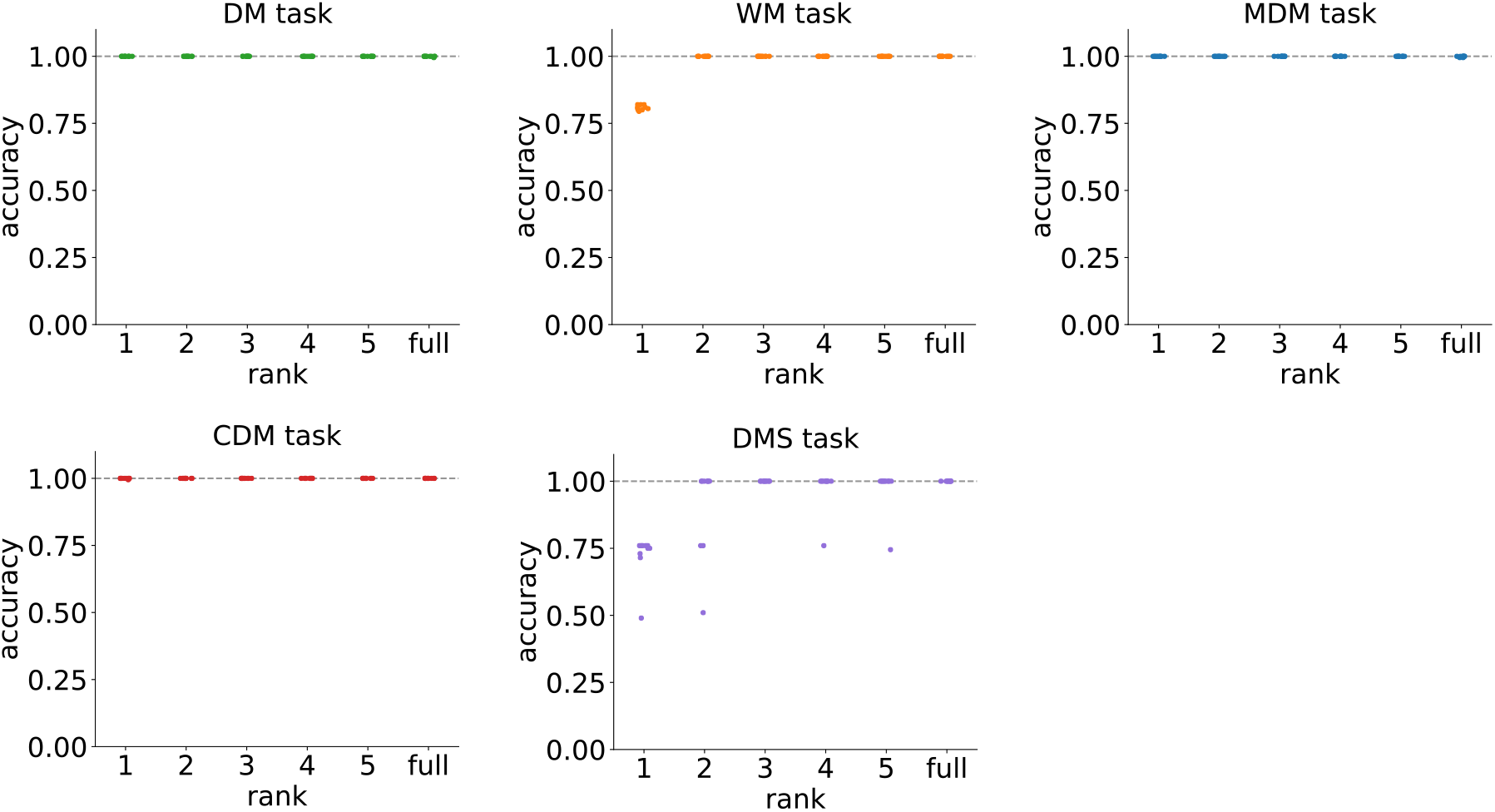
Determination of the minimal rank for each task. For each task and each rank *R* between 1 and 5, ten rank-*R* networks were trained with different random initial connectivity. For each task, a panel displays the performance of trained networks as function of their rank.

**Figure S3.**
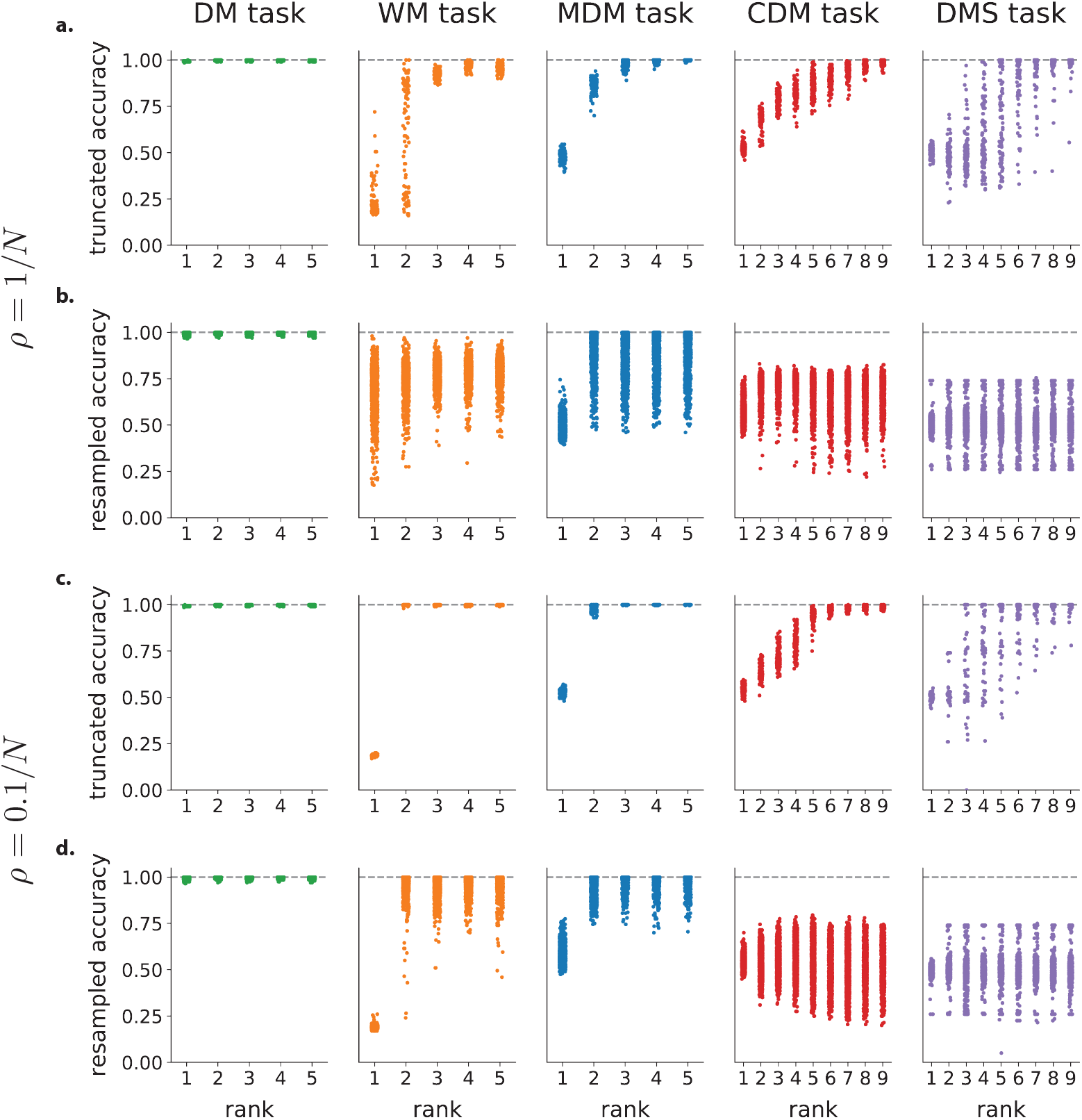
Analysis of trained full-rank networks. (a)-(b) Analysis of full-rank networks trained with initial connectivity weights of variance 1*/N* (100 networks for each task). (a) Performance of truncated-rank networks. Following [Schuessler et al., 2020b], we extract from full-rank networks the learned part of the connectivity Δ***J = J*** − ***J***_0_ defined as the difference between the final connectivity ***J*** and the initial connectivity ***J***_0_. We then truncate Δ***J*** to a given rank *via* a singular value decomposition, and add it back to ***J***_0_. For each task, a panel displays the performance of the obtained networks as function of the rank used for the truncation. (b) Resampling analysis of truncated networks. Starting from the truncated networks in (a) we fit multivariate Gaussians to the distribution of their Δ***J*** in the corresponding connectivity spaces. We then generate new networks by resampling from this distribution, as done on the trained low-rank networks for Fig. 1i-l. For each task, a panel displays the performance of the obtained resampled network as function of the rank used for the truncation. (c)-(d) Same analyses as (a)-(b) for sets of networks trained with initial connectivity weights of variance 0.1*/N* (100 networks for each task, for DMS 49/100 networks that had an accuracy < 95% after training and were ignored). Networks with weaker initial connectivity are better approximated by their resampled low-rank connectivity. This is due to the fact that larger initial connectivities induce correlations between Δ***J*** and ***J***_0_ [Schuessler et al., 2020b]. The resampling destroys both this correlation and the population structure, leading to performance impairements even when the population structure is potentially irrelevant.

**Figure S4.**
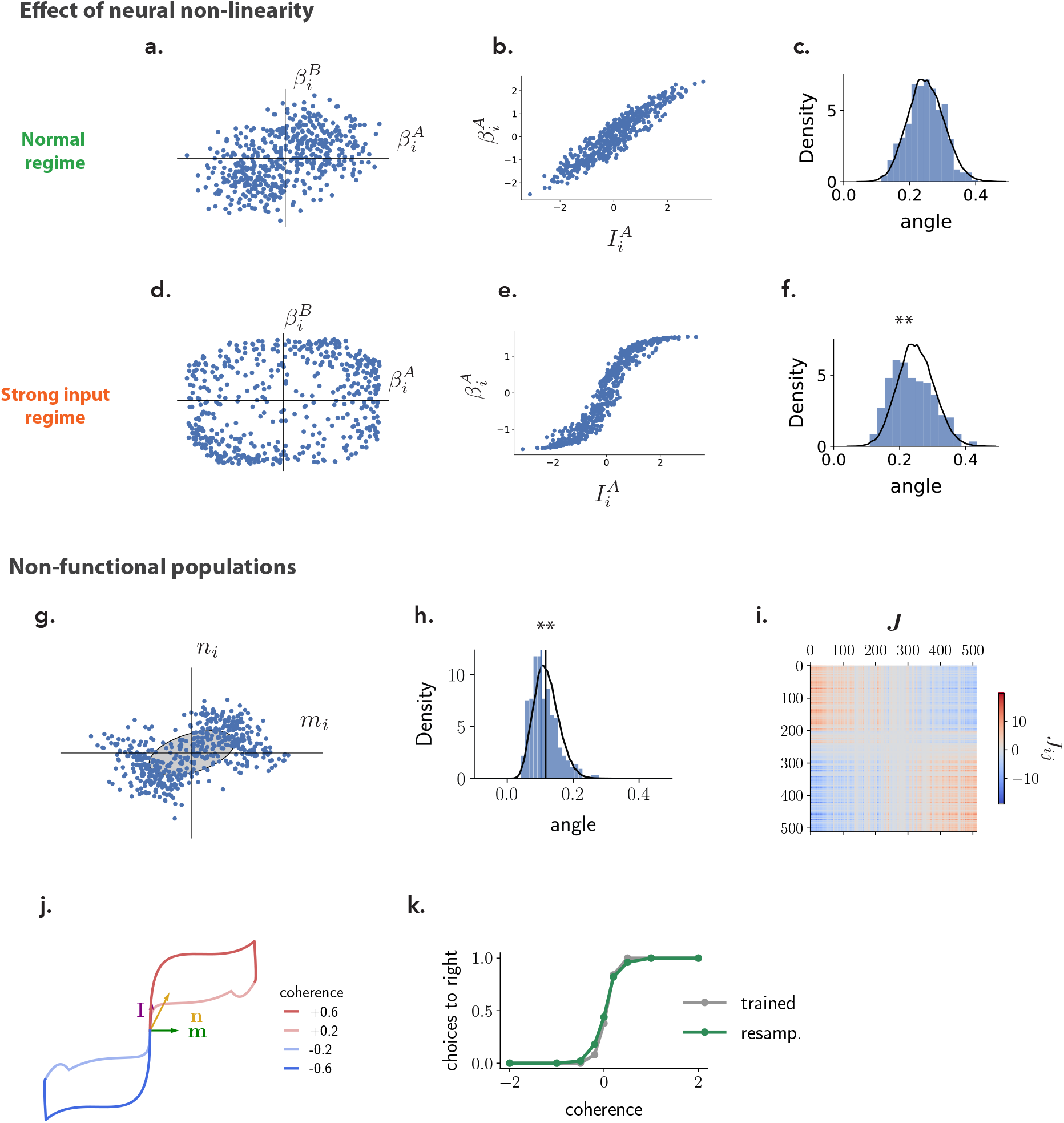
Limitations of ePAIRS analyses. This figure illustrates two situations in which the ePAIRS test leads to false positives, and identified non-random population structure that is not computationally relevant. (a)-(f) The activity in a single network trained on the MDM and used in Figure 1 was compared in two conditions: (a-c) in response to inputs of the scale used for training; (d-f) in response to inputs scaled by a factor 10. The corresponding regressor spaces were then tested for non-random population structure via the ePAIRS test. No evidence for population structure in selectivity or connectivity space is found for inputs in the range used for training (a-c). Using stronger inputs however leads to positive ePairs in the selectivity space, although the underlying connectivity is identical (d-f). (a) Slice of the selectivity space for this network representing regression coefficients for each neuron with respect to inputs A and B. (b) In this input regime, regression coefficients with respect to inputs are linear functions of the components along the corresponding input vectors (each point represents a neuron in the network). (c) an ePAIRS test on the selectivity space in that case leads to a non-significant outcome (*p* = 0.48, *c* = 0.03). (d) As in (a), for the same network, but driven with inputs 10 times larger than those used for training. The individual units are in that case driven to saturation so that the points in the selectivity space are concentrated along the borders of a square. (e) Same as (b) in the strong input regime. The relation between the original input vector and the obtained regression coefficients reflects the underlying non-linearity as neurons are driven to saturation. (f) ePairs on the square-like distribution in selectivity space shown in (d) rejects the null hypothesis for random population structure (*p* < 10^−5^, *c* = 0.3). (g)-(k) An example network trained on the Perceptual Decision Making task exhibiting spurious, computationally irrelevant population structure detected by the ePAIRS test. This network was obtained by using a different scaling of recurrent weights during training than in the rest of the manuscript. For all networks in the main text, the recurrent connectivity was defined as 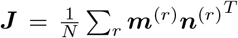 with entries of vectors ***m***^(*r*)^ and ***n***^(*r*)^ being of order 1 and the 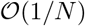 scaling of the connectivity matrix being explicitly added in the network dynamics. For this example the recurrent connectivity was defined as 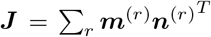 with entries of the connectivity vectors being of order 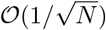and the scaling of the connectivity matrix being this time implicit, which led to different gradient descent dynamics and to significantly different solutions. Here a rank-one network of 512 neurons is shown. (g) Scatter plot of the entries of each neuron on the recurrent connectivity vectors ***m*** and ***n***, showing two clusters symmetrical with respect to the mean. Note that this cluster structure is very different from those seen in the rest of the paper, which corresponded to zero-mean clusters with different covariance matrices, while here two non-zero-mean clusters are visible. (h) The ePAIRS test detected evidence for non-random population structure on the connectivity space (which is here 4-dimensional, composed of vectors ***I***, ***n***, ***m*** and ***w***. Here, *c* = 0.35, *p* < 10^−8^). (i) The two clusters seen in the scatter plot can also be made apparent in the connectivity matrix ***J*** if its entries are properly ordered, here by ascending values of *m_i_ + n*_*i*_. (j) State-space response trajectories to different stimuli projected on the ***m*-*I*** plane are similar to those found for the network shown in Fig. 2. (k) As for the network in Fig. 2, networks resampled from a Gaussian distribution fit to the connectivity space of the trained network (black ellipse in panel a) performed equally well as the trained network, showing that the population structure found by the ePAIRS procedure was not computationally relevant, and might be an artifact of learning.

### S1 Parametrization and collective dynamics for mixture of Gaussians connectivity vectors

In this section we show how connectivity vectors with entries drawn from mixtures of multivariate Gaussians can be constructed from independent Gaussians, as mentioned in Eq. (23). We then derive the dynamics of the internal collective variables (Eq. (39)) in this setting.

We considered distributions of connectivity parameters characterized by *P* covariance matrices **Σ**_*p*_, and zero means ***μ***_*p*_ = **0**, *p* = 1, … , *P*. For a neuron *i* belonging to population *p*, each vector entry 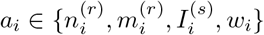 is constructed as a linear transformation of the same set of values 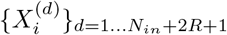

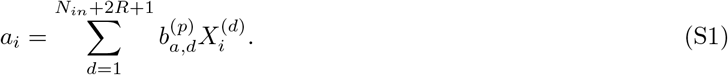

Here the 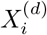 are drawn from 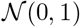, independently for each *i* and *d*. The linear coefficients 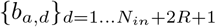 are different for each connectivity vector *a* ∈ {*n*^(*r*)^, *m*^(*r*)^, *I*^(*s*)^, *w*}, but identical across neurons within a given population. These sets of coefficients therefore determine the covariance 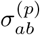 between entries of connectivity vectors within a given population *p*:

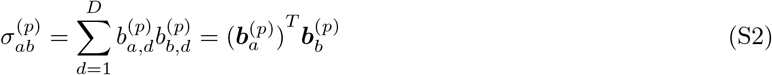

The row-vectors 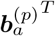 in fact correspond to the rows of the Cholesky factorization of the covariance matrix. We next turn to the derivation of Eq. (39). With the parametrization for the entries of connectivity vectors defined in Eq. (S1), the recurrent inputs to the *r*-th internal collective variable Eq. (34) can be written as

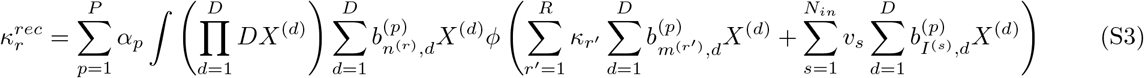

with 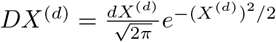. For a given *p*, we then compute each of the *D* integrals 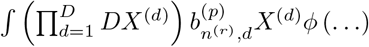 applying successively Stein’s lemma

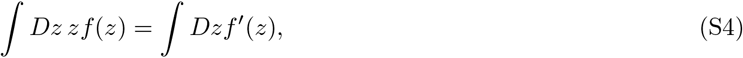

and using the fact that a sum of independent Gaussians is a Gaussian with variance given by the sum of variances, so that

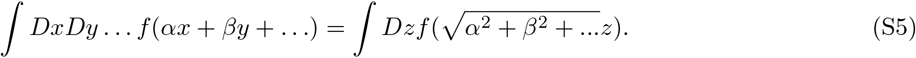

This leads to

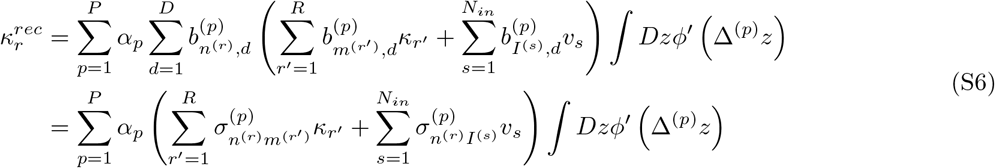

with

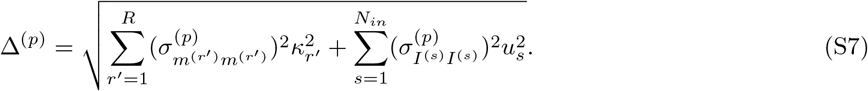

Inverting the sums on *p* and *r*′, *s* indices and assuming that input vectors ***I***^(*s*)^ are orthogonal to the output vectors {***m***^(*r*)^}_*r*=1,…,*R*_ (as in all the reduced models described in the section below), we get the compact description in terms of effective couplings for the dynamics of internal collective variables Eq. (39)

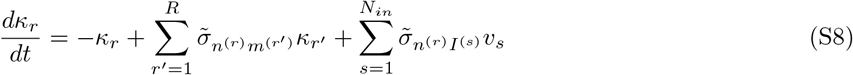

with, for any two vectors ***a***, ***b***, the effective couplings

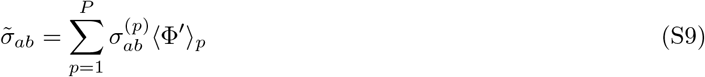

and averaged gains

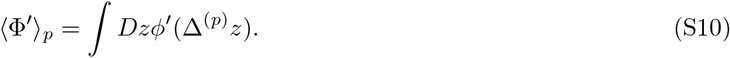

### S2 Theoretical analysis of reduced models

Here we examine reduced network models, that were minimally parametrized to solve each task by relying on the same network dynamics as the trained networks presented in the main text. The minimal parameter sets correspond to subsets of the computationally important covariances between the connectivity vectors of the trained networks. These parameters were first set by hand and then, if necessary, fine-tuned with the ADAM optimizer to solve the task with optimal accuracy. We first report on how to parametrize connectivity vectors to build these networks. We then examine the effects of these parameters on mean-field collective dynamics and show their implication in task solving.

#### S2.1 Perceptual decision-making network

The network trained on this task was of unit rank, and consisted of a single population. Such a network can be minimally parametrized using three covariances *σ*_*nm*_, *σ*_*nI*_ and *σ*_*mw*_ (Supp. Fig. S5a). This can be obtained with an input vector 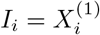 and a pair of recurrent connectivity vectors given by:

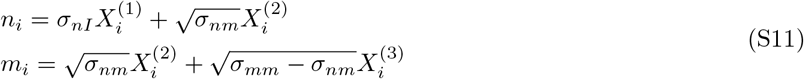

for *i* = 1, … , *N* , with 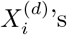 drawn independently from zero-mean Gaussian distributions of unit variance. The readout components were taken as

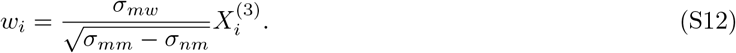

The dynamics of the single internal collective variable is then given by

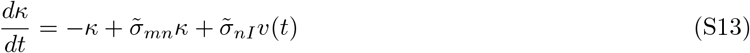

with effective couplings given by equation (S9), i.e. the covariances scaled by the global gain factor

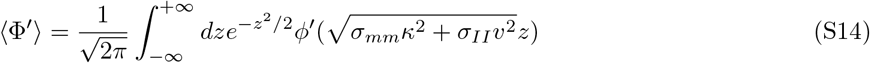

These dynamics can be graphically summarized as in Fig. 2e and lead to network dynamics that match the ones of trained networks (Fig. S5b).

The autonomous dynamics of the network is determined by the parameter *σ*_*nm*_ that controls (i) the qualitative shape of the dynamical landscape, with a transition from a single stable fixed-point (*σ*_*nm*_ < 1) to two symmetric fixed-points (*σ*_*nm*_ > 1) and (ii) the time-scale 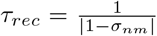 with which the network state relaxes or diverges from the initial condition ***x*** = **0** at the beginning of each trial (Fig.S5c,d, [Mastrogiuseppe and Ostojic, 2018]). The integration of the filtered input *v*(*t*) by *κ* is controlled by *σ*_*nI*_, the covariance between the input vector ***I*** and the input-selection vector ***n***. For instance for *σ*_*nI*_ = 0, *v*(*t*) is projected on a direction orthogonal to the input-selection vector and is not integrated by the recurrent activity (Supp. Fig. S5g light shade line).

Finally, the covariance *σ*_*mw*_ between the output vector ***m*** and the readout vector ***w*** controls the extent to which the readout is driven by *κ*, with no drive of the readout in case of orthogonal output and readout vectors, *σ*_*mw*_ = 0 (Supp. Fig. S5f light shade line).

The network connectivity of equation (S11), also involved the variance *σ*_*mm*_ of the connectivity vector ***m***. Changing *σ*_*mm*_ influences the autonomous dynamics of the network (Supp. Fig. S5c) by influencing the gain of the neurons (see Eq. (S14)).

For the reduced model shown in the main text, the non-zero covariances were: *σ*_*nm*_ = 1.4, *σ*_*nI*_ = 2.6 and *σ*_*mw*_ = 2.1.

#### S2.2 Parametric working-memory network

The network trained on this task was of rank two, and consisted of a single population. A minimal parametrization of this network involves six covariances 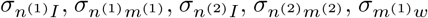 and 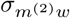 (Supp. Fig. S6a).

This can be obtained with an input vector 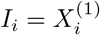 and two pairs of recurrent connectivity vectors:

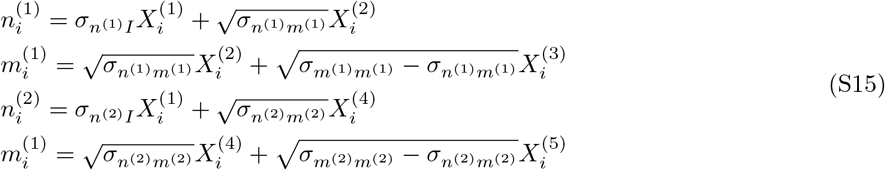

for *i* = 1, … , *N* , with 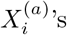 drawn from zero-mean Gaussian distributions of unit variance. The readout components were taken as

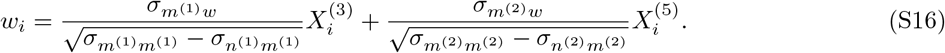

The dynamics of the two internal collective variables is then given by:

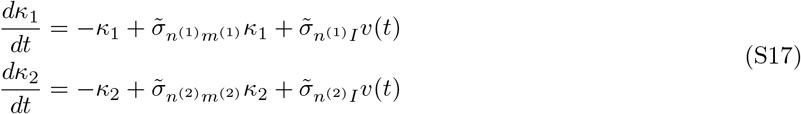

with effective couplings given by equation (S9), i.e. the covariances scaled by the global gain factor

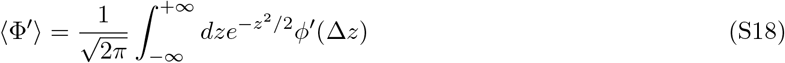

with

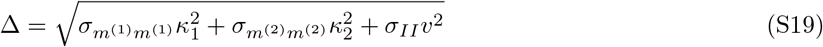

These dynamics can be graphically summarized as in Fig. 2i and reproduce the dynamics of trained networks as shown in Supp. Fig. S6b. Supp. Fig. S6c shows the dynamical phase portrait on which recurrent activity evolves. It approximates a line attractor [Seung, 1996] on the direction ***m***^(1)^ as the covariance 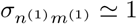 sets the network close to the bifurcation point of Supp. Fig. S5c. On the second direction ***m***^(2)^ the dynamics relax with a time scale set by the covariance 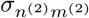. For the reduced model shown in the main text, the non-zero covariances were: 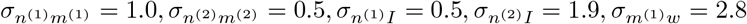 and 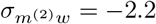.

#### S2.3 Context-dependent decision-making network

The networks trained on this task were of unit rank and consisted of either two or three populations depending on the training procedure (see methods section 4.3.3, supplementary section S3 and Sup. Fig. S9).

##### Two-population network

Such a network can be minimally parametrized using 4 non-zero covariances on each population. This can be obtained with the two sensory input vectors generated independently 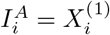, 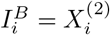, irrespective of the population structure. The connectivity vectors are structured in two sub-vectors.

For *i* in population 1:

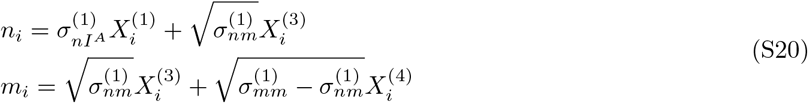

while for *i* in population 2:

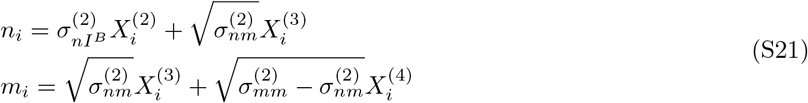

with 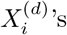 ’s drawn from i.i.d. centered Gaussian distributions of unit variance. The readout vector is taken as

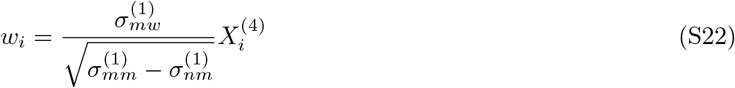

for *i* in population 1 and

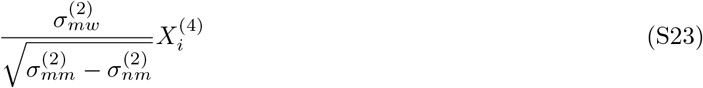

for *i* in population 2. Importantly the contextual input vectors are also structured in two sub-vectors, such that for *i* in population 1:

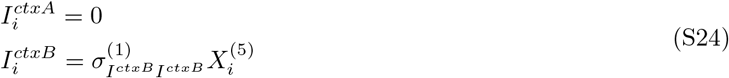

while for *i* in population 2:

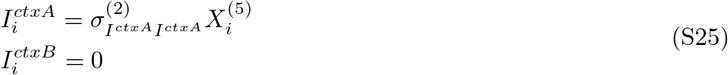

with 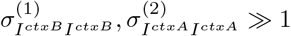.

The recurrent activity is then described by a single internal collective variable, graphically summarized in Fig. 4a:

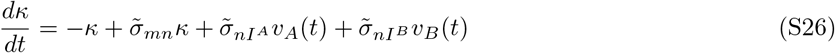

The time evolution of the internal collective variable is coupled to the two inputs through the two inputs through the two effective couplings 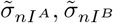 time evolution of the internal collective variable is coupled to

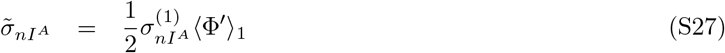

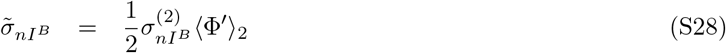

The recurrent dynamics are supported equally by the two populations:

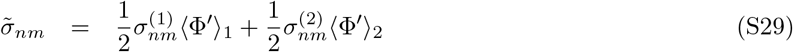

with averaged gains given by equations (S10) and standard deviations of currents onto each population

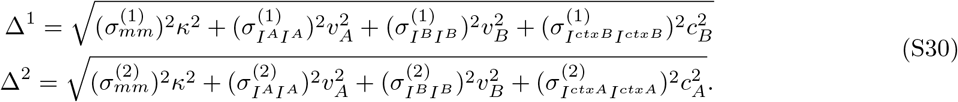

The obtained dynamics are similar to the trained networks, graphically illustrated in Fig. 4c, with contextual inputs controlling the gain of each of the two populations (Supp. Fig. S7b). This control relies on the large amplitude of the weights of contextual input vectors, 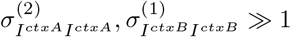, as illustrated in Supp. Fig. S7c where we show the effect of varying these parameters on the network readout. In this implementation, each of the two populations selectively integrates one of the two sensory inputs thanks to the non-zero covariances between sensory input and input-selection vectors 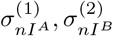, as illustrated in Supp. Fig. S7d.

The non-zero covariances for the implementation of the solution presented in the main text are given by 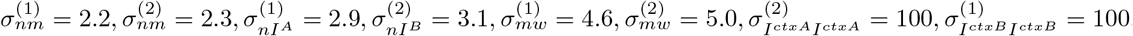.

##### Three-population network

For the context-dependent decision-making task, we also examined a network relying on three populations. In this network, two populations selectively gate inputs as in the two-population network, but the recurrent interactions that implement evidence integration are segregated to a third population. Here we describe the corresponding reduced model.

As for the two-population network, the two sensory input vectors are generated independently 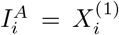, 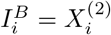, irrespective of the population structure. The pair of recurrent connectivity vectors is structured in three sub-populations. For *i* in population 1:

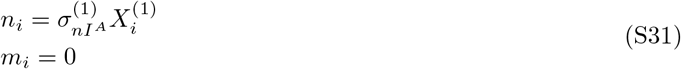

for *i* in population 2:

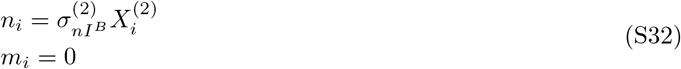

and for *i* in population 3:

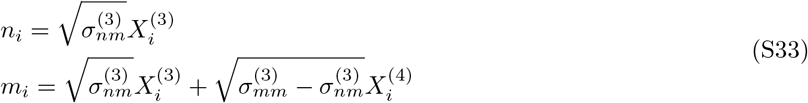

for *i* = 1, … , *N*, with 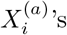 drawn independently from centered Gaussian distributions of unit variance. The readout vector reads only from the third population:

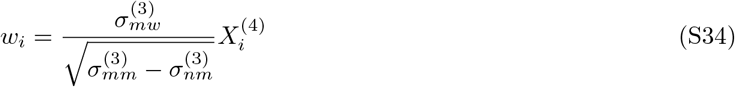

The contextual inputs are the same as in the two-population network. The overall expression for the time evolution of the internal collective variable is unchanged compared to the two populations solution Eq. (S26). Each of the effective couplings between *κ* and inputs is supported by one of two populations

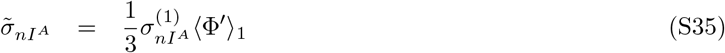

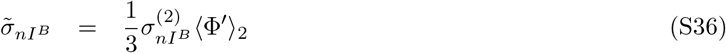

and the self-coupling of the internal collective variable is supported by the third population

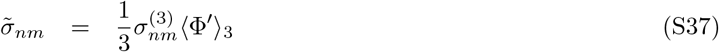

with averaged gains given by equations (S10) and standard deviations of currents onto each population by

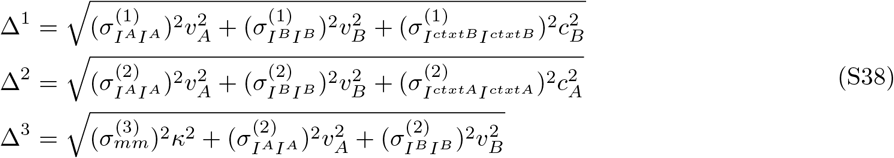

In this three-population implementation, the contextual inputs do not control the gains of neurons in the third population and thus modulate only the effective couplings that mediate the influence of sensory inputs. The non-zero covariances for an implementation of this solution are given by 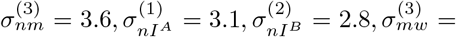 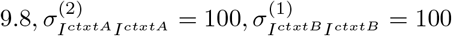.

#### S2.4 Delay-match-to-sample network

Networks trained on this task were of rank two and consisted of two populations. Here we propose a minimally parametrized network (Supp. Fig. S8a) that, similarly to the trained network presented in the main text, relies on the ability of inputs to control the autonomous dynamics of the network. The pairs of recurrent connectivity vectors defined on the first population are coupled to each other through covariances 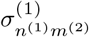 and 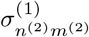 [Mastrogiuseppe and Ostojic, 2018; Beiran et al., 2021]:

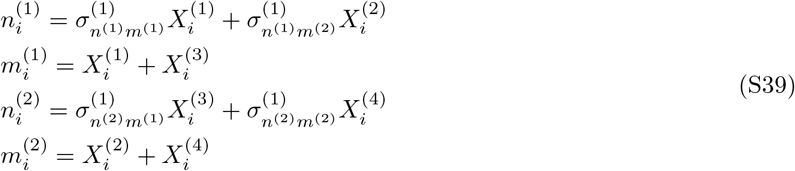

with covariances chosen such that the trivial fixed-points ***x*** = **0** is an unstable spiral point, and the dynamics defined by the first sub-population generate a limit cycle. As shown by a linear stability analysis of the dynamical equation for internal collective variables, this dynamical feature arises when the covariances are such that the following matrix has complex eigenvalues with positive real-parts [Mastrogiuseppe and Ostojic, 2018; Beiran et al., 2021]

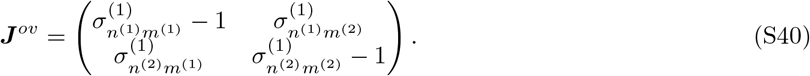

This first population is coupled to a second population which, in the absence of inputs, cancels the rotational dynamics, through the relationships 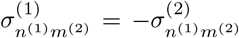 and 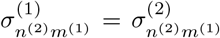. The second population also implements a pair of fixed-points that will be used to store the identity of the first stimulus throughout the delay and report the match/non-match decision. The connectivity sub-vectors on the second population can then be written as:

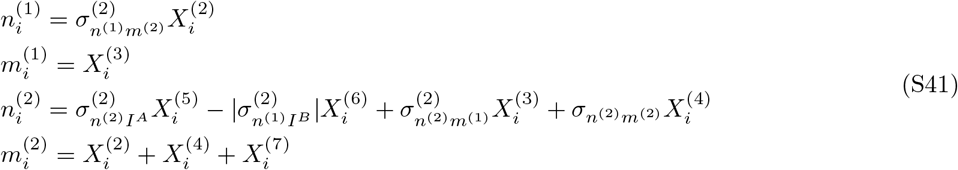

The readout vector reads only from the second population:

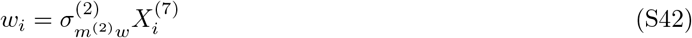

The input vector ***I***^*B*^ also stimulates only the second population, pushing the dynamics towards one fixed point on the direction ***m***^(2)^

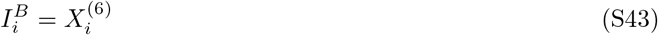

while the input vector ***I***^*A*^ activates the two populations. For units in the second population

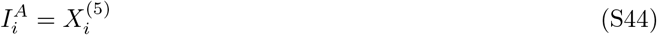

pushing the dynamics towards the other fixed point on the direction ***m***^(2)^, while for *i* in the first population

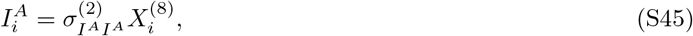

with 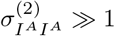

Such a connectivity leads to the dynamical equation for the two internal collective variables

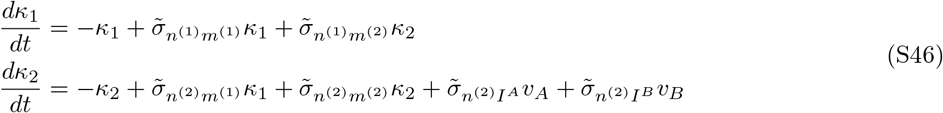

with the effective couplings mediating inputs of the form

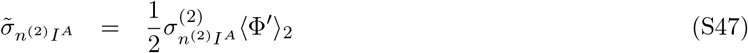

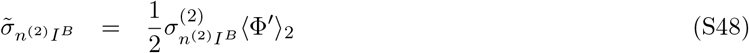

and effective couplings governing the autonomous dynamics:

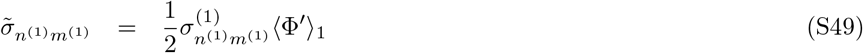

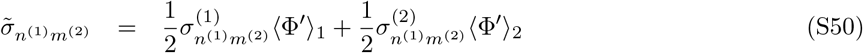

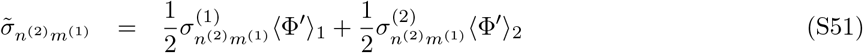

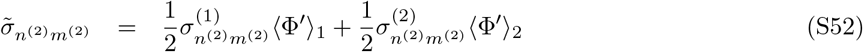

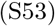

with averaged gains given by equations (S10) and standard deviations of currents onto each population

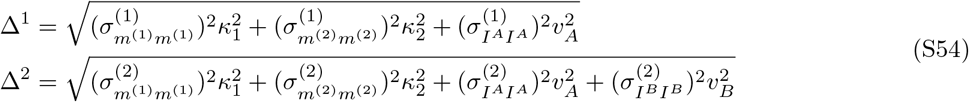

This dynamics can be graphically summarized as in Fig. 4g. It reproduces the dynamics of trained rank two networks presented in the main text (Supp. Fig. S8b), by relying on the same network mechanism, with input *A* controlling the gains of neurons in population one (Supp. Fig. S8c, middle) and thus shaping the dynamical landscape on which internal collective variables evolve (Supp. Fig. S8d). The important non-zero covariances of the reduced model are given by: 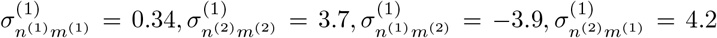 for the first population and 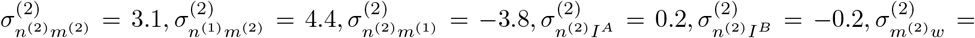 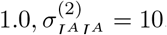.

### S3 Non-uniqueness of network implementation for a given task

We observed that varying training parameters on a given task can lead to various network implementations. We identified three factors that contribute to such variability.

A first factor is the determination of the network parameters that are trained (e.g. number of pairs of recurrent connectivity vectors *R*, whether input vectors are trained or not, scaling of trained parameters with network size, etc.). An example of this is provided by training a rank-one network on the context-dependent decision-making task, without training any of the input vectors (while the contextual-input ***I***^***ctxA***^ and ***I***^***ctxB***^ vectors are trained for the rank-one networks presented in the main text). Supp. Fig. S9 reports the analysis of such a trained network, showing that training leads to a network with three functional populations, whose implication in the computations are reproduced and detailed in a reduced model (section S2.3), and which is reminiscent of the one found in [Yang et al., 2019]. Another such example concerns the number of pairs of recurrent connectivity vectors allowed during training. For instance if training a rank-two networks on the perceptual decision-making task, one could exhibit networks with a ring-like slow manifold [Mastrogiuseppe and Ostojic, 2018], which gives rise to a single, non-linear collective variable embedded in a two-dimensional subspace.

A second factor is task parametrization. For instance we observed that training on the parametric-working memory task with fixed delays between the two stimuli, while they are drawn randomly here, leads to solutions that exploit network oscillations with periods fine-tuned to the delay, rather than a line attractor (not shown). Another such example can be put forward for the context-dependent decision-making task. Here we trained networks on a two-alternatives forced choice version of this task in which every stimulus requires one out of two responses (section 4.3.3) and found that multiple populations were required for the implementation (Supp. Fig. S11). In a Go-Nogo setting, where the alternatives are to either respond or not, flexible input-output associations can be implemented with a single population, through a mechanism based on biasing the response threshold rather than modulating the gain [Mastrogiuseppe and Ostojic, 2018].

A third factor is the stochastic nature of the training procedure, with initial connectivity being randomly drawn for each training, as well as the stochastic split of training examples into batches inherent to stochastic-gradient-descent-based methods used here. In Supp. Fig. S14, we show the dynamics of a network trained on the delay-match-to-sample task obtained for the same task parametrization and the same trained parameters. Similarly to the solution described in the main text, it relies on gain modulations through external inputs to shape the dynamics of the network in the *κ*_1_ − *κ*_2_ plane. However, that solution relies on four stable fixed points in the autonomous dynamics, two of them encoding the memory of the first stimulus (A/B) while the 2 other encode the final decision (match/non-match). The mechanism relies on different input-driven dynamics, each implementing bistable dynamics with a separatrix that moves just enough to execute a XOR operation during the second stimulation.

**Figure S5.**
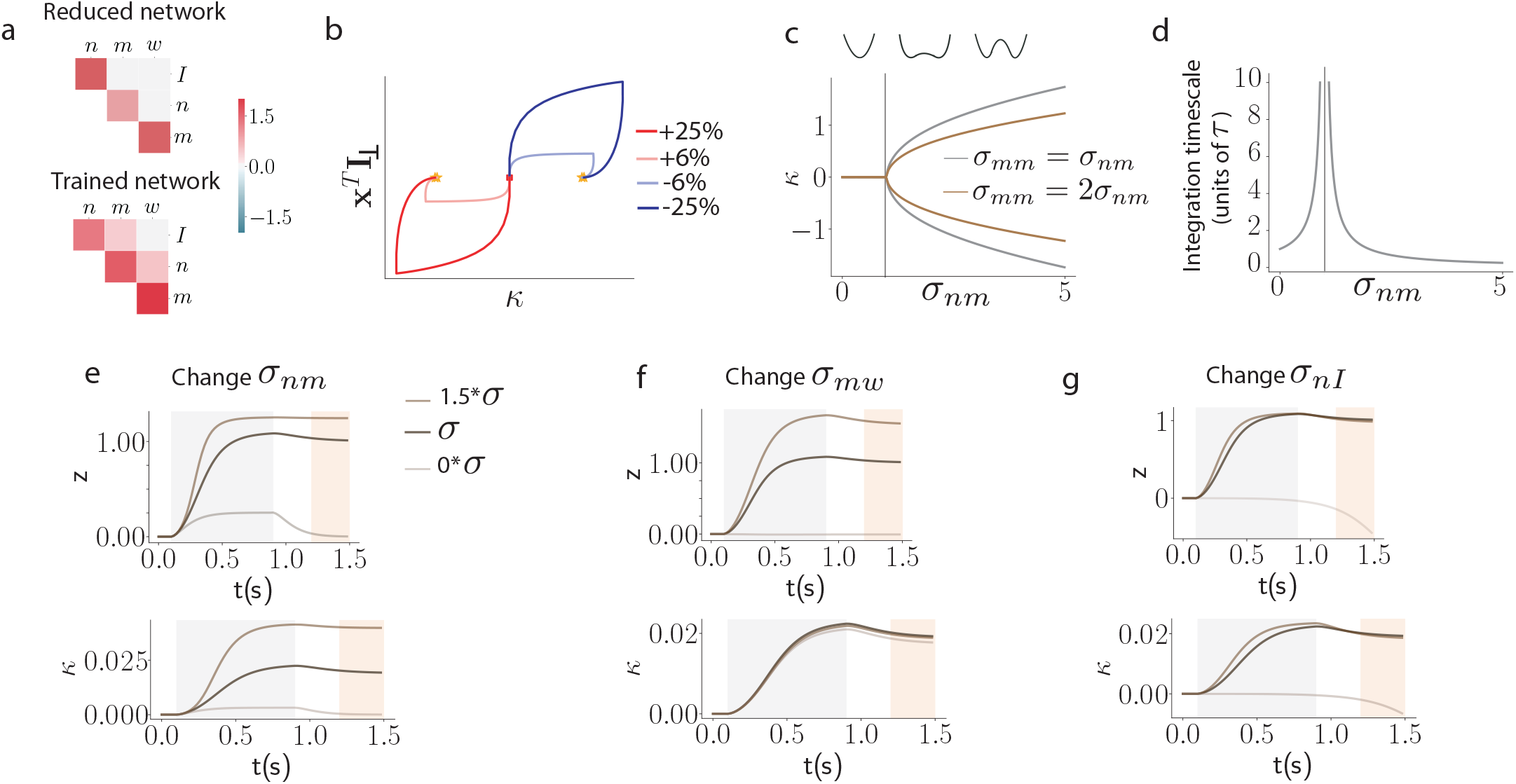
Theoretical analysis of reduced models for the perceptual DM task. (a) Covariances between connectivity vectors of reduced and trained networks. (b) Neural trajectories are embedded in the two dimensional subspace spanned by vectors ***m*** and ***I***, such that neural activity is fully characterized by the two projections ***x***^*T*^ ***I*** and ***κ*** = ***x***^*T*^ ***m***. Lines of different colors stand for different values of the input to the network. (c) Bifurcation analysis of the autonomous dynamics showing the value of the internal collective variable *κ*^*^ at the stable fixed-points of the network. Insets represent the shape of a potential *V* (*κ*) from which dynamics are derived (such that 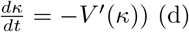 Time-scale of network dynamics around the network state at trial start ***x*** = **0** for *σ*_*mm*_ = *σ*_*mn*_. (e,f,g) Changes in readout (top) and internal collective variable (bottom) dynamics as features of the network connectivity are varied at 0 and 1.5 times their original value (see section S2.1 for details of connectivity parameters).

**Figure S6.**
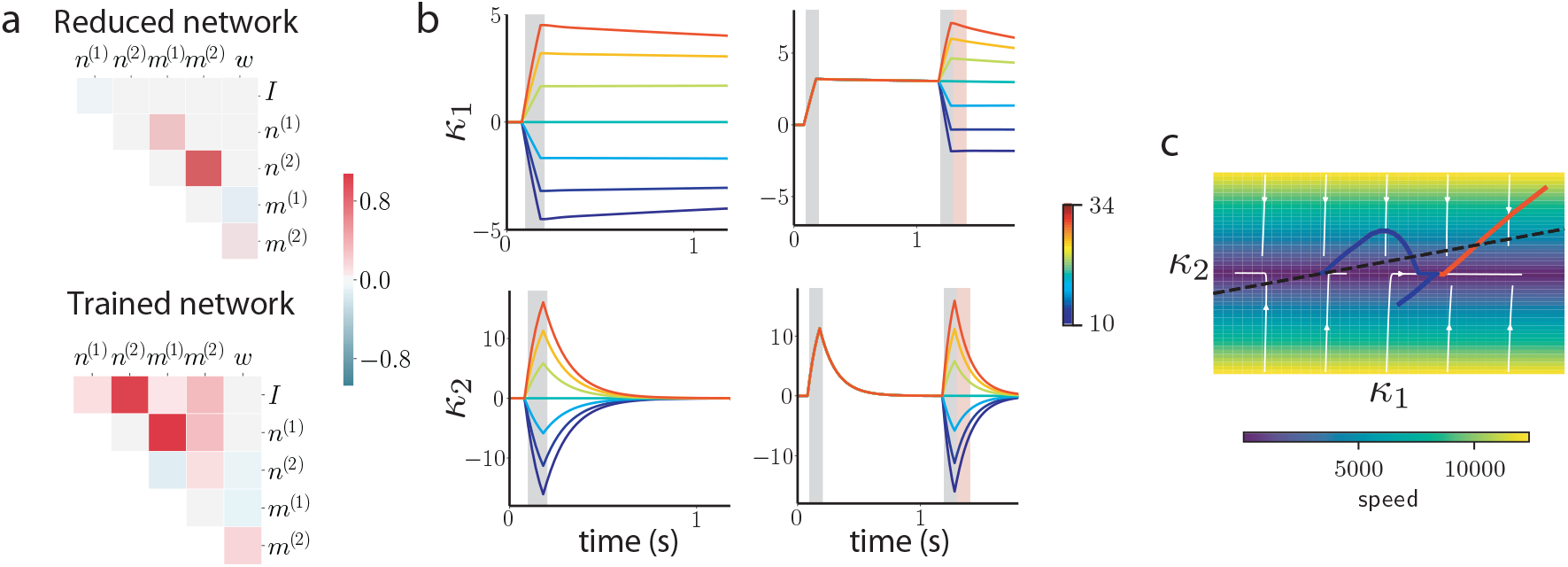
Theoretical analysis of reduced models for the parametric working memory task. (a) Covariances between connectivity vectors of reduced and trained networks. (b) Low-dimensional dynamics of internal collective variables. Left: responses to the first stimulus (colors represent different values of *f*_1_). Right: responses throughout the whole trial to a range of values for the second stimulation (*f*_1_ fixed at 30Hz, colors represent different values of *f*_2_). (c) Dynamical landscape on which the two internal collective variables evolve. From yellow to blue color, decreasing norm of the flow field 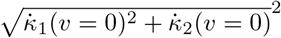 (arbitrary units). Full lines depict two trajectories corresponding to *f*_1_ = 22Hz for both and *f*_2_ = 30Hz (blue) and *f*_2_ = 14Hz (orange). The dashed line represents the direction of the readout vector ***w***.

**Figure S7.**
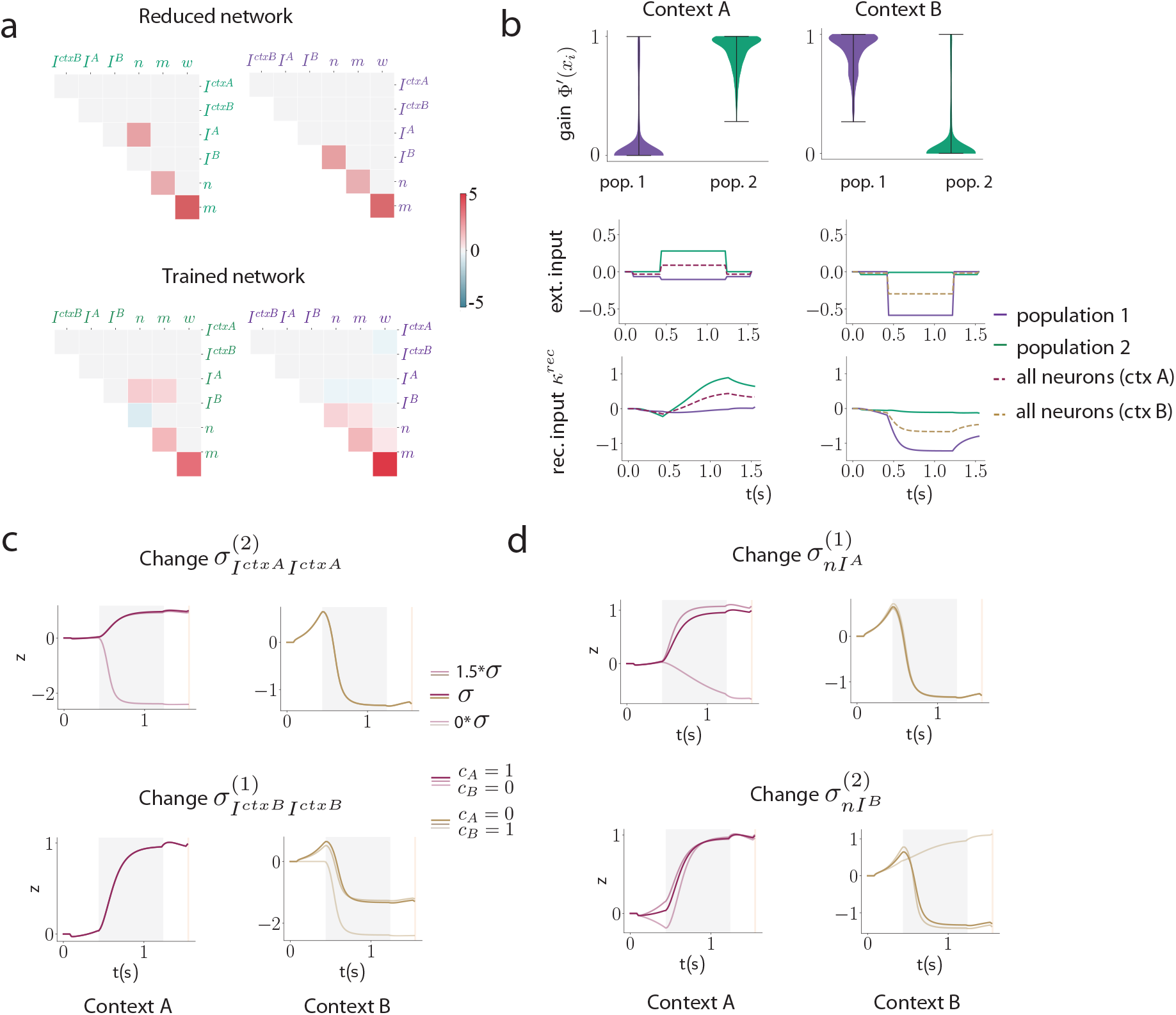
Theoretical analysis of reduced models for the context-dependent task. (a) Covariances between connectivity vectors of reduced and trained networks. (b) Top row: Distribution of single neuron gains across the two populations, in the two contexts. Middle: contributions of each population to the effective inputs to the internal collective variable (defined as in Figure 4d). Bottom: contribution to the recurrent feedback on the internal collective variable. (c,d) Changes in readout dynamics as network connectivity is varied.

**Figure S8.**
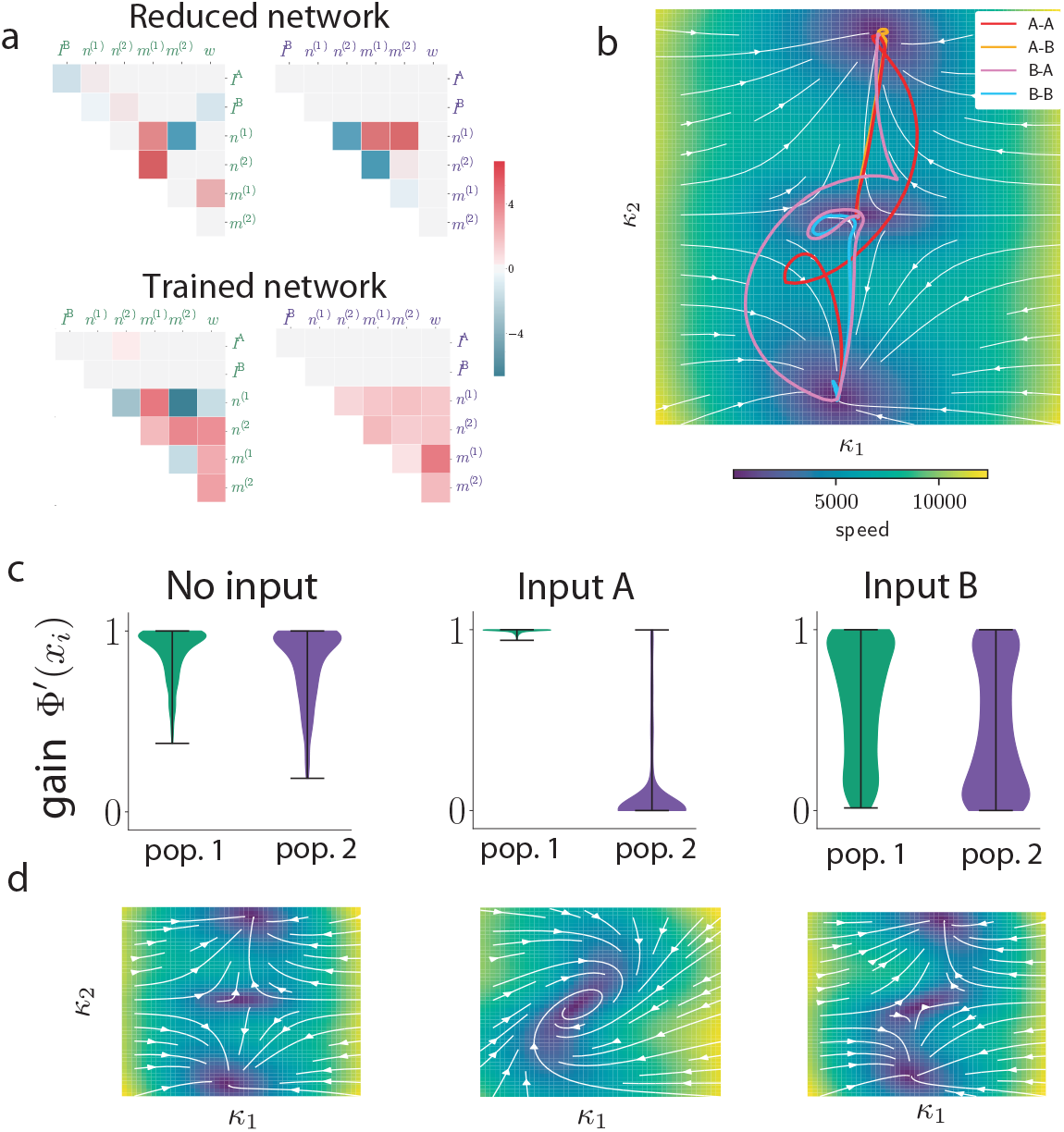
Theoretical analysis of reduced models for the delay-match-to-sample task. (a) Covariances between vectors of reduced and trained networks. (b) Trajectories of activities in the 2-dimensional space spanned by internal collective variables. (c) Distributions of individual neuronal gains in each of the two populations in the present of inputs. (d) Dynamical landscape in which the internal collective variables evolve in the various stimulation conditions of the task.

**Figure S9.**
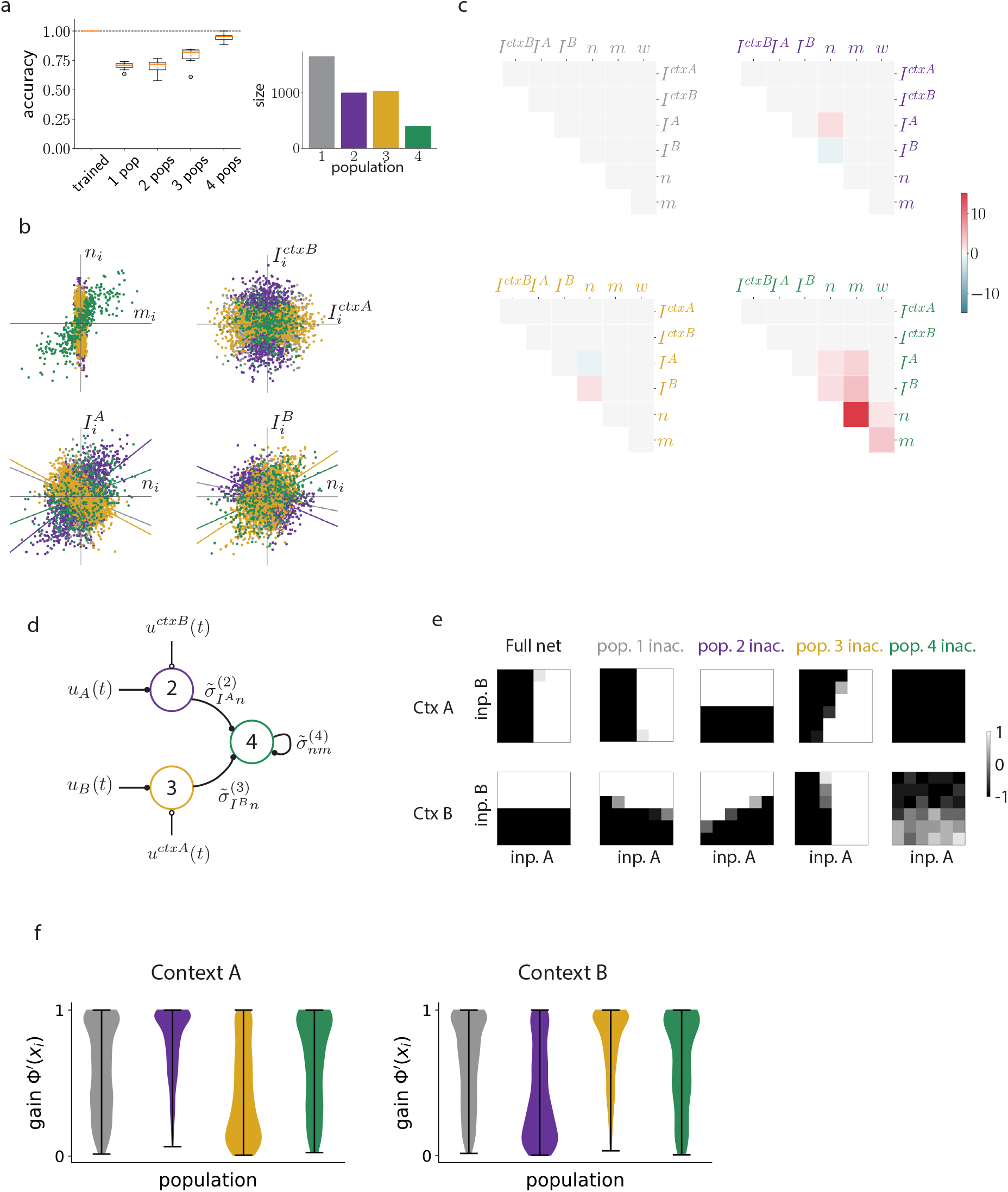
Alternative implementation of the CDM task. A network trained with different hyperparameters offers an example of an alternative solution for the CDM task, using 3 effective population and a fourth one accounting for neurons that are not involved in the task. (a) We found that 4 populations were sufficient to explain the computational mechanism used by the trained network, in the sense that resampling networks from a mixture-of-Gaussians fit to the connectivity space gives functioning networks (median accuracy of 95%), which is not the case with a lower number of populations. Note that even though the mechanism uses only 3 effective populations, the clustering and resampling procedure has to take into account a fourth one (population 1, in grey), which constitutes most of the network but is not effectively used in the task (panel e). (b) These populations all have zero mean in the connectivity space but different covariance structures as shown on these 2d projections of the full 7-dimensional connectivity space. In particular, population 4 (in green) is characterized by strong entries on the ***m*** vector and a positive covariance between its ***m*** and ***n*** entries (see (c)), showing that it can perform the effective evidence integration. Population 2 (in purple) presents strong entries on the ***I***^*ctxB*^ vector along with a positive covariance between its ***n*** and ***I***^*A*^ entries, showing it can transmit the entry signal *u*_*A*_ to the integratory feedback loop driven by the ***n***-***m*** loop and supported by population 4, unless it is driven to a low-gain regime by the strong entries on ***I***^*ctxB*^. Note that the effective couplings between input and recurrence vectors that drive the computation have to be computed at the level of the whole network (following equation (7)), even though these couplings might be supported by covariance structures in only one sub-population. In a complementary manner, population 3 (in orange) has strong entries on the ***I***^*ctxA*^ vector and a positive overlap between its ***I***^*B*^ and ***n*** entries. Finally, population 1 (in grey) does not present any obvious structure. (c) Upper-right triangle of the empirical covariance matrices for each of the four populations, showing features explained previously. (d) Illustration of the mechanism used by the network in terms of its sub-populations and couplings between variables. (e) Inactivation experiments confirm the role of each population. Here we plot psychometric response matrices similarly to what is shown in figure 3. Inactivating population 1 entirely results in little harm to the performance of the network. Inactivating population 2 results in a complete loss of performance in context A, and an unchanged performance in context B, which confirms its role in transmitting evidence for input A. Conversely, inactivating population 3 results in a loss of performance in context B and not in context A, confirming its role in transmitting evidence for input B. (f) The gain of populations 2 and 3 are differentially modulated by the context as shown in these density histograms representing the gain of each neuron in the population in each context.

**Figure S10.**
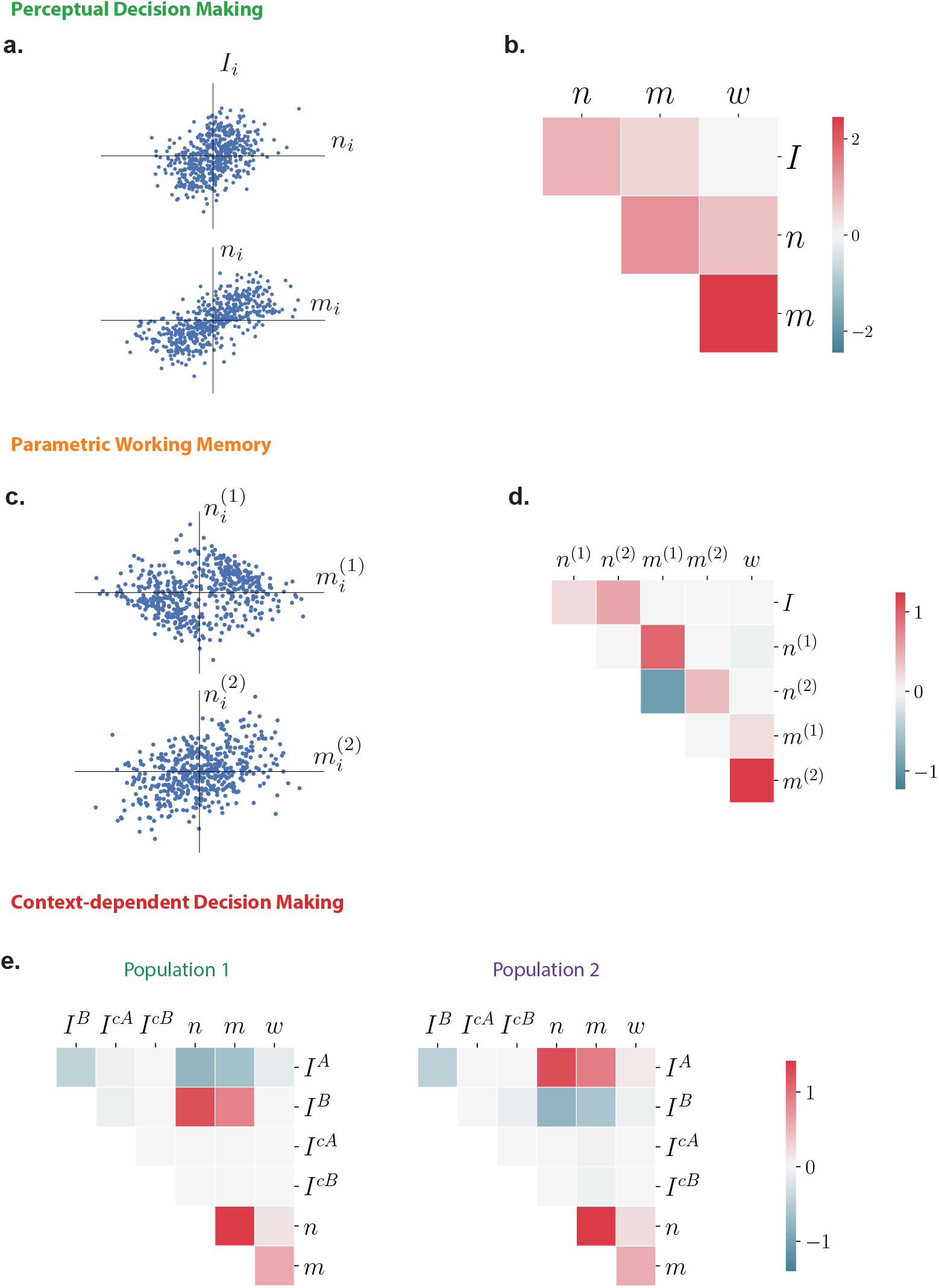
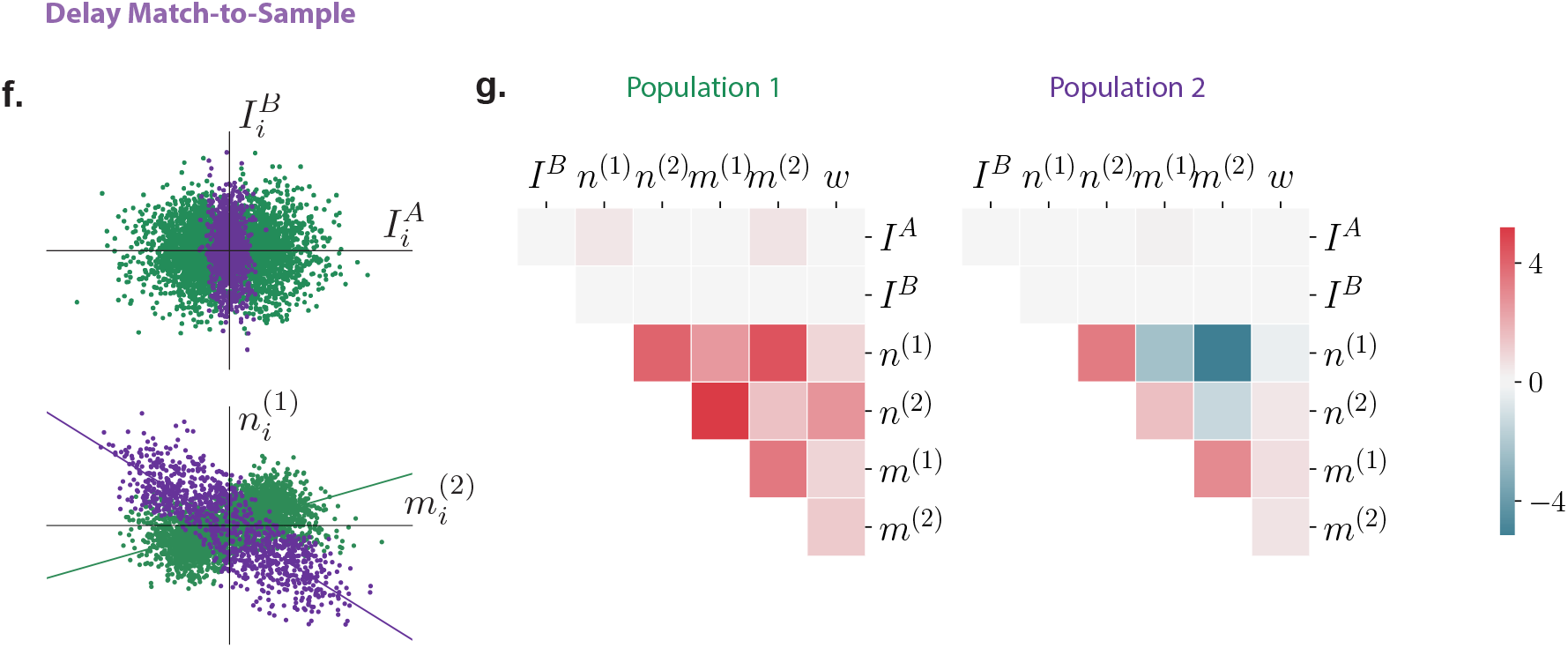
Statistics of connectivity in trained networks. (a) Two 2d projections of the connectivity space for the network trained on the DM task, showing the positive covariances between parameters *I*_*i*_ and *n*_*i*_, as well as between *n*_*i*_ and *m*_*i*_, which translate into positive overlaps between the corresponding vectors in neural state-space. (b) Upper-right corner of the covariance matrix between connectivity parameters, showing the same positive correlations. (c) Two slices of the connectivity space for the network trained on the WM task. Note that this network has been trained with different hyperparameters than those used in figure 1, and been chosen for a particularly simple usage of its two latent variables. This solution however exhibits significant clusteriness contrarily to the networks used for figure 1, and is an example of a situation where clusters can appear during training without being computationally relevant (see Figure S4). (d) Corresponding upper-right corner of the covariance matrix, showing the strong positive overlap between vectors ***n***^(1)^ and ***m***^(1)^ which drive the line attractor along direction ***m***^(1)^. The transient encoding onto direction ***m***^(2)^ is obtained thanks to a much lower overlap between vectors ***n***^(2)^ and ***m***^(2)^. (e) Upper-right corners of the different covariance matrices corresponding to each of the two populations identified on the network trained on the CDM task and used in Figures 3 and 4. Note in particular how input signal vectors ***I***^*A*^ and ***I***^*B*^ have opposite correlations with vector ***n*** depending on the population, and how contextual input vectors ***I***^*ctxA*^ and ***I***^*ctxB*^ don’t overlap with the recurrent connectivity vectors, reflecting their role as pure gain-modulators. (f) Two slices of the connectivity space for the network trained on the DMS task used in Figures 3 and 4, showing the difference in variance along the parameter 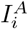 between both populations, and the opposite overlaps they induce between recurrent connectivity vectors ***n***^(1)^ and ***m***^(2)^. (g) Corresponding upper-right corners of the covariance matrices for each population.

**Figure S11.**
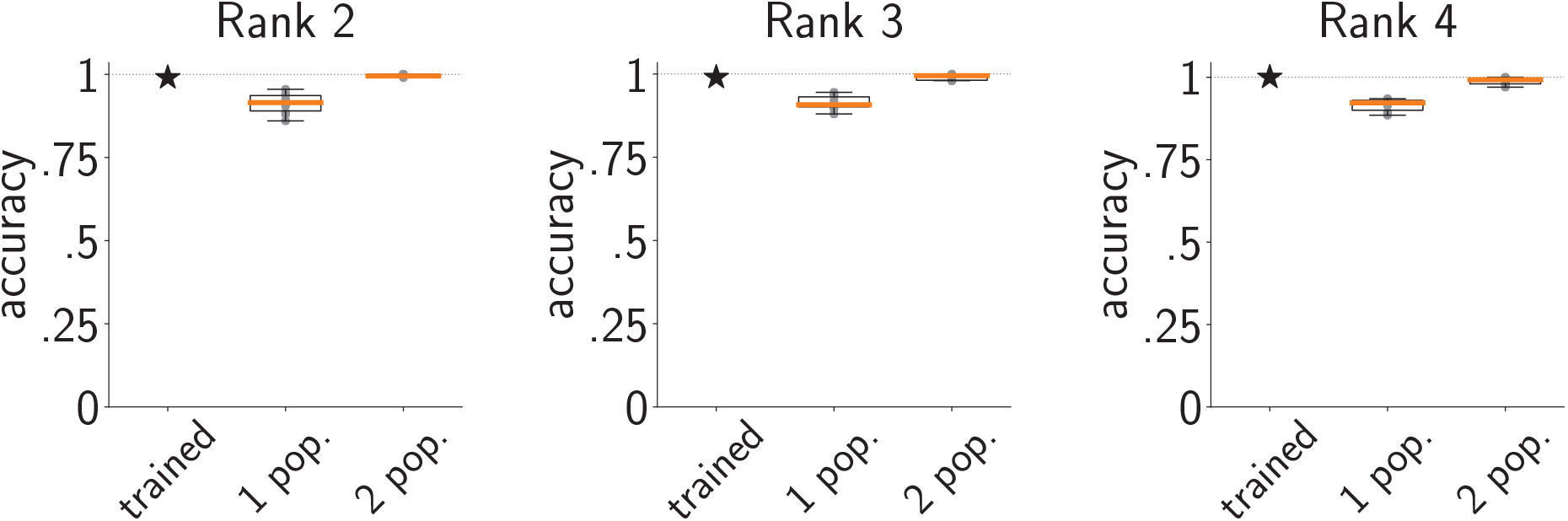
Increasing the rank maintains the requirement for population structure. For this figure we have trained low-rank networks with a rank higher than 1 on the CDM task, fitted a single Gaussian or a mixture of 2 Gaussians to the obtained connectivity space, and retrained the obtained distribution (Methods 4.7) to obtain resampled networks with a performance as high as possible. Even with this additional layer of retraining of the fitted distributions (which is only present in the main text for the DMS task) the obtained single-population networks fell short of performing the CDM task with a good accuracy. Here, 10 draws of a single network for each combination of rank and number of populations are shown (orange line : median, box: 1st and 3rd quartiles, whiskers: min and max). Note that the obtained performance seems high, at around 90% for all ranks for the single-population networks, but hides a heavy overfitting that networks are able to do with a higher-dimensional space.

**Figure S12.**
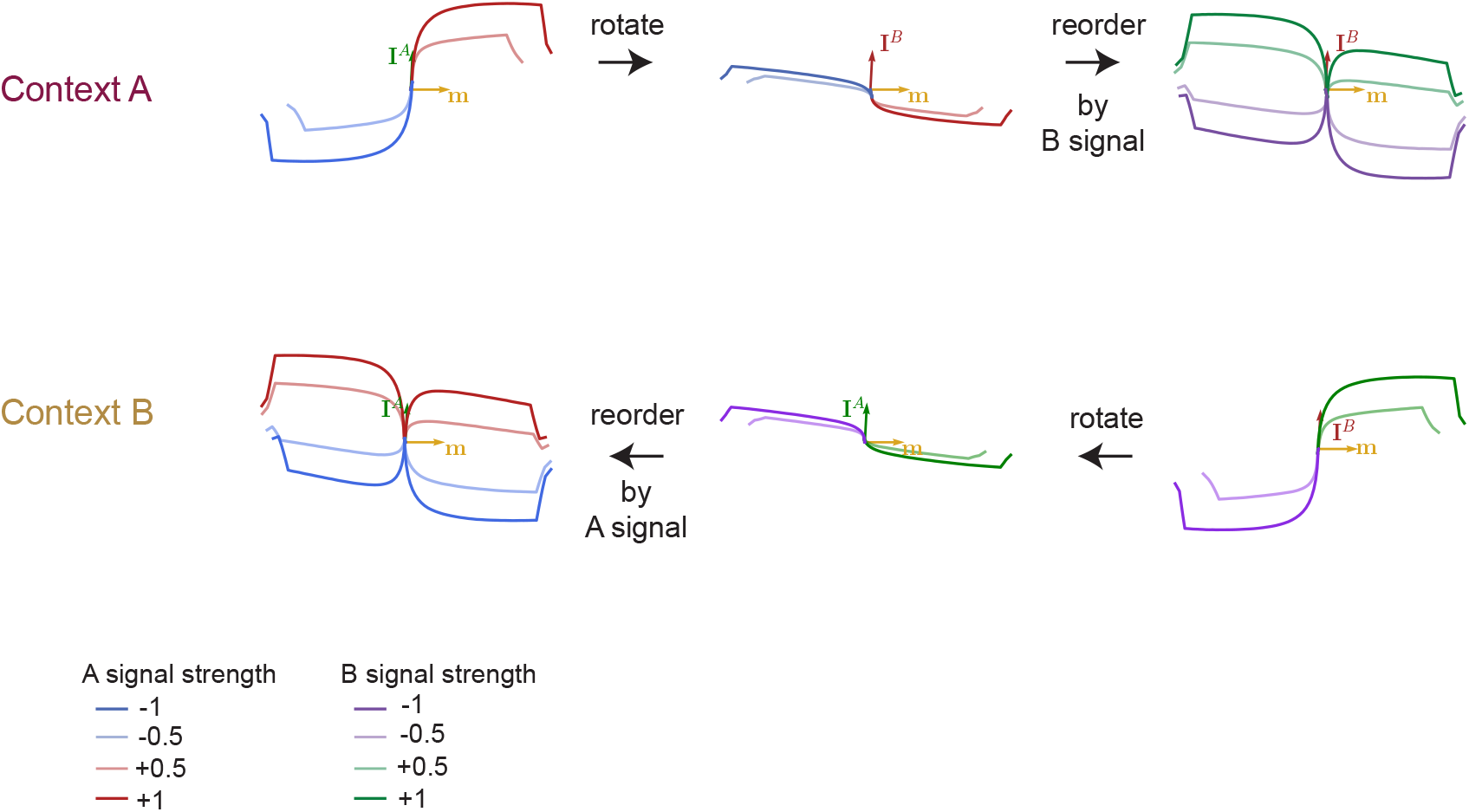
Context-dependent decision making state-space dynamics. Here we reproduce figures akin to those presented in [Mante et al., 2013] for the trained low-rank network used in figures 3 and 4. We generate 32 conditions corresponding to different combinations of context, signal A coherence and signal B coherence and then project condition-averaged trajectories either on the plane spanned by the recurrent connectivity vector ***m*** (which corresponds to the choice axis) and the input vector ***I***^*A*^, or on the ***m − I***^*B*^ plane. Similarly to what was observed in [Mante et al., 2013], signal A strength is encoded along the ***I***^*A*^ axis, even when it is irrelevant (lower left corner), and signal B strength is encoded along the ***I***^*B*^ axis, even when it is irrelevant (top right corner).

**Figure S13.**
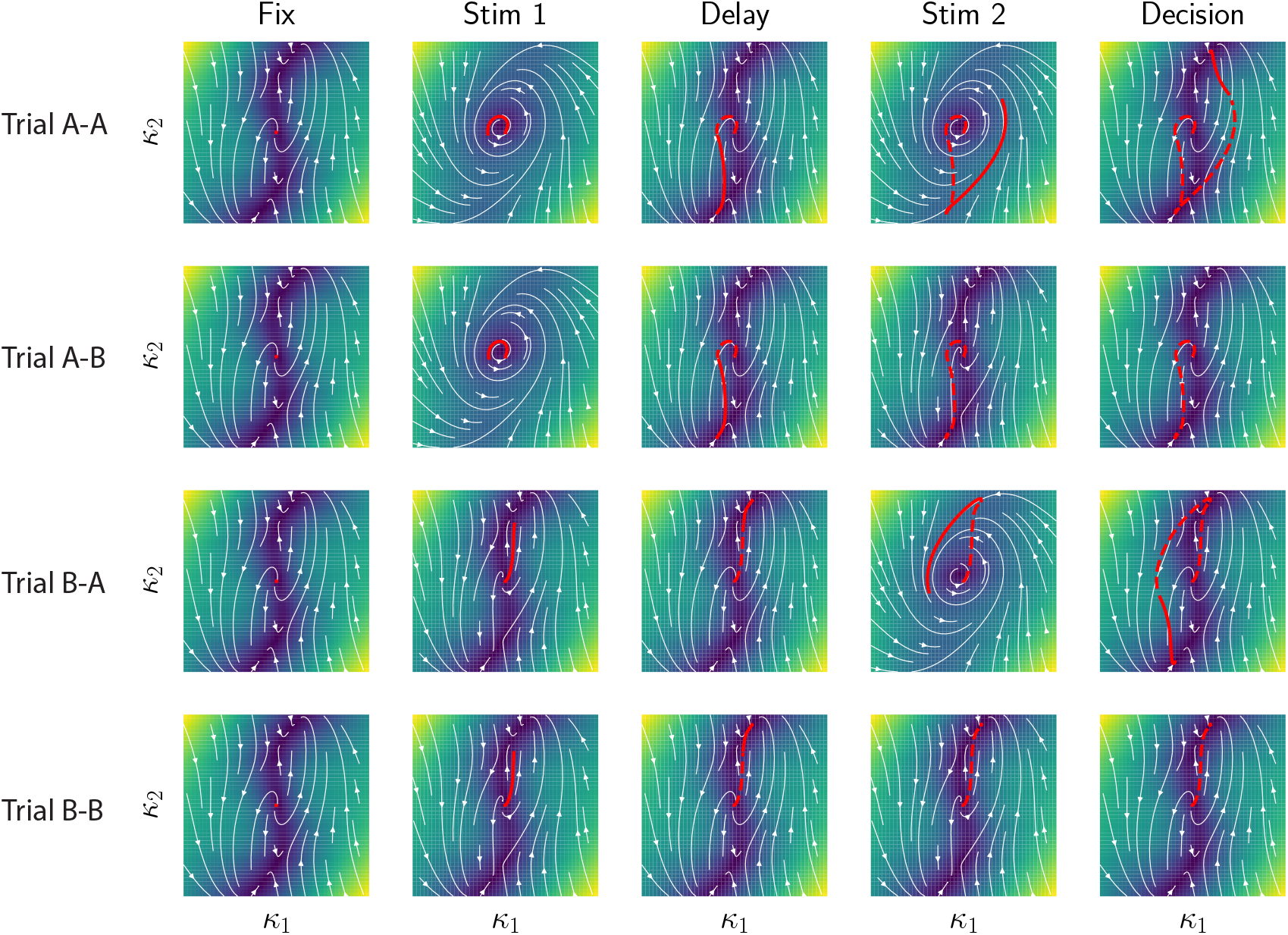
Unrolled dynamics in the DMS network. This panel presents the trial-averaged state-space trajectories of the network presented in Figures 3 and 4, unrolled by trial epoch (for each column the filled red line corresponds to the trajectory in the current epoch, the dashed line to the previous parts of the trajectory).

**Figure S14.**
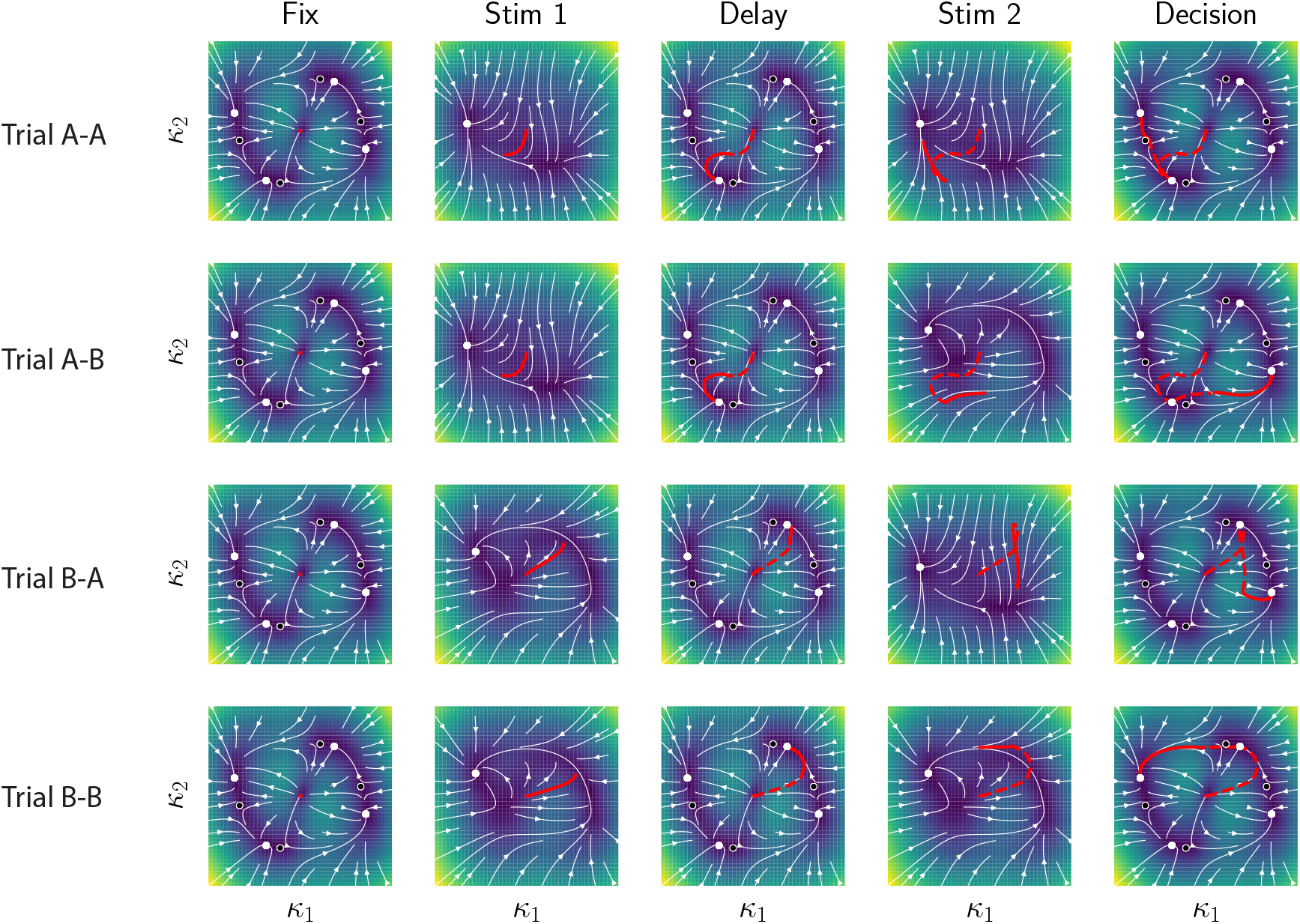
Alternative DMS solution. The network presented in Figures 3 and 4 of the main text is a particular implementation of the task, but different solutions were found as we retrained networks from different initial random connectivity, even as all hyperparameters were kept constant. Here, we present dynamics for another rank-2 implementation of the DMS task which has the particularity of assigning symmetric roles to A and B inputs. It is based on four stable fixed points in its autonomous dynamical landscape, two of those (top and bottom) used for encoding the memory of the first stimulus, and the two other (left and right) used for encoding the final match/non-match decision.

**Figure S15.**
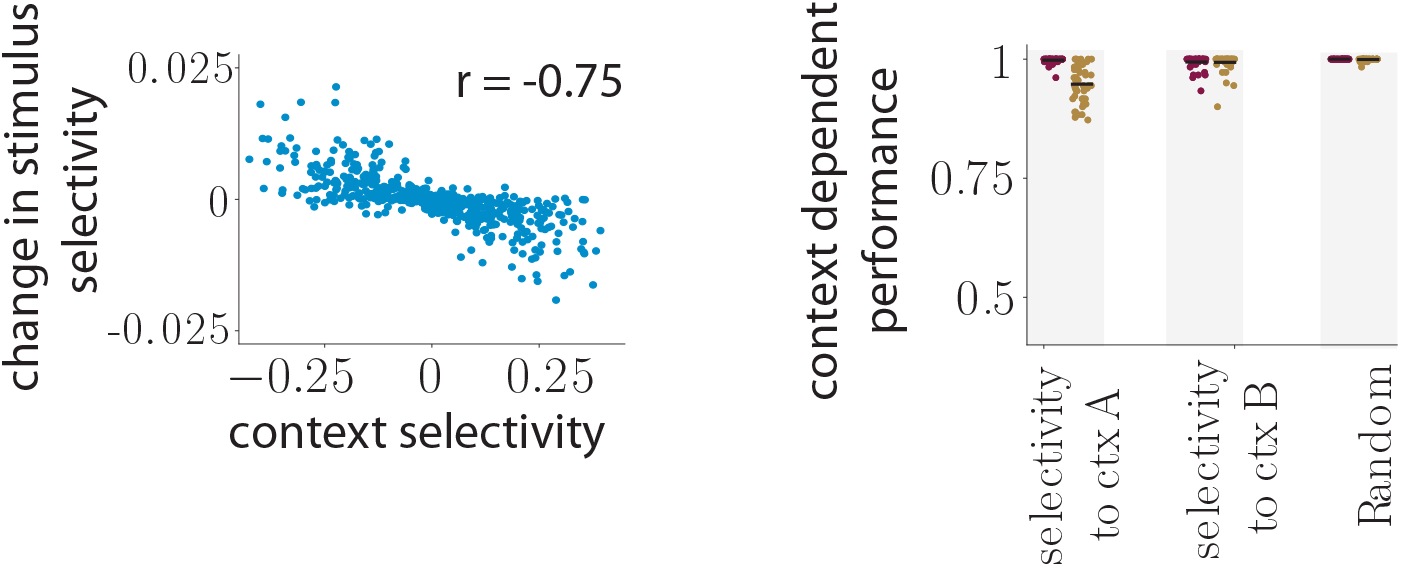
Control the strength of context cues in the MDM task. Here the context input vectors have been multiplied by a factor five compared to the network analyzed in Fig. 5g. (a) Context cues are thus able to set the functioning point of some neurons closer to the saturating part of the transfer function, leading to the observation of non-linear mixed-selectivity between context and changes in sensory representation with context. (b) As opposed to the CDM task, this particular feature of selectivity is not functional as revealed by specifically inactivating neurons with a high selectivity to context A or B, showing a similar decrease in behavioral performance as for randomly selected neurons.

